# ECM remodeling and spatial cell cycle coordination determine tissue growth kinetics

**DOI:** 10.1101/2020.11.10.376129

**Authors:** Anna P. Ainslie, John Robert Davis, John J. Williamson, Ana Ferreira, Alejandro Torres-Sánchez, Andreas Hoppe, Federica Mangione, Matthew B. Smith, Enrique Martin-Blanco, Guillaume Salbreux, Nicolas Tapon

## Abstract

During development, multicellular organisms undergo stereotypical patterns of tissue growth to yield organs of highly reproducible sizes and shapes. How this process is orchestrated remains unclear. Analysis of the temporal dynamics of tissue growth in the *Drosophila* abdomen reveals that cell cycle times are spatially correlated and that growth termination occurs through the rapid emergence of a population of arrested cells rather than a gradual slowing down of cell cycle time. Reduction in apical tension associated with tissue crowding has been proposed as a developmental growth termination mechanism. Surprisingly, we find that growth arrest in the abdomen occurs while apical tension increases, showing that in this tissue a reduction in tension does not underlie the mechanism of growth arrest. However, remodeling of the extracellular matrix is necessary for tissue expansion. Thus, changes in the tissue microenvironment, and a rapid exit from proliferation, control the formation of the adult *Drosophila* abdomen.

## Introduction

During development, patterns of growth must be tightly coordinated in time and space. This is dictated by chemical signals (hormones, morphogens, nutrients) and the physical environment (Boulan et al., 2015; Irvine and Shraiman, 2017; Penzo-Mendez and Stanger, 2015). However, due to the difficulty of measuring quantitative parameters of tissue growth in living organisms, we lack an integrated understanding of growth arrest *in vivo*. This has limited our ability to analyze developing systems *in vivo* with the same level of precision as expanding populations of unicellular organisms or animal cells in culture (Loeffler and Schroeder, 2019; Sauls et al., 2016).

Nevertheless, studies of the *Drosophila* wing imaginal disc have yielded several models of tissue growth and size determination (Boulan et al., 2015; Gou et al., 2020; Irvine and Shraiman, 2017). Wing disc cells arrest in G2 at the larval/pupal transition, when the organ has reached approximately 30,000 cells (Martin et al., 2009). As this tissue progresses through its most substantial growth phase, cell proliferation gradually slows down, with a cell cycle time of around 6-10 h increasing to over 20-30 h at the larval-pupal transition (Johnston and Sanders, 2003; Mao et al., 2013; Martin et al., 2009; Milan et al., 1996; Wartlick et al., 2011b). This suggests that a mechanism triggered well in advance of the tissue reaching its target size leads to a slowdown and eventual arrest of proliferation.

Proposed models to explain the growth kinetics of the developing wing disc generally assume a role for the growth-promoting action of diffusible morphogens or spatial and temporal changes in the mechanical state of disc cells (Gou et al., 2020; Irvine and Shraiman, 2017; Wartlick et al., 2011a). Thus, spatial variations in growth rates have been suggested to lead to a build-up of tensile and compressive forces that in turn affect growth rates through mechanical feedback (Aegerter-Wilmsen et al., 2007; Hufnagel et al., 2007; Shraiman, 2005). Laser ablations have indeed suggested that apical tension diminishes as wing disc development proceeds, correlating with decreased proliferation, providing a potential mechanism for growth termination (Pan et al., 2018; Rauskolb et al., 2014). Increased mechanical tension has also been demonstrated to drive cell division in a number of experimental systems, including the mouse skin and mammalian cells in culture (Irvine and Shraiman, 2017; LeGoff and Lecuit, 2015). Thus, an attractive mechanism to account for developmental size control is one by which mechanical tension promotes growth, and as cell density increases, tension decreases, leading to growth termination.

The recent development of analytical tools that capture cell and tissue deformations (Etournay et al., 2016; Guirao et al., 2015), combined with progress in live imaging *in vivo* (Mavrakis et al., 2010) have transformed our ability to derive quantitative parameters that can be used to understand and model complex developmental processes. A limitation of studies in the *Drosophila* wing disc is its inaccessibility for these live imaging tools to follow individual cell behavior and mechanical properties over time in the growing tissue. Thus, we chose to analyze the growth of the *Drosophila* abdominal epidermis, where individual cells can be imaged and tracked *in vivo* during the pupal stages when the animal is sessile (Bischoff and Cseresnyes, 2009; Mangione and Martin-Blanco, 2018; Prat-Rojo et al., 2020).

The adult *Drosophila* abdominal epidermis develops from small islands (nests) of cells called histoblasts, which are specified during embryonic development (Guerra et al., 1973). Each abdominal hemisegment contains four nests, two located dorso-laterally (dorsal anterior and dorsal posterior), one located ventrally and one spiracular nest located laterally (Madhavan and Madhavan, 1980; Roseland and Schneiderman, 1979), Figure 1A). During the larval period, the histoblasts are quiescent (G2 arrest) (Bryant and Schneiderman, 1969; Mandaravally Madhavan and Schneiderman, 1977), and are induced to enter the cell cycle by the pulse of the steroid hormone ecdysone that triggers the larval/pupal transition (0 h After Puparium Formation – APF) (Ninov et al., 2007). Following a period of three cleavage divisions, the histoblasts enter an expansion phase (around 14-16 hAPF) when they proliferate and grow, displacing the surrounding Larval Epidermal Cells (LECs, large polyploid cells that formed the larval epidermis), which are extruded and engulfed by macrophages (hemocytes) patrolling the hemolymph underneath the epidermal layer (Bischoff and Cseresnyes, 2009; Michel and Dahmann, 2020; Nakajima et al., 2011; Ninov et al., 2007; Prat-Rojo et al., 2020; Teng et al., 2017) (Figure 1B). LEC death requires both ecdysone signaling and displacement by the expanding histoblast nests (Michel and Dahmann, 2020; Nakajima et al., 2011; Ninov et al., 2007; Prat-Rojo et al., 2020; Teng et al., 2017). Once the histoblasts cover the entire abdominal surface, proliferation ceases and differentiation proceeds to give rise to the adult abdomen. Here, using live imaging and cell tracking, we analyze the growth of the dorsal histoblast nests at the cellular and tissue scale to uncover how cellular behaviors give rise to tissue growth kinetics.

**Figure 1.**
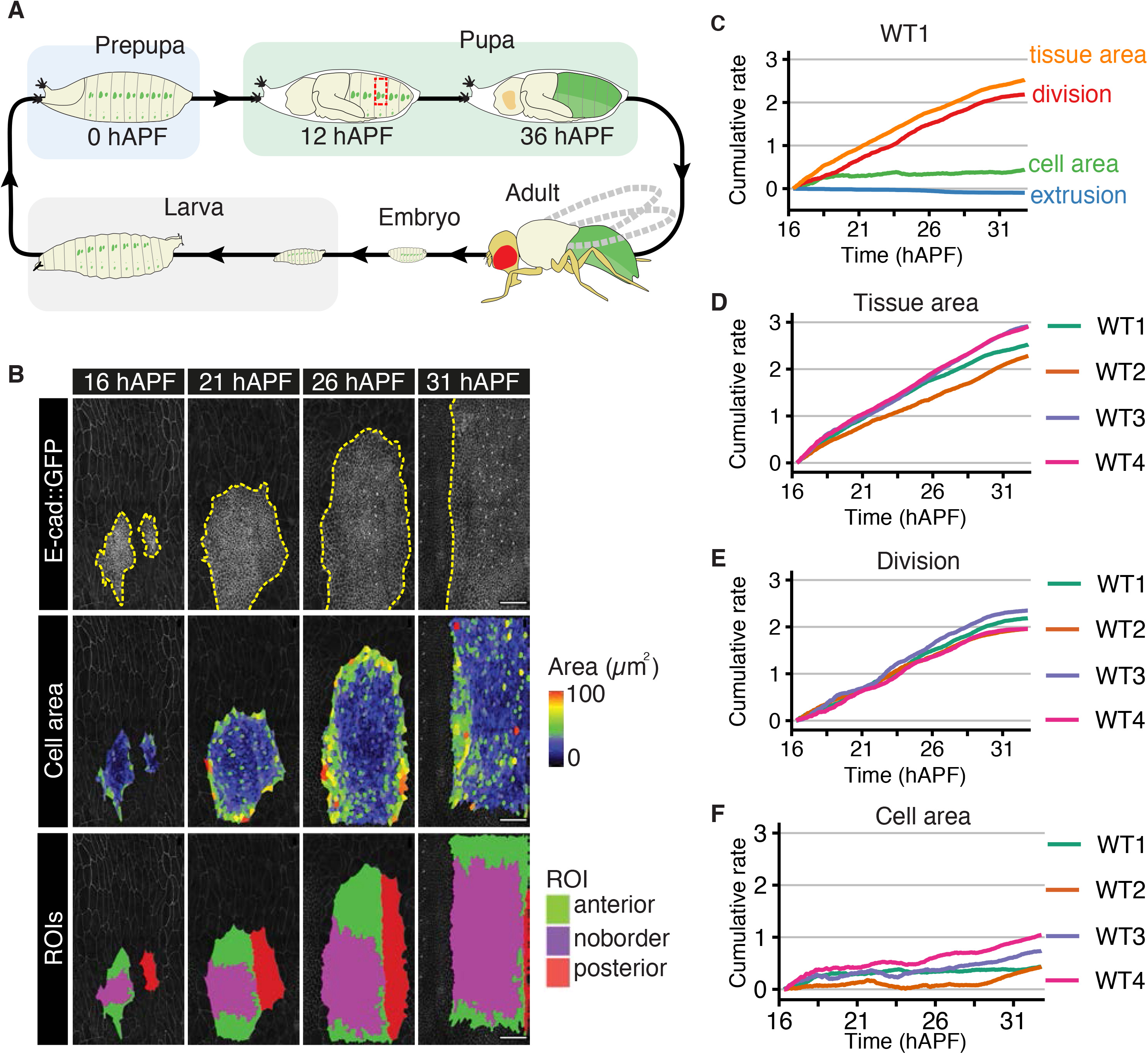
Quantitative analysis of cellular contributions to abdomen growth. (A) *Drosophila* life cycle, with histoblasts in green. The red dotted rectangle on the pupa indicates the field of view imaged in (B). Adapted from (ISBN 9780879694722). (B) Top row: Histoblasts in live pupa expressing *E-cad∷GFP* at the time points indicated. Yellow dotted line surrounds the histoblast nests. At 16 hAPF, the anterior (larger) and posterior (smaller) histoblast nests are still visible and surrounded by the LECs. The nests then spread and eventually occupy the entire surface of the segment. Middle row: Heat map showing cell areas. Bottom row: labelled regions of interest (ROIs); Anterior (green+magenta), posterior (red) and the ‘no border’ ROI (magenta). Scale bars=50 *μ*m. Dorsal is to the top, anterior is to the left in all images, unless otherwise indicated. (C) Decomposition of the cumulative area expansion rate of the tissue into contributions from cell division, cell area change and cell extrusion (WT1, noborder ROI). (D-F) Cumulative tissue area expansion rate (D) and contributions from cell division (E) and cell area (F), for 4 different WTs. WTs are time-aligned based on the appearance of sensory organ precursor (SOP) cells (Figure S1G).

## Results

### Measuring the growth kinetics of the *Drosophila* dorsal abdomen

To derive quantitative parameters describing the growth of the *Drosophila* abdomen in wild type animals, we imaged cell junctions with E-cad∷GFP from 16 h to 31 hAPF (see (Mangione and Martin-Blanco, 2020) and STAR Methods) (Figure 1A, B). The resultant movies were processed and analyzed using a custom-built pipeline to track cells (Figure 1B, Figure S1A-E, STAR Methods, Movie 1). We then defined an anterior “no border region of interest” (noborder ROI) consisting of complete lineages of histoblasts that never come into contact with the edge of the image frame during the movie (Figure 1B, bottom row, Movie 1 and Supplementary Theory). The emergence of specialized cells called Sensory Organ Precursors (SOPs) was used to temporally align four wild-type movies (Figure S1F-K and STAR Methods).

### The dorsal histoblast nests expand through cell division

To analyze tissue growth, we performed a shear decomposition analysis of the segmented wild type movies using Tissue Miner (Etournay et al., 2016; Merkel et al., 2017). We focused our analysis on the expansion kinetics of the histoblast nests by examining how the relative rates of change in cell area and cell number, due to either proliferation or delamination contribute to the area expansion rate of the tissue (Figure 1C-F, Figure S1L-O). This quantitative analysis showed that the dorsal histoblast nests grow primarily via cell division (Figure 1C, E), as previously proposed (Ninov et al., 2007). The average cell area increase contributed modestly to tissue area expansion, and mainly in the early stages of expansion (Figure 1C, F). Cell loss (“extrusion”), through either cell death or delamination, had a minimal contribution to histoblast nest deformation (Figure 1C). The cell area contribution to growth is far more variable than that of cell divisions (Figure 1E, F). Furthermore, cell areas in the anterior nest are also spatially variable, with cells around the periphery of the nest having a larger apical area than their counterparts in the nest center (Figure 1B, middle row). This may be due to their location at the interface with LECs, which have been shown to exert forces on the boundary histoblast when undergoing apoptosis (Prat-Rojo et al., 2020; Teng et al., 2017). As cell division is the dominant factor in histoblast nest growth, we focused subsequent analysis on their proliferation dynamics.

### Histoblasts divide with a narrow distribution of cycle times, uncorrelated with cell geometry but correlated in space

When examining the contribution of cell division to tissue expansion (Figure 1E), we noticed that cell number increased in alternating periods of slower and faster rates. Indeed, plotting the cell division rate (number of cells created by division per unit time, relative to total cell number) as a function of time for different WT movies revealed an oscillatory behavior with a period of ~ 4 h (Figure 2A). About three oscillatory peaks can be observed after 16 hAPF before the cell division rate starts to decrease around 28 hAPF (Figure 2A). All four analyzed wild type movies exhibited similar dynamics in the decay of cell division rate, although the oscillation peaks were variable in time between the different animals (Figure 2A). We explored whether the average cell cycle duration was oscillating through time and found that, in contrast to the proliferation rate, the average cycle time (~4.5h±0.6, mean ±SD over 4 analyzed WTs) varied only weakly in time (Figure 2B, S2A).

**Figure 2.**
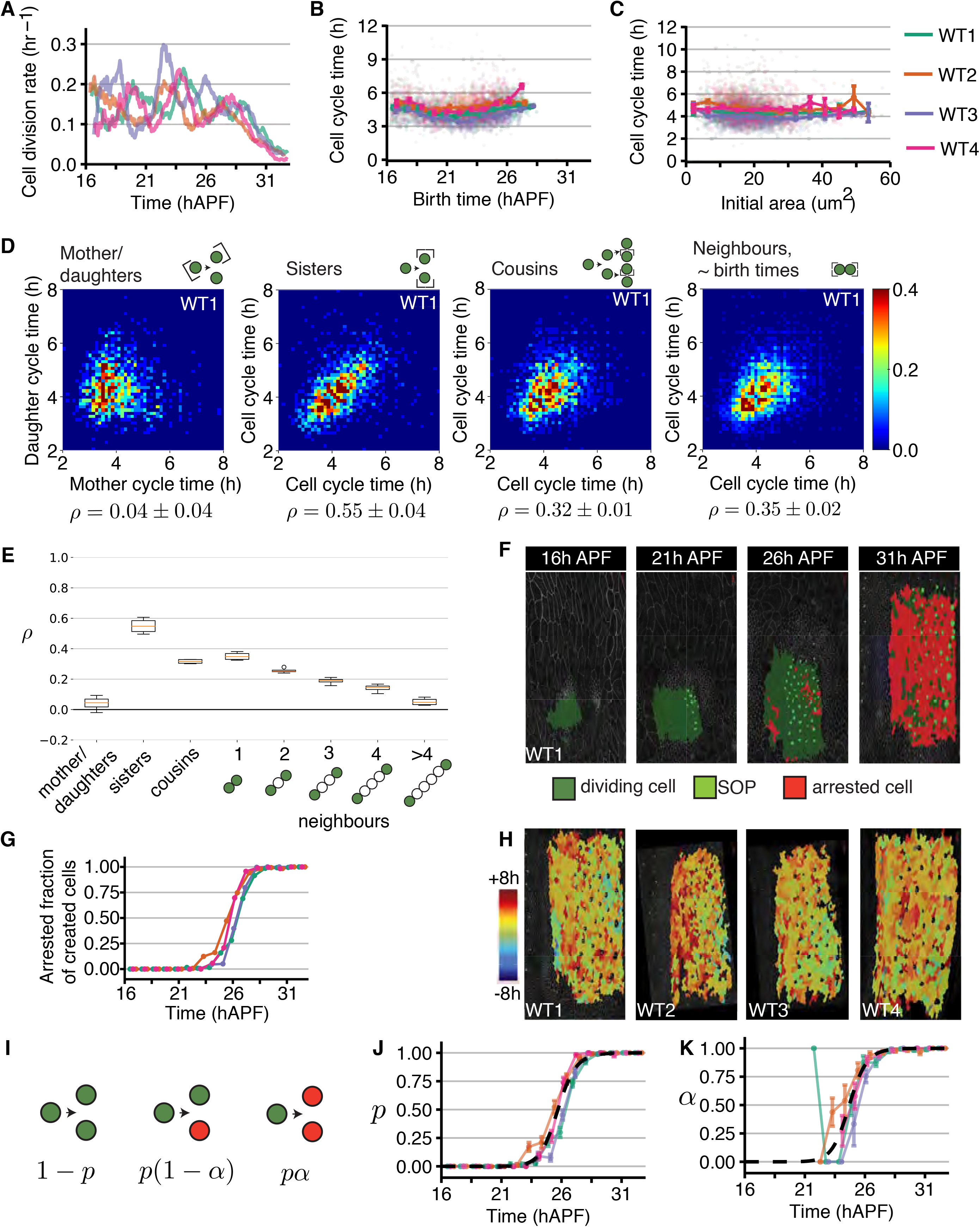
Histoblast cell cycle times and transition to proliferative arrest. (A) Cell division rate in each WT movie as a function of time (after application of moving average, with a top-hat smoothing kernel with window size 0.875 h). The cell division rate oscillates up to ~28 hAPF before decaying. Data from noborder ROIs. (B) Cell cycle time as a function of birth time for each WT movie. Data from noborder ROIs. (C) Cell cycle time as a function of initial cell apical area, for each WT movie. Larger dots: binned data. Error bars: SEM for the bin. Faint smaller dots: individual data points. Data from noborder ROIs. (D) Probability density of pairs of cell cycle times, where pairs are taken between, from left to right, mother-daughters, sisters, cousins, and nearest-neighbors whose birth time differ by less than 0.5 hours. Cells are taken from the visible anterior nests. Sister cells are excluded from neighbor correlations. *ρ*, Pearson correlation coefficient (see Supplementary Theory). Mean±SD are calculated over the values for the 4 WTs. Spearman correlation coefficients are *ρ*_s_ =0.06±0.04 (mother-daughters), *ρ*_s_ =0.62±0.03 (sisters), *ρ*_s_ =0.33±0.03 (cousins), *ρ*_s_=0.41±0.02 (nearest-neighbours born at similar time). (E) Pearson correlation coefficient *ρ* of cell cycle times for mother-daughter pairs, sisters, cousins, and increasing cell-cell distance (sister cells excluded). (F) Snapshots showing the appearance of SOPs and arrested cells within the noborder ROI (WT1 movie). (G) Fraction of created cells that are arrested as a function of time, for all WT movies (color code as in panels A-C). (H) Snapshots of the final frame for each WT movie with arrested cells colored by the time of their appearance, relative to each movie’s “switch time” for the probability of arrested cell creation (panel J). (I) Schematic defining the parameters *p* and α characterizing arrested cell creation. (J) Probability *p* that a division creates at least one arrested cell as a function of time. Each movie’s *p* curve can be fitted with its own Hill function (not shown) and the switch time of that function is referenced in (H). Black dashed line, manual fit to the average movie behavior. (K) Probability *α* that a cell division gives rise to two arrested cells, conditioned on the cell division giving rise to at least one arrested cell. Blacked dashed line, as in (J).

Work in cell culture and several *in vivo* systems has suggested that geometric constraints influence proliferation, for instance cells with a larger surface area are generally more likely to divide than cells with a constrained area (Irvine and Shraiman, 2017; LeGoff and Lecuit, 2015; Lopez-Gay et al., 2020). We therefore wondered if there was a relationship between cell cycle time (time between two divisions) and cellular geometric features (Figure 2C). Surprisingly, the cell cycle time was neither correlated with histoblasts’ initial area at birth (Figure 2C) nor with the cell mean area or mean cell elongation over the entire cell cycle time (Figure S2B, C). This suggests that geometric constraints do not strongly influence histoblast proliferation rate, and therefore do not account for the oscillations observed.

We then examined whether cycle times of mother and daughter cells were correlated, but found only a weak correlation coefficient (Figure 2D, E, S2D, Pearson correlation coefficient *ρ* = 0.04 ± 0.04). In contrast, we noticed that sister cell cycle times, and, to a lesser extent, cousin cell cycle times were clearly correlated (Figure 2D, E, S2D, *ρ* = 0.55 ± 0.04 for sisters and *ρ* = 0.31 ± 0.01 for cousins). Clusters of cell cycle synchrony have been suggested to occur in the growing wing disc based on fixed imaging of cell cycle markers (Milan et al., 1996). Do spatial correlations between neighboring cells explain that cousins have correlated lifetimes, despite daughters being uncorrelated with mothers? Indeed, nearest neighbors born within a 30 minute interval of each other had cell cycle times which were positively correlated (Figure 2D, E, S2D, *ρ* = 0.35 ± 0.04). Looking at cycle times for pairs of cells as a function of their distance, the cell cycle time spatial correlation decreased on the scale of a few cells (Figure 2E, S2E). We wanted to test if these measured spatial correlations could explain correlations measured for cousin cells. We therefore obtained correlations between cells that are not cousins but are sharing the same statistical distribution of birth time difference and neighbor relations as cousins (Figure S2F-H, section 1.5 in the Supplementary Theory). Correlations between cycle time for these cells was comparable to cousin correlations, indicating that proximity relations can to a large extent explain cousin-cousin correlations.

Given that cycle times are spatially, but not temporally correlated, can the oscillation in cell division rate be attributed to the relatively narrow cell cycle time distribution (Figure S2A, coefficient of variation of cell cycle times 0.22±0.02, mean ±SD over 4 WTs)? To test this, we simulated a growing population of cells with cycle time chosen stochastically from the experimentally measured cell cycle time distribution. If cells initiate growth sufficiently synchronously, the proliferation rate shows a damped oscillatory behavior, with the oscillation amplitude decaying as the population becomes progressively unsynchronized (see Figure S2I and Figure 3H below). This suggests that a narrow distribution of cell cycle time together with synchronized initiation of growth, can explain the oscillatory behavior in cell division rates.

**Figure 3.**
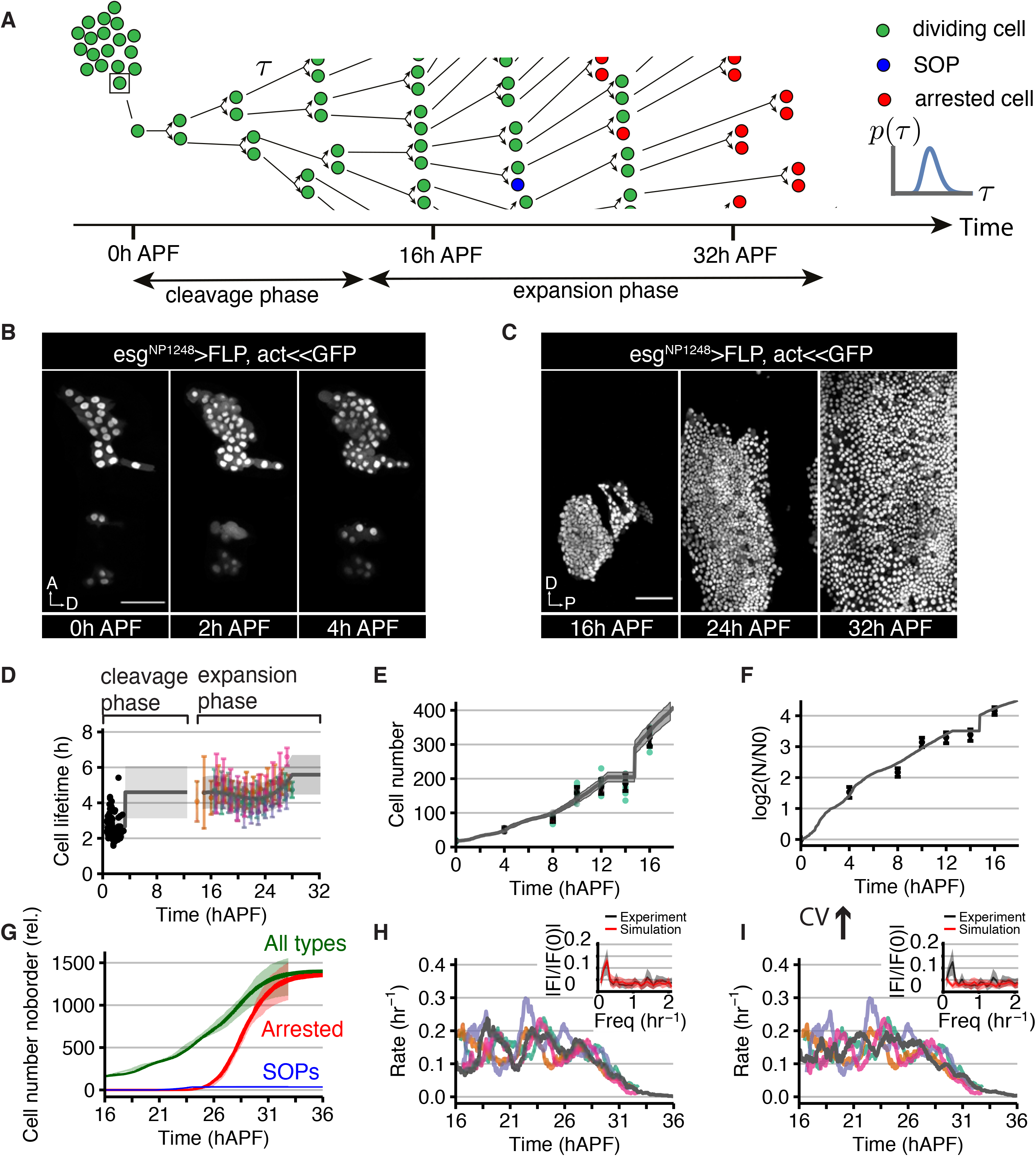
Kinetics of histoblast growth can be explained by changes in cell cycle times and stochastic transition of individual cells to an arrested state. (A) Schematic of histoblast growth simulations. Cells take their cycle time from a probability distribution changing in time and become arrested or SOPs according to time-evolving probabilities. (B, C) Example time-lapse confocal images of histoblast nests at the times indicated during the pre-pupal (B) and pupal (C) stages. Histoblasts are labelled by driving nls-GFP expression in this tissue. In B, the orientation of the animal is rotated compared with other panels. Scale bar = 50 *μ*m. (A = Anterior, D = Dorsal, P = Posterior) (D) Black points: experimental measurements of cycle times available in the first 3 hAPF. Colored points and error bars: binned mean and standard deviation of cycle from WT1-WT4. Grey line and ribbon: mean and standard deviation of cell cycle time inputted to the simulation. In the time period covered by the colored points, the grey line is a 3^rd^-order polynomial fit to the data points. (E) Quantification of anterior nest cell numbers from 0 hAPF until 16 hAPF (green dots, with average and standard deviation indicated in black). During the first 10 h of pupal development histoblasts undergo 3 divisions followed by a transient pause in cell number increase until the expansion phase begins at around 15 hAPF. Grey line and ribbon: mean +/− SD of trajectories from a representative simulation, as described in the text. (F) Data from (E) scaled so that the vertical axis represents the mean number of division cycles per cell. (G) Comparison of normalized cell numbers for different cell classes in the noborder ROI between experimental measurements (lighter ribbons) and base case simulations (darker lines and ribbons). Cell numbers for different WTs are normalized as described in Supplementary Theory. (H) Comparison of the experimental cell division rates within the noborder ROI from each WT movie (colored lines) with a representative simulation (grey line). Moving average applied as in Figure 2A. Inset: Absolute value of the Fourier transform of the division rate data prior to 28.5 hAPF, normalized to its value at 0 frequency. The height and sharpness of the main peak around 0.25 hr^−1^ characterizes the oscillatory component of waves of division. Inset; black: mean and SD from n=4 experiments, red: mean and SD from n=10 simulations. (I) As H but for a simulation in which the CV after 14.7 hAPF is doubled to 0.4. The peaks in the division rate become noticeably less sharp and more disordered, also reflected in the Fourier transformed data (inset).

### Histoblasts transition to growth arrest

We wished to determine why cell division rates decay strongly after ~28 hAPF (Figure 2A). In the *Drosophila* wing growth termination is associated with a gradual increase in cell cycle time (Mao et al., 2013; Martin et al., 2009; Wartlick et al., 2011b). In contrast, histoblast cell cycle times vary little throughout development and do not increase at late stages to an extent that can account for proliferation arrest (Figure 2B). The discrepancy between average cycle times and average cell division rate implies that a fraction of histoblasts stop dividing. We therefore computationally labelled cells that did not divide until the end of our movies, which we denote as arrested histoblasts (Figure 2F). Cell division rates have almost reached zero by the end of each movie, indicating that histoblast cells have largely stopped proliferating by ~31 hAPF (Figure 2A). Labelling these arrested cells revealed a spreading pattern of emerging arrested histoblasts, which progressively cover the entire histoblast nest (Figure 2F, Movie 2). These arrested cells start to appear with a low probability around 24 hAPF and after 28 hAPF, nearly all newborn cells are arrested (Figure 2G). Do arrested histoblasts appear in a systematic spatial pattern within the nest? To test this, we labelled all tracked arrested cells in the final frame of our movies (which, by our definition, have all exited the cell cycle) according to their time of birth (Figure 2H). We did not find a consistent spatial trend across all WT movies (Figure 2H, Movie 2).

We therefore treated the appearance of arrested cells as a stochastic process, and quantified, for every cell division, the probability to give rise to one or two arrested cells (*p*) and, given that at least one of the daughter cells is arrested, the probability that both the two daughter cells are arrested (*α*) (Figure 2I-K). The probability *p* and *α* increased sharply from about 0 before 24 hAPF to almost 1 after 28 hAPF, indicating that histoblasts transition abruptly to proliferative arrest, rather than gradually increasing cell cycle times as reported in other tissues like the *Drosophila* wing disc.

### Simulations recapitulate cell number increase from 0 hAPF to 36 hAPF

We wished to use numerical simulations to test whether (i) cycle times randomly chosen from a narrow distribution and (ii) stochastic transition to cell cycle arrest with a probability changing over time could quantitatively account for histoblast growth (Figure 3A). We first quantified the number of histoblasts between 0 hAPF and 16 hAPF, during the prepupal and early pupal stages (Figures 3B-C). At early times, before ~3 hAPF, we could follow cell division by time lapse and measure an average cell cycle time of ~2.7h (Figure 3D and Movie 3). Later, between ~4 hAPF and ~16 hAPF, continuous live imaging was not possible because of the extensive movements of the pupae. Therefore, we quantified the number of cells in the anterior histoblast nest in fixed images at different times between 0 hAPF and 16 hAPF (Figure 3E, 3F). This indicated that the short cell cycle time measured before 3 hAPF was not maintained at later times (Figure S3A) but instead was slowing down (Figure 3D-E). As previously described (Ninov et al., 2007), our quantification was consistent with roughly 3 cleavage divisions occurring up to ~12 hAPF (Figure 3F). Cell number plateaus between ~12 hAPF and ~14 hAPF, before undergoing a sudden rise between ~14 hAPF and ~16 hAPF; supporting previous observations that growth occurs in two phases, a cleavage phase up to ~12 hAPF and an expansion phase starting after 14 hAPF (Ninov et al., 2007).

Using these data, we performed a simulation of tissue growth where (i) sister cells take a cycle time at their birth out of a bivariate normal probability distribution with a time-varying mean, coefficient of variation (CV) and fixed sister correlation coefficient, (ii) cells pause their divisions between 12.5 hAPF and 14.7 hAPF but still age, giving rise to a burst of cell division around 15 hAPF (iii) cells stop proliferating according to the experimentally measured probability distribution, (iv) cells transition to SOPs with a fixed probability in a time window (Figure 3A, Supplementary Theory). The mean and SD of the simulated cycle time probability distribution before 3.3 hAPF and after 14.7 hAPF were taken according to experimentally measured distributions (Figure 3D, Supplementary Theory). Between these two phases, the mean cell cycle time was taken as the starting value of the expansion phase at ~15 hAPF (Figures 3D, S3B). This model of tissue growth could closely match the increase in the total number of cells, the total number of arrested cells and the number of SOPs over time (Figure 3E-G, see Figures S3C-F for effect of changing model parameters). The simulated cell division rate exhibited oscillations comparable in period and amplitude to experimental data, as well as a similar decay phase (Figure 3H). As expected, oscillations in simulated cell division rate were strongly dependent on the CV of cell cycle time, becoming much flatter with less precise cycle times (Figure 3I). We conclude that the essential features of cell number increase in histoblasts are accounted for by the key mechanisms captured in our simulations.

We noticed that in our simulations, oscillations in cell division rate after 16 hAPF were enhanced by the pause in cell division that we introduced, which tends to resynchronize cell division at the end of the pause, whereas a different implementation of a pause that did not cause such resynchronization resulted in flatter oscillations (Figure S3C). To test whether our simple simulation rules capture the cell division rate oscillation independently of this choice, we simulated tissue growth for different WTs after 16 hAPF, taking the initial cell number and experimental times to division as initial conditions (Figure S3G). The oscillation in cell division rate was still present in these simulations, but in some cases was less pronounced than in experiments (Figures S3H-J). These observations suggest that additional couplings between cells in the tissue might contribute to further enhance the oscillation in cell division rate.

Overall, we identify two key events in abdomen growth: a pause in proliferation between the cleavage and expansion phases, and a sudden transition of individual cells to arrest, ending proliferation. Given the reported effects of tissue mechanics on proliferation in several biological systems, we next investigated whether these key steps were associated to changes in mechanical properties.

### Junctional tension increases during abdomen development

We explored whether mechanical stresses were changing in histoblast nests over time. We performed single junction ablations in different regions of the anterior nest and calculated the recoil velocity of the junction vertices (Figures 4A-E, S4A). We found two distinct behaviors depending on the location of junctions with respect to the LECs: although there was little change in the recoil velocity of junctions at the LEC/histoblast boundary, junctions away from the boundary showed an increase recoil after 21 hAPF, as well as a bias towards the dorso-ventral (DV axis) (Figures 4C-E and S4A). We tested whether this increase in junction recoil after ablation was associated to cell shape changes, but found no clear correlation between junction length and recoil velocity at all developmental stages (Figure S4B). As LECs are contiguous with the histoblasts, we also examined if they experienced changes in tension. Using laser ablation, we excised individual LECs, and followed the subsequent LEC deformation (Figures 4F-H, S4C-E, Movie 4). At 16 and 21 hAPF, LECs showed minimal shape changes (Figure 4G, H, S4C). At 26 hAPF however, there was a large reduction in apical area upon ablation (Figure 4G, H, S4C) and the shape contraction was slightly more pronounced along the antero-posterior (AP) than the DV axis. This is consistent with a recently reported increase in recoil velocity in single LEC junction ablations between 20 and 27 hAPF (Michel and Dahmann, 2020). Surprisingly, we therefore observe that recoil velocities of histoblast and LECs increase over developmental time.

**Figure 4.**
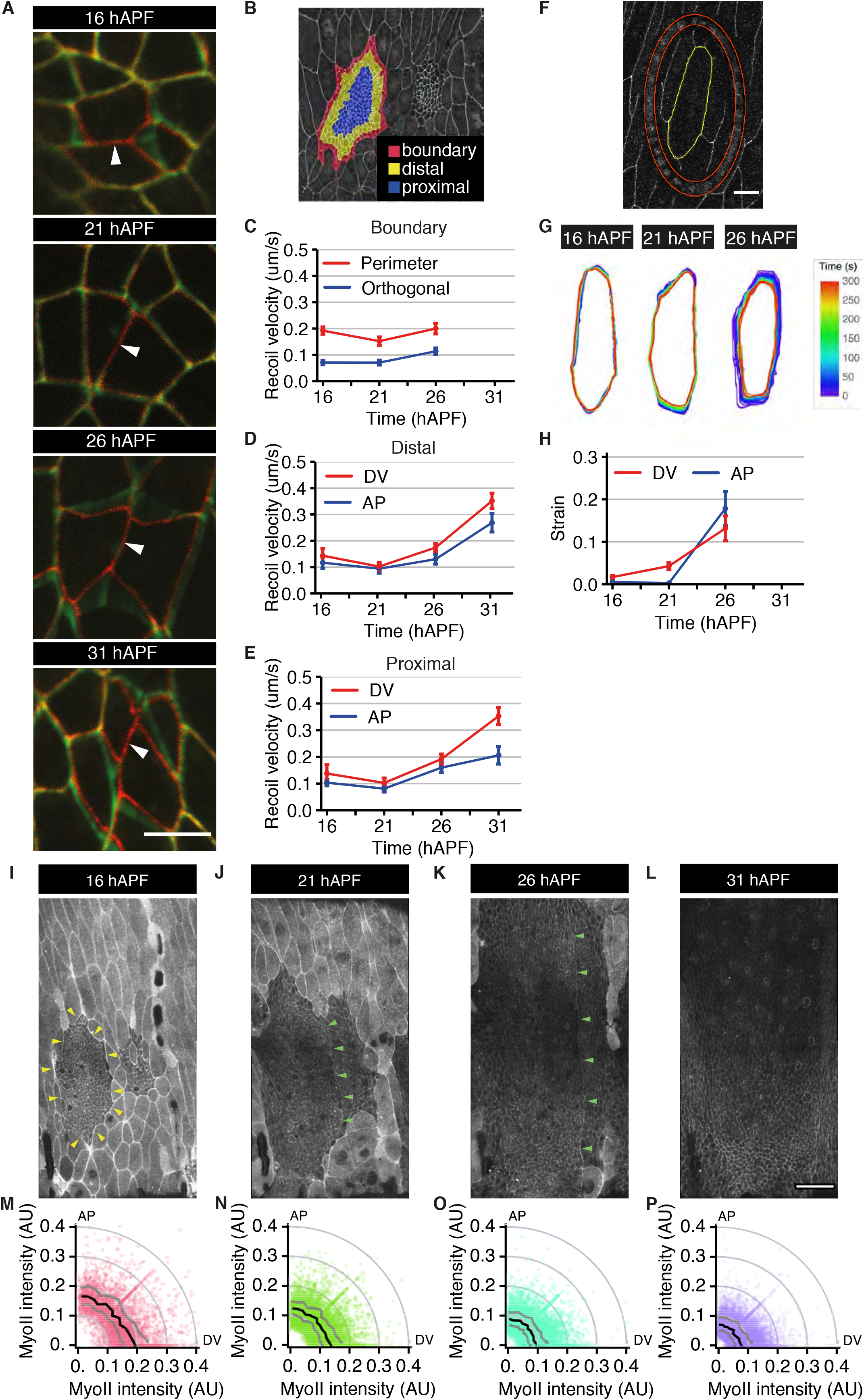
Junctional tension increases through development. (A) Snapshots of laser ablation experiments. Red color; before laser ablation; green color; time frames after laser ablation. Scale bar = 5 *μ*m. (B) Confocal image of dorsal histoblast nests at 16 hAPF where the position of boundary, distal and proximal cells is highlighted. (C-E) Quantification throughout development of recoil velocity for single junction ablations for cells located along the boundary (C, perimeter junctions n=38, 31, 29 and orthogonal junctions n=39, 31, 29 at successive times), or in the distal (D, DV junctions n=16, 19, 30, 20 and AP junctions n=16, 9, 24, 21) and proximal (E, DV junctions n=16, 11, 33, 21 and AP junctions n=21, 22, 38, 22) positions within the nest. (F) Snapshot of an LEC at 16 hAPF in the resting state after an annular ablation in an E-cad∷GFP-expressing animal. Excised LECs were segmented (yellow) and their shape change analyzed to assess strain. (G) Representative LEC segmentations after ablation, temporally overlaid for each developmental stage examined. (H) Quantification of Hencky’s true strain for LEC AP (short) and DV (long) axes, after ablation. (I-L) Example apical projections of confocal images from a pupa expressing Sqh∷GFP (Myo II) and E-cad∷mKate2 at the indicated time points. Note the higher levels of Myo II intensity at the Histoblast-LEC boundary (yellow arrowheads), and the Myo II supra-cellular cable at the histoblast AP nest boundary (green arrowheads) (Umetsu et al., 2014). Scale bar = 50μm. (M-P) Polar-plots of apical Myo II intensity (radius) as a function of junction angle (black line denotes median intensity and dark grey denotes the inter-quartile range) at different times (as in I-L).

Examining Myosin II (Myo II) levels at junctions (Figures 4I-P, S4F, G, Movie 5) showed that the highest levels in histoblasts are at the histoblast-LEC boundary (Figure 4I-K, Michel and Dahmann, 2020), likely explaining the large interfacial recoil after ablation between the histoblast nests and LECs (Figure 4C). However, Myo II junctional levels globally decline throughout development, whilst also switching from a slight intensity bias along DV junctions at 16 hAPF to being isotropic at 31 hAPF (Figure 4M-P). We also observed no strong dependency of Myo II intensity on junction length (Figure S4F). We conclude that the increase in recoil velocity over time is not linked to an increase in Myo II junctional levels.

Junctional tension is only one of the components generating forces in the tissue, so we wondered if the large-scale tissue tension was changing over time in the same way as junctional recoil velocity. To test this, we performed apical annular cuts with a diameter of about 10 cells in a defined region of the anterior histoblast nests at different time points (Figure 5A, Movie 4). Consistent with observations of single-junction ablations, the strain and recoil velocity measured in the AP and DV directions were increasing over time (Figure 5B, S5A, B, Star methods). In addition, the spatial pattern of cell area contraction within excised discs exhibited a striking change over time. Cell area contraction was mostly confined to the disc edge at 16h, and this edge bias disappeared at 31 hAPF (Figure 5C). Since a free elastic disc under isotropic tension should constrict uniformly (Supplementary Theory), we reasoned that this observation indicated that cell movement was initially limited by an external elastic resistance. We therefore compared experimental patterns of isotropic shear (relative cell area change) and anisotropic shear (change in cell elongation) to a continuum model where the tissue is described as an elastic material under active tension, adhering to an external substrate through elastic links (Figure 5D-H, S5D-F). Fitting this model to spatial profiles of excised discs and tissue outer boundary deformation at different times (Figure 5E), we found that the parameter describing external resistance to the disc deformation was strongly decreasing over time (Figure 5F). The model also indicated that the tissue internal AP tension was roughly constant over time (Figure 5G), while the DV tension increased after 26 hAPF (Figure 5H). We therefore conclude that build-up of compressive stresses does not occur in the histoblast nest, and therefore is not responsible for growth arrest as has been suggested for other tissues (Irvine and Shraiman, 2017)

**Figure 5.**
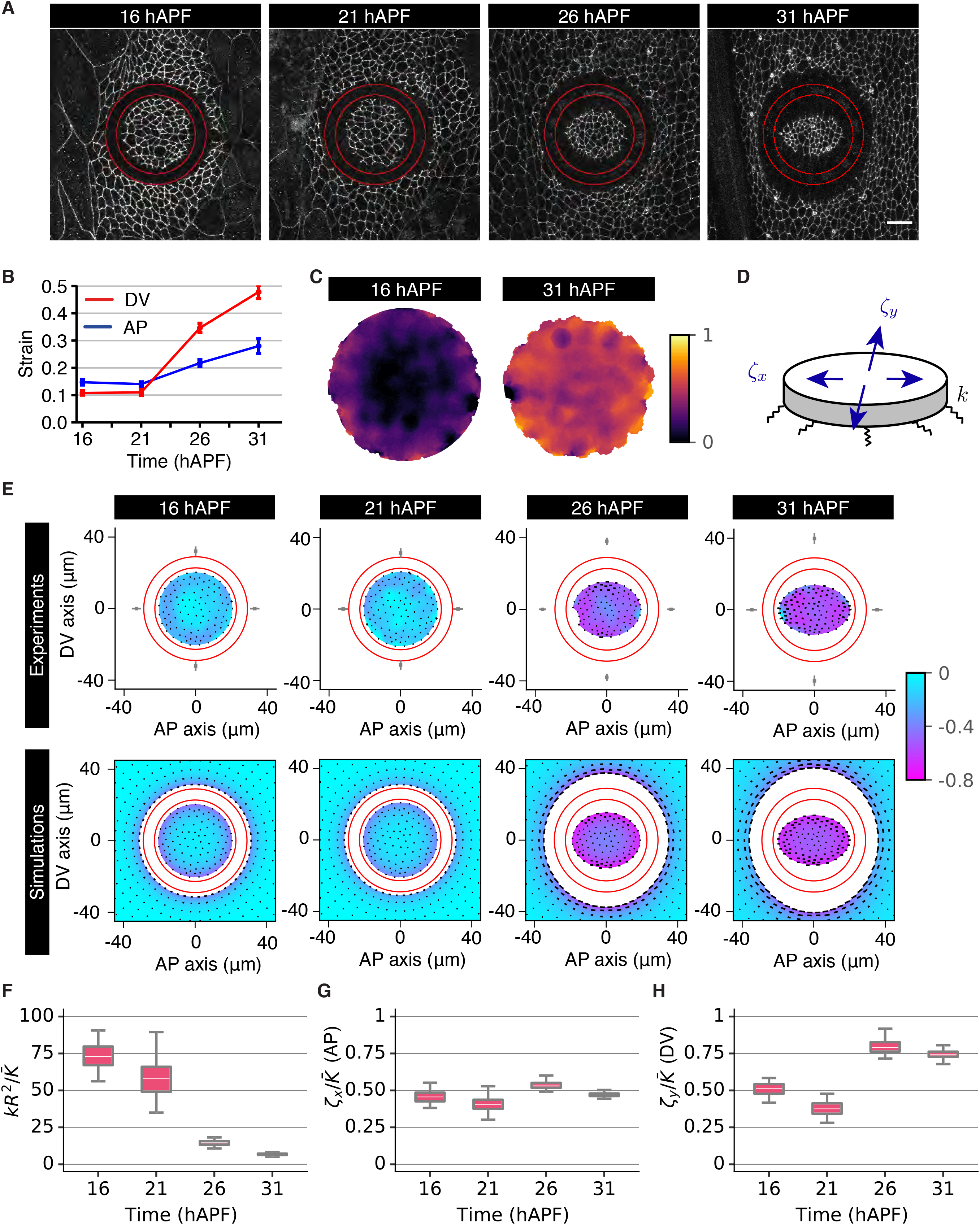
Deformation of excised histoblasts reveals a reduction in external resistance to histoblast movement. (A) Example time-lapse confocal images of histoblasts at the resting state following an annular ablation (ablated region outlined in red) in E-cad∷GFP-expressing animals, at different stages of abdomen development. Scale bar = 5μm. (B) Quantification of Hencky’s true strain in the Dorsal-Ventral (DV) and Anterior-Posterior (AP) axes of excised histoblasts throughout development. n=23, 24, 29, 16 experiments at successive times. (C) Averaged spatial map of relative area change after excision at 16 hAPF and 31 hAPF, plotted on the undeformed discs (n=20 (16hAPF) and n=15 (31 hAPF) experiments). As time progresses, the relative area change becomes more homogeneous in the disc (D) The excised disc is described as an elastic material, subjected to anisotropic active tension *ζ*_*x*_, *ζ*_*y*_, adhering to a substrate through elastic links, with effective elastic modulus per area *k*. (E) Experiment (top) and simulation (bottom) deformation plots of excised histoblast discs. Color code: relative area change, black lines: anisotropic shear or change in cell elongation. Deformation fields are plotted on the deformed disc and are obtained from measurements before and 3 minutes after ablation for experiments. Red circles indicate ablated region. Gray squares and error bars in experimental plots: deformation of the outer circle of the ablated ring (mean±95% confidence interval), averaged between top/down and left/right deformations. Data is obtained from n=20, 12, 13, 15 experiments at successive times. (F, G, H) Fitted model parameters to excised disc deformations, as a function of time. (F) Normalized ratio of external elastic modulus per area *k* to tissue elastic modulus 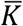. (G, H) Normalized AP and DV tensions, respectively.

### The basal extracellular matrix is degraded during histoblast expansion from 13 hAPF

Why does the external resistance to tissue deformation appear to decrease over time? The apical surface of the histoblasts and LECs is in contact with the pupal cuticle, while the basal surface is attached to a basement membrane containing Collagen IV (Viking – Vkg in flies) (Ninov et al., 2010 and Figure S6A). We therefore quantified basal extracellular matrix (ECM) dynamics during pupal abdominal development. We investigated the dynamics of the three major *Drosophila* basal ECM components, Perlecan (*Drosophila* Trol), Collagen IV and Laminin B1 (LanB1) from 4 hAPF to 32 hAPF. During the pre-pupal stages (4 hAPF – 12 hAPF), all three components are present as a dense network across the entire abdomen (Figure S6B, C), but are slowly degraded (Figure S6B, C, Movie 6). At 12 hAPF, head eversion compresses the ECM network along the AP axis, leading to an increase in ECM component intensity (Movie 6, Figure S6B, C). This is followed by degradation of all three ECM components across the entire abdominal region from around 13 hAPF, consistent with previous work indicating that Collagen IV under the histoblasts is degraded between 16 and 28 hAPF (Ninov et al., 2010). We found that Perlecan is degraded at a faster rate than Collagen IV and Laminin B1 (Figure S6B, C, Movie 6).

We examined at higher spatial resolution whether the basal ECM is degraded in a similar manner under the histoblasts and LECs, after 16 hAPF. Both Perlecan and Collagen IV are degraded at a similar rate under both cell populations, and by 21 hAPF we only detect a residual punctate signal in hemocytes (Figure 6A, B, Movie 6). LanB1, on the other hand, is degraded under the LECs, but is maintained at a higher level under the histoblasts than under the LECs (Figure 6A, B, Movie 6). To identify potential mechanisms for ECM degradation, we imaged a transcriptional reporter for the secreted *Drosophila* matrix metalloprotease, MMP1 (Wang et al., 2010). *mmp1* is expressed in cells below the epidermis (e.g. muscles, fat body - FB and hemocytes), and weakly in the polyploid LECs, but not in the histoblasts (Figure 6C). *mmp1-GFP* levels rapidly increased during histoblast expansion, suggesting that MMP1 release by the underlying cells triggers the remodeling of the ECM (Figure 6C, D, Movie 7).

**Figure 6.**
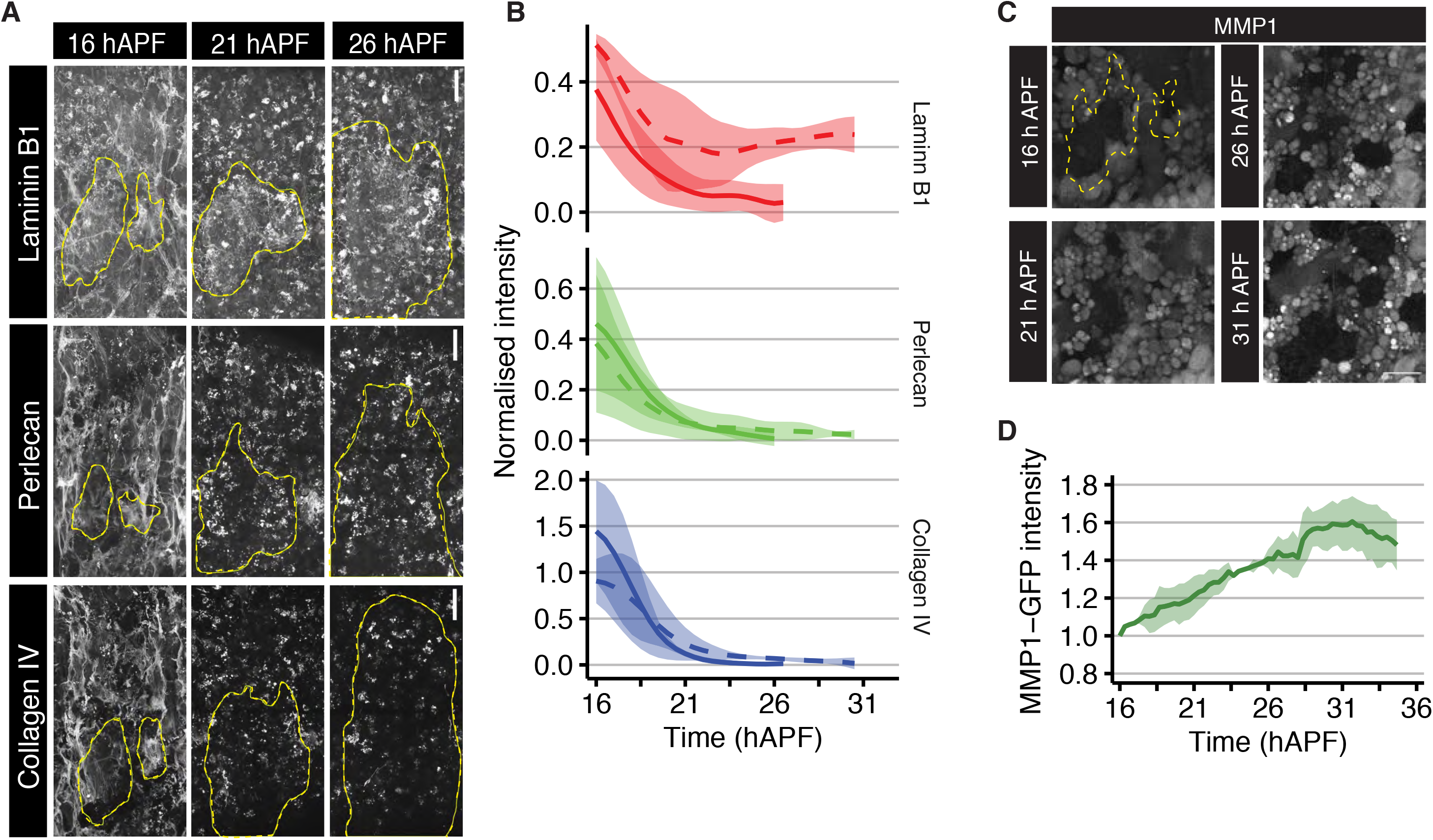
Basal extracellular matrix remodeling. (A) Example maximum intensity projection of confocal images of the three major basal ECM components during pupal stages. Histoblast nests are outlined in yellow. Scale bars = 50μm. (B) Quantification of ECM components intensity underneath LECs (solid lines) and histoblasts (dotted lines), after subtraction and normalization to the lowest measured ECM component intensities under LECs (see STAR Methods), throughout pupal development (error bars: SD, n=3 for each genotype). (C) Still images of MMP1-GFP reporter expressing pupae in the basal hemolymph throughout development. Scale bar = 50 *μ*m. Histoblast nests are outlined in yellow. (D) Quantification of MMP1-GFP maximum projection intensity in the basal hemolymph, measured every 20 minutes (Error bars: SD, n=2). Intensities are normalized to the mean intensity at 16 hAPF.

To test whether the reduction in basal ECM components caused the loss of external mechanical resistance inferred from annular ablations, we overexpressed MMP1 or TIMP (Tissue inhibitor of metalloproteases, an endogenous MMP inhibitor) in the LECs (*32B-GAL4* driver) and performed annular cuts on the histoblasts. Area contraction was less concentrated at the edge of excised discs in pupae overexpressing MMP1 than in WT at 21h APF (Figure 7A) and the disc overall deformation magnitude was more pronounced at early times and comparable at later times (Figure 7B, C, S7A-D). In contrast, in TIMP overexpression at 26 hAPF, the area deformation profile was concentrated near the boundary of the disc and the overall deformation magnitude was reduced compared to WT (Figure 7A-C, S7E-H). These results are intuitively consistent with the ECM providing external resistance to tissue deformation; as resistance decreases when the ECM is degraded early (MMP1 overexpression), and increases when the ECM persists for longer (TIMP overexpression). To obtain a quantitative read-out, we fitted our continuum model to experimental deformation profiles in MMP1 and TIMP overexpression, as for annular ablations in WT pupae (Figure 5D). This confirmed that external resistance to the disc deformation was decreasing early in MMP1 overexpression and was strongly increased in TIMP overexpression (Figures 7F, G). Tissue tension was generally similar to WT in these perturbations, except for an increase of isotropic tension at 16 hAPF in MMP1 overexpression (Figures 7H, I and S7I, J). Overall, these results are consistent with a progressive disappearance of essential ECM components after 13 hAPF, resulting in reduced external resistance to deformation of the histoblast epithelium.

**Figure 7.**
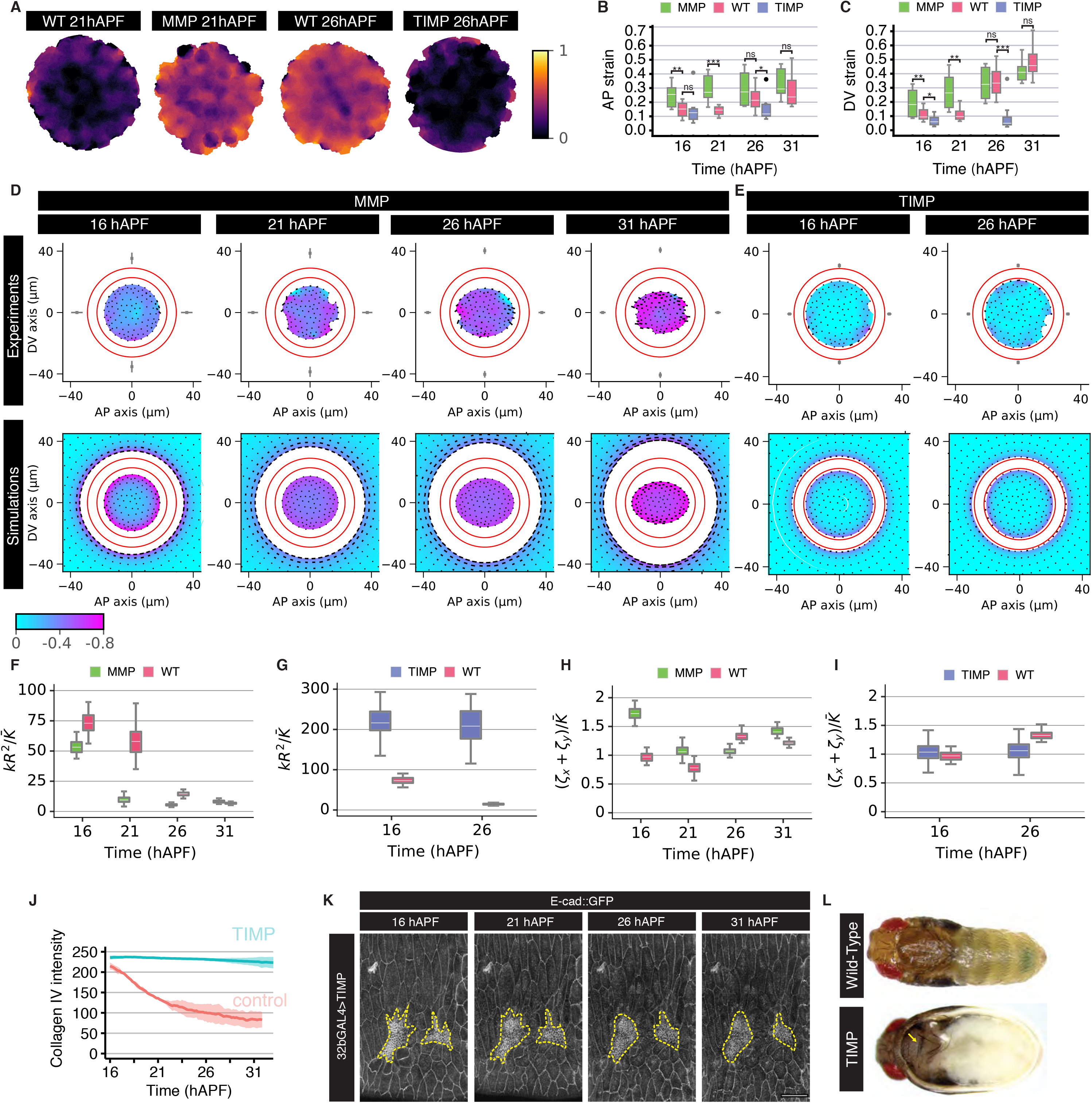
Basal ECM remodeling is necessary for histoblast nest expansion. (A) Averaged spatial map of relative area change after excision at 21h APF for MMP and WT, and at 26 hAPF for TIMP and WT, plotted on the undeformed discs. The relative area change is more homogeneous in the MMP perturbation than in the WT, indicating reduced external resistance to tissue deformation. From left to right, n=12, 7, 13, 5 experiments. (B, C) Quantification of Hencky’s true strain along the AP (B) and DV (C) axes, calculated from annular ablations in pupae expressing MMP1 or TIMP under the control of *32B-GAL4*. WT data is repeated from Figure 5. Mann-Whitney test was used to compare populations, with 0.05>*>0.01>**>0.001>***. From left to right for both plots n=12, 23, 7, 7, 24, 7, 29, 9, 6, 16 experiments. (D, E) Experiment (top) and simulation (bottom) deformation plots for excised histoblast discs, at 4 different time points, in pupae expressing MMP1 (D) or TIMP (E) under the control of *32B-GAL4*. Representation is as in Figure 5E. From left to right, n=11, 7, 7, 6, 7, 5 experiments. (F-I) Fitted model parameters for excised discs deformation, as a function of time, compared to parameters for WT (red, data as in Figures 5F, S5D-E). (F, G) Normalized ratio of external elastic modulus per area *k* to tissue bulk elastic modulus 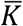 in pupae expressing MMP1 (green, F) and TIMP (blue, G). (H, I) Normalized isotropic (sum of AP and DV tensions, *ζ*_*x*_+*ζ*_*y*_) tensions in pupae expressing MMP1 (green, H) and TIMP (blue, I). (J) Quantification of the mean intensity of Collagen IV from movies of pupae expressing HA (control) or TIMP under the control of *32B-GAL4* (Error bars: SD, HA n=2, TIMP n=2). (K) Stills from a movie of a pupa expressing E-cad∷GFP, and overexpressing TIMP under the control of *32B-GAL4* at the time points indicated. Scale bar = 50 *μ*m. Histoblast nests are outlined in yellow. (L) Images of WT and TIMP expressing pharate pupae. Yellow arrow indicates the normal development of the thorax. In contrast, the abdominal cuticle is unpigmented and devoid of sensory bristles.

### Basal ECM remodeling is necessary for histoblast nest expansion

The ECM degradation that occurs between 14 and 21 hAPF coincides with the transition from the histoblast cleavage divisions to the expansion divisions (Figure 3). We therefore investigated whether blocking ECM remodeling has an effect on histoblast proliferation by overexpressing TIMP, which as expected prevented Collagen IV degradation (Figure 7J, S7K, Movie 7). Next, we analyzed the behavior of the histoblasts while preventing basal ECM degradation with TIMP overexpression (Figures 7K). No proliferation was observed in the histoblasts during the time period observed, from 16 to 31 hAPF (Figure 7K, Movie 7), which is strikingly different from extensive proliferation seen in the wild type histoblasts over the same time period (Figure 1B). The lack of proliferation was not due to early pupal death since our experimental samples developed to the pharate adult stage with a normal head, thorax and wing (Figures 7L, S7L). Moreover, a pupal wing imaged simultaneously with the abdomen in a TIMP-expressing pupa proliferated normally at 16 hAPF (Figure S7M, Movie 7). In contrast, the larval abdomen failed to be replaced by the histoblasts, as evidenced by the lack of pigmentation and sensory bristles (Figure 7L, S7L). Thus, the expansion phase of cell division in the histoblasts fails to start in animals overexpressing TIMP. This provides evidence that basal ECM remodeling triggers the start of cell growth and proliferation during *Drosophila* abdominal epithelium expansion.

## Discussion

Our quantitative analysis has allowed us to identify the major landmarks of abdomen growth. We find that following ~3 cleavage divisions, a pause in proliferation rate precedes an expansion phase. After ~ 4 cell divisions (Table S1), the expansion phase ends with cells individually transitioning to arrest. Beyond this global picture of histoblast growth, our dataset and simulations provide several surprising insights.

### Correlations in cell cycle times

We find that cell cycle times are not temporally correlated, nor are they dependent on cell geometry, but exhibit spatial correlations on a scale of a few cells (Figures 2, S2). Tracking of single cells in culture has led to the identification of a pervasive phenomenon known as the “cousin-mother inequality”, whereby cousins display a surprising degree of cell cycle time correlation given weak mother/daughter correlations (Mosheiff et al., 2018; Sandler et al., 2015). This phenomenon has been attributed to the interplay of the cell cycle machinery with the circadian clock (Chakrabarti et al., 2018; Sandler et al., 2015), or the inheritance of cell mass or cell cycle regulators over several generations (Kuchen et al., 2020). In our developing *in vivo* system, we propose that spatial correlations largely explain the “cousin-mother inequality” in cell cycle time correlations. While we cannot distinguish whether these spatial correlations arise from cell-cell communication or exposure of neighbors to a common microenvironment, we note that spatial patterns of cell cycle time vary significantly in time and between pupae (Figure S2E). We speculate that these neighbor correlations are a signature of cell-cell communication mechanisms which might also ensure that tissue-wide events, such as transition to arrest take place in a coordinated manner.

### Growth termination: a gradual or abrupt process?

The *Drosophila* wing imaginal disc is arguably the system in which growth and final size control have been studied most extensively (Gou et al., 2020). A striking aspect of this system is that cell proliferation decelerates progressively during roughly half of the larval growth period (Gou et al., 2020). A similar scenario was observed during postnatal growth of the mouse skin, where the cell cycle time of epidermal progenitors gradually increases over sixty days following birth (Dekoninck et al., 2020). In contrast, we find that termination of tissue growth is not mediated by a progressive increase in cell cycle time, but by the sharp, stochastic transition of a growing population of cells to proliferative arrest, with no visible large-scale pattern (Figure 2). This is reminiscent of models which have been proposed for growth and differentiation of neurons in the retina (He et al., 2012), growth of mouse intestinal crypts (Itzkovitz et al., 2012) or the expansion of the mouse embryonic skin (Lechler and Fuchs, 2005). In these tissues, a relatively abrupt transition in modes of progenitor divisions switch the tissue from expansion to maintenance and differentiation. This suggests that different developing tissues can achieve reproducible sizes using radically different growth termination strategies. It is possible that the sharp transition we observe here allows for fast expansion of the histoblast nest, at the cost of a less refined control over the final number of cells than would be allowed by a progressive slow-down in proliferation, as observed in the wing disc or mouse skin (Itzkovitz et al., 2012).

### Mechanical control of developmental growth

Mechanical control of growth is an attractive model for tissue size determination supported by the study of the effects of cellular density on the proliferation of tissue culture cells (Irvine and Shraiman, 2017; Panciera et al., 2017). In this scenario, crowding leads to cell cycle arrest as cells reach confluence through compaction, driving a reduction in cell area. By analogy to cells in culture gradually filling empty space on the culture dish surface, it is tempting to speculate that histoblasts, which grow within a confined planar surface, are subjected to mechanical crowding leading to growth termination. However, we did not find a signature of a feedback of cell area affecting the cell division rate. Furthermore, laser ablation experiments indicate that the tissue tension is either constant (in the AP direction) or increasing (in the DV direction) (Figure 5, S5). Decreased Myo II accumulation has been proposed to mediate the effect of cell crowding on proliferation in the wing disc (Pan et al., 2018; Rauskolb et al., 2014). However, though an area of lower apical Myo II levels emerges in the medial part of the anterior dorsal nest from around 21 hAPF (Figure S5F), this does not lead to a similar pattern of increased cell cycle time or early exit from the proliferation phase. Finally, the transition from growth to arrest in the abdomen occurs between ~24 and ~28 hAPF, whereas the displacement of the last LECs and fusion of the dorsal nests at the midline takes place between 32 and 36 hAPF (Madhavan and Madhavan, 1980; Michel and Dahmann, 2020). Thus, changes in cell geometry or mechanical tension do not appear to provide a universal growth termination cue.

### Matrix remodeling and tissue growth control

The transition between the prepupal cleavage divisions to the expansion phase represents a key step in abdominal development. This involves both a lengthening of the cell cycle, and a resumption of tissue growth. The depletion of Cyclin E stores accumulated during the larval growth period is thought to account for the increase in cell cycle length (Ninov et al., 2007). Here, we show that remodeling of the basal ECM is an essential step to allow nest expansion to take place. Dynamic remodeling of the ECM plays an instrumental role in organ development by orchestrating processes such as cell migration and rearrangements (Ramos-Lewis and Page-McCaw, 2019; Walma and Yamada, 2020). Partial degradation of the basal ECM (Collagen IV and Perlecan are degraded, while some Laminin persists) initiates at around 13 hAPF and is required for histoblast expansion to occur. Indeed, blocking ECM degradation through the expression of TIMP prevents the transition to expansion divisions (Figure 7K, L).

How is the degradation of the basal ECM triggered at the correct time? MMP1 and MMP2, the two *Drosophila* MMP family members, are responsible for most basement membrane turnover, and have been implicated in the remodeling of a number of tissues during metamorphosis (Diaz-de-la-Loza et al., 2018; Ramos-Lewis and Page-McCaw, 2019). We did not detect high levels of the MMPs in the histoblasts or LECs, but MMP1 levels dramatically rose in the FB and hemocytes beneath the epidermal epithelial layer (Figure 6C, D). The hemocytes might be implicated in epidermal basement membrane remodeling, since they are highly active at that stage and take up much of the GFP-tagged Collagen IV and Perlecan in large phagocytic vesicles (Figure 6A). Alternatively, the release of MMPs into the hemolymph as the FB disperses might trigger the degradation of the epidermal ECM and histoblast nest expansion. Indeed, in response to the pupariation ecdysone pulse, FB cells secrete MMP1 and MMP2, causing the destruction of the cell-cell junctions and ECM that hold them together between 6 and 12 hAPF (Bond et al., 2011; Jia et al., 2014).

How does ECM degradation enable tissue growth? One possibility is that loss of signaling from the ECM via integrins signals the onset of the expansion divisions. However, this would be unexpected as in many systems, integrin signaling promotes rather than inhibit growth and proliferation (Hamidi and Ivaska, 2018). A second possibility is that the histoblast nests are prevented from expanding because cell motion is impaired by strong adhesion to the ECM: the increased tissue-ECM attachment observed in TIMP pupae (Figure 7G) might explain the lack of growth of the histoblasts in this condition. Finally, basement membrane remodeling might free a source of trapped growth factor/morphogen, or allow a ligand present in the hemolymph to access the histoblasts. Indeed, the ECM has been shown to act as a growth factor reservoir (Bonnans et al., 2014; Hynes, 2009) or to restrict morphogen diffusion (Ma et al., 2017; Tian and Jiang, 2014; Wang et al., 2008) in several tissues. It is interesting to consider this repressive effect of the ECM on developmental proliferation in the context of the proposed role of the normal cellular microenvironment in limiting cancer formation (Bissell and Hines, 2011). It is likely that oncogenic transformation involves the exploitation of developmental ECM remodeling mechanisms to convert the ECM from an anti-tumor to a pro-tumor environment that promotes proliferation and tissue invasion (Bissell and Hines, 2011; Chang and Chaudhuri, 2019).

## Acknowledgements

We thank Y. Bellaïche, D. Bohmann, B. Stramer, K. Irvine and the Bloomington Drosophila Stock Centre for fly stocks. We are grateful to M. Renshaw (Crick Advanced Light Microscopy facility) and the Crick Fly Facility for support, and R. Etournay, M. Popovic, M. Merkel and H. Brandl for help with Tissue Miner. We are grateful to B. Aerne for generating the UAS-HA fly stock. We thank JP Vincent and J. Briscoe for critical reading of the manuscript. JRD is funded by a Sir Henry Wellcome Fellowship (201358/Z/16/Z). AF is funded by the European Union’s Horizon 2020 research and innovation programme under the Marie Skłodowska-Curie grant agreement MSCA-IF-EF-ST No 795060. This work was supported by a Wellcome Trust Investigator award (107885/Z/15/Z) to NT. Work in the Salbreux and Tapon labs was supported by the Francis Crick Institute, which receives its core funding from Cancer Research UK (FC001317, FC001175), the UK Medical Research Council (FC001317, FC001175), and the Wellcome Trust (FC001317, FC001175).

## Author contributions

Conceptualization: all authors

Funding acquisition: JRD, AF, GS, NT

Experiments: APA, JRD, AF

Theory, simulations: JW, ATS, GS

Experimental methodology: FM, EMB

Software: JRD, JW, ATS, AH, MBS

Supervision: GS, NT

Writing – original draft: APA, JRD, JW, AF, AH, MBS, GS, NT

Writing – review & editing: all authors

## Movie Legends

**Movie 1. Overview of imaging and quantification pipeline to assess tissue growth dynamics of the *Drosophila* pupal abdomen.**

Example movies for WT1 after the initial movie has been processed through the image analysis pipeline highlighted in Figure S1.

**Movie 2. Appearance of arrested histoblasts shows no consistent spatial pattern.**Overlays of SOPs, dividing histoblasts and arrested histoblasts for each WT movie as mentioned in Figure 2.

**Movie 3. Example movie of early histoblast cleavage divisions.**Histoblasts were labelled with nls-GFP.

**Movie 4. Annular ablations on histoblasts and LECs.**

Example movies of annular ablations on WT, MMP and TIMP over-expression histoblasts and WT LECs at the indicated developmental timepoints.

**Movie 5. Myo II apical intensity during histoblast development.**

Movie of the apical junctions of histoblasts labelled with Ecad∷mCherry and Sqh∷GFP during development. Scale bar=50μm.

**Movie 6. Basal ECM remodeling during the pre-pupal and pupal stages.**

Example movies of the basal ECM components LamininB1, Perlecan and Collagen IV during pupal development. Scale bar = 50μm.

**Movie 7. Histoblast and pupal wing epithelial dynamics with over-expression of TIMP.**Both the pupal wing and histoblasts were imaged at the same developmental time, but whilst histoblasts are arrested, cell divisions still occur in the pupal wing.

## STAR Methods

### RESOURCE AVAILABILITY

#### Lead contact

Further information and requests for resources and reagents should be directed to and will be fulfilled by the Lead Contact, Nic Tapon.

#### Materials availability

All fly stocks are available upon request or from the Bloomington *Drosophila* Stock Center (see key resources table).

#### Data and Code Availability

Data and code are available upon request from the corresponding authors. The software developed for the analysis of annular ablation experiments is provided free of charge via a Github repository (see key resources table).

### EXPERIMENTAL MODEL AND SUBJECT DETAILS

All experiments were performed in *Drosophila melanogaster* (see key resources table and figure genotypes table for details of strains used). Flies were maintained at 25°C (unless otherwise indicated) on food generated by the Francis Crick Institute Media Facility (360 g agar, 3600 g maize, 3600 g malt, 1200 mL molasses, 440 g soya, 732 g yeast extract, 280 mL of acid mix (500 mL propionic and 32 mL orthophosphoric acid) and 50 L water).

### METHOD DETAILS

#### FLY GENETICS AND IMAGING

##### Fly rearing

Flies were raised at 25°C. Pupae were collected at a pupal stage before head eversion, kept at 25°C and then monitored hourly for head eversion to calculate pupal age as 12 hAPF, four hours prior to imaging at 16 hAPF.

The *UAS-GAL4-GAL80*^*ts*^ system was used as (Brand and Perrimon, 1993; McGuire et al., 2004). We induced *32B-GAL4* driven expression of TIMP and HA (control) in a temperature sensitive manner using the *tub-GAL80*^*ts*^ system, co-expressing *E-cad∷Tom* and *vkg-GFP*. Virgins of the following genotype: *E-cad∷Tom*, *vkg-GFP*; *32BGAL4* were crossed to homozygous males *tub-GAL80*^*ts*^; *UAS-TIMP* and *UAS-HA* (II). Crosses were kept at 25°C. Larvae were raised at 18°C and placed at 29°C for 7 hours prior to imaging. Pupae were collected at a pupal stage before head eversion, kept at 29°C and then monitored hourly for head eversion to calculate pupal age as 12 hAPF, three hours prior to imaging at 16 hAPF.

##### Imaging

Pupae were dissected and mounted as described (Mangione and Martin-Blanco, 2020). All movies were acquired on a Zeiss LSM 880 confocal microscope at 25°C, with a Plan-Apochromat 40x/1.3 oil DIC M27 objective, unless stated. For the dorsal-lateral field of view, images were acquired as two-tiles of 1024×1024 pixels with a 10% overlap, with 20-30 z-slices 1μm apart, and the tiled stacks were fused in Zen Blue. WT *E-cad∷GFP* movies were acquired with a frame every 2.5 minutes, TIMP overexpression movies with *E-cad∷GFP* every 5 minutes, *E-cad∷mCherry* and ECM components during the pupal stages every 10 minutes, *mmp1-GFP* every 20 minutes, and the *E-cad∷mKate2; Sqh∷GFP* movies every 5 hours. The TIMP overexpression movies with *vkg∷GFP* were filmed at 29°C, every 20 minutes.

For imaging of the ECM during pre-pupal stages, pre-pupae were placed dorsal-laterally with their posterior in Voltalef 10s to image the abdominal segments. Movies were filmed on a Zeiss LSM 880 confocal microscope at 25°C, with a Plan-Apochromat 40x/1.3 oil DIC M27 objective. Images were acquired as two by four tiles of 512×512 pixels with 10% overlap, 24 z-slices 2μm apart every 30 minutes, and the tiles fused in Zen Blue.

##### Laser ablations

Pupae were dissected and mounted as above. All movies were acquired on a Zeiss LSM 780 confocal microscope with a Coherent Chameleon NIR tunable laser, at 25°C. For single junction ablations, an alpha Plan-Apochromat 63x/1.46 Oil Korr M27 objective was used to acquire a single tile of 512 × 512 pixels with 5x zoom every 500ms for 31 seconds. Junctions were ablated with a wavelength of 780nm at 35% power with a 4 x 25 pixel ROI positioned across the junction with the ablation taking 0.254ms. The vertex location was identified manually and the recoil velocity calculated using the methodology outlined in (Liang et al., 2016).

For annular ablations, a Plan-Apochromat 40x/1.3 oil DIC M27 objective was used to acquire a single 512 x 512 pixel tile with 5 z-slices 1μm apart every 5s for 5 – 10 minutes. For generating ROIs, the freehand ROI tool was used to draw between two circles/ellipses. For histoblasts, the ROI was a circular annulus with an inner diameter of 45.54μm (110 pixels) and an outer diameter of 57.96μm (140 pixels), and the multi-photon was set at 780nm between 25–35% power, and the ablation was of a single z-plane at the apical surface and lasted 9.3ms. For LECs, an elliptical annulus ROI with inner diameter dimensions of 28.98μm (70 pixels) by 57.96μm (155 pixels) and outer dimensions of 38.916μm (94 pixels) by 57.96μm (185 pixels), and the multi-photon was set at 780nm at 40% power, and the ablation was of a single z-plane and lasted 8.3ms. To analyze the various parameters, we followed the methodology outlined in (Bonnet et al., 2012). Briefly, cells within the inner disc were segmented using Skeletor and the inner disc axes lengths measured. To obtain strain, a bounding ellipse was fitted to calculate the long and short axis and then the ellipse matrix was multiplied by a rotation matrix and transpose rotation matrix, to calculate the deformation along the DV (y) and AP (x) axes; the shear component was close to zero. Strain was then calculated by taking the natural log of the non-ablated axis length (L) divided by the relaxed axis length (Lr); (*ε* = ln(L/Lr). To calculate disc recoil velocity, we measured the change in axis length between successive frames. To calculate relaxation time, we took the inverse of the gradient from a straight line model fitted to recoil velocity as a function of axis length.

##### Quantification of cell number in early pupal stages

For pupal staging, white pupae (0 hAPF) were collected with a paintbrush. After selection, animals were transferred to fresh vials and allowed to develop at 25°C until time of dissection. Before dissection pupae were gently cleaned with 1xPBS and placed in double sided tape. First, both the anterior and posterior ends were cut with scissors. Then, pupae were bisected laterally along the antero-posterior axis. Animals were then transferred to sterilized 1x PBS and the internal organs were cleaned from the epidermis by flushing with 1x PBS. Using forceps, the epidermis was detached from the pupal case and transferred to an eppendorf. Fixation was performed for 20 minutes in 4% formaldehyde. After fixation, the epidermis was rinsed three times in 1xPBS (3×5 minutes). Finally, the tissue was equilibrated in Vectashield (with DAPI) overnight and mounted on coverslips.

For the first hours of pupa development, 0 hAPF animals were selected and imaged every 5 minutes for the following 4 hours to allow for manual scoring the time of first and second divisions.

##### Myo II, ECM and MMP1 quantification

To quantify apical Sqh∷GFP intensity throughout development, movies were generated of E-cad∷mKate2; Sqh∷GFP as previously mentioned with a five hour frame rate to minimize photo-bleaching. Using the E-cad channel, the apical surface was projected as mentioned previously, for both E-cad and Sqh channels. The E-cad was then segmented using the AI algorithm and manually corrected. The intensity of Sqh was calculated for each junction, as well as the length and angle of junction relative to the DV axis.

To quantify ECM dynamics, movies were acquired as mentioned previously. For analyzing pre-pupal dynamics an ROI of the final abdominal position within the field of view was generated and the mean intensity of ECM components measured over-time. Intensity data was normalized to the value at 13 hAPF for each movie which is the first timepoint after head eversion where all movement has ceased. For analyzing pupal dynamics an ROI was generated for the ECM under histoblasts and LECs, and the intensity for each cell population calculated over-time. Values were subtracted and normalized to the mean value of the five frames with the lowest mean intensity under the LECs.

##### Figure genotypes

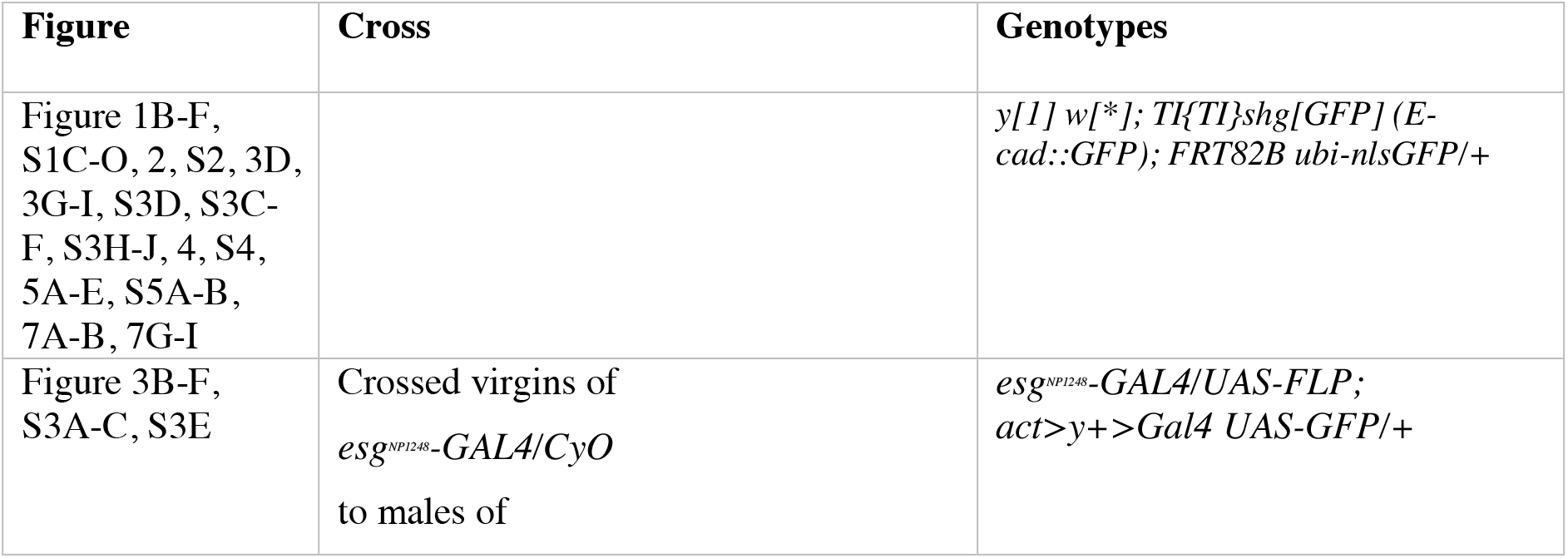
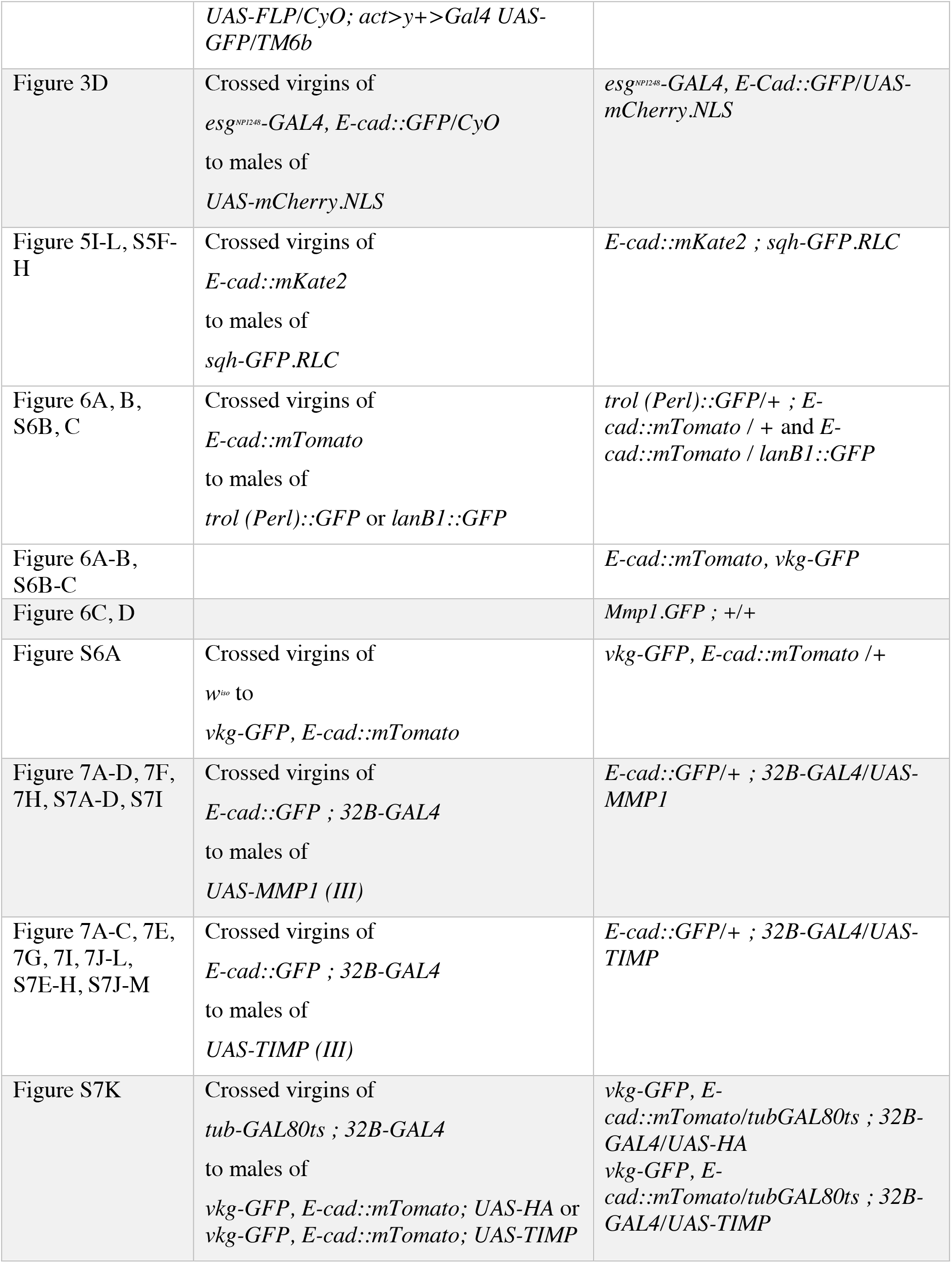

#### MOVIE SEGMENTATION AND ANALYSIS

##### Overview of the segmentation and tracking pipeline

The aim of the image analysis pipeline was to identify, track and classify individual cells within the developing histoblast nests. The image processing was performed on projected cell surfaces which were subsequently segmented (skeletonized) first before being tracked. These two steps were designed to be complementary. The segmentation informed the tracking which highlighted issues with the segmentation. Any errors detected through the cell tracking allowed the segmentation to be corrected and improved. The subsequent analysis in TissueMiner required a segmented and tracked cell image sequence which was of sufficiently good accuracy in order to provide reliable and meaningful interpretations. Therefore, in addition to an automated tracker, a set of interactive tools were developed to provide the option to manually correct the segmented image sequences and the results from the tracking.

###### Post-imaging

**Input**: Microscope creates .czi file with Z-stacks in time

Open .czi Zeiss Microscopy Image file in ImageJ/Fiji and export as .btf (BigTif)

**Output**: BigTif: Z-stacks in time

###### Projection

**Input**: Btf file saved in ImageJ/Fiji, full Z-stacks for each time point

Open surface projection programme in Matlab

Use GUI to manually identify the desired projection regions in every 10th frame

Projection programme automatically interpolates projection for the frames in between

**Output**: Tif: single projected slice for each time point

###### Skeletonization

**Input**: Tif: single projected slice for each time point

Using Skeletor, set threshold to between 0.2-0.3

Generates separate .tiff files of individual skeletonized frames

Use ImageJ to concatenate frames

**Output**: Tif: skeletonized image for each time point (Skeleton v1)

###### Manual correction

**Input**: Tif: Skeleton v1, and Tif: single projected slice for each time point

Add Matlab files in Manual Correction folder to Matlab file path

Add projection and Skeleton v1 images to Matlab file path

Run SkeletonStart.m

Load Projection Tif stack

Load Skeleton Tif stack

Manually correct as many missing or extra junctions as possible using hotkeys

Save corrected skeleton

**Output**: Tif: Corrected skeletonized image for each time point (Skeleton v2)

###### Preliminary tracking and further manual correction

**Input**: Tif: Corrected skeletonized image for each time point (Skeleton v2)

Add Matlab files in Tracking Correction folder to Matlab file path

Add projection and Skeleton v2 images to Matlab file path

Run TrackingCorrectionStart.m

Load Projection Tif stack

‘Find cells’, Matlab finds the centroids of uploaded skeleton

‘Tracking’, preliminary tracking identifies points where track is lost due to skeleton errors

‘Problems’ selected from drop-down menu, this will automatically start selecting lost tracks

Manually correct the junction errors identified

Click ‘Export Skeleton’ to save corrected Skeleton v3

Binarize stack in ImageJ

Use ‘SaveAsSingleTIFFs’ Matlab programme to separate into individual tifs

**Output** : Individual Tif files, corrected binarized skeletonized image for each time point

(Skeleton v3)

###### Tissue Analyser skeleton processing

**Input**: Skeleton v3: corrected binarized skeletonized image for each time point

Transfer individual Tifs of Skeleton v3 into Tissue Analyzer

Use the Detect Bonds (Save Watershed) function in Tissue Analyser to export Tissue Miner-compatible skeleton into individual folders

No blur and no removal of cells with x pixels

Use Mac terminal to transfer the individual Tifs into individual folders within the same directory

CODE

~~~
for d in */ ; do (cd “$d” && pwd && cp handCorrection.tif”‥/dv$(basename “$d”).tif”
);done;
~~~

Concatenate individual tifs to a single Tif stack in ImageJ

This creates an RGB Tif stack in time, use ImageJ to create 8-bit image

Check for any new errors that may have been introduced by Tissue Analyser processing by repeating the preliminary tracking correction

Correct errors and repeat Tissue Analyser processing

**Output**: Tif: Corrected skeletonized image for each time point (Skeleton #4)

###### Tracking in Matlab

**Input: Tif file, corrected skeletonized images over time (Skeleton #4)**

Alter AutoTrackingStart.m code to ensure it has the correct filename of skeleton #4 To start tracker:

CODE:

~~~
cd ~/filepath/Tracker/data
export MATLABPATH=/home/ainsli01/Documents/Tracker:/home/ainsli0 1/Documents/Tracker/data
nohup matlab -nodesktop -nodisplay “noFigureWindows -nosplash -r “cd(’/home/ainsli01/Documents/Tracker/data’);AutoTrackingStart; quit” −logfile logfile.out <
/dev/null &
~~~

This will generate three .tif stacks over time, TrackedCellsRGB: each cell colored with a unique cell I.D., DivisionsRGB: divisions highlighted in blue, ErrorsRGB: errors highlighted in red

**Output**: RGB Tif stacks of Tracked Cells, Divisions and Errors

###### Using ErrorRGB output in to correct skeleton

**Input**: Skeleton v4 and ErrorRGB Tif stack

In ImageJ, separate red channel from ErrorRGB Tif stack, and subtract the skeleton resulting in an image with only red cells

Binarize red cells and use ‘Analyze Particles’ in ImageJ to generate list of coordinates and time points of red cells

Open skeleton in Image J, use ‘SpecifyArea’ macro to find errors in list of particles:

IMAGEJ MACRO CODE:

~~~
macro“SpecifyArea [c]”{run(“Specify…”);}
~~~

Enter x,y,t coordinates, automatically takes you to error

Annotate list of errors, label as either: tracking, division, skeleton or edge (edge errors can be ignored)

**Output**: Skeleton #5 and a list of annotated errors

###### If skeleton errors above 50, RETURN TO FIRST STAGE OF MANUAL CORRECTION If skeleton errors below 50, correct skeleton errors and divisions in ImageJ

Correcting missed divisions:

- Using the ‘SpecifyArea’ macro go through Divisions Tif stack finding the errors annotated as ‘Divisions’
- Use ‘pick color’ to select division blue color, and use ‘fill’ function (4-connected) to fill just-divided cells that have been missed

Correcting skeleton:

- Correcting raw skeleton tiff stack using 1-pixel thick paintbrush in ImageJ, taking care to only generate junctions that are 1-pixel thick
- Copy-paste new junction into TrackedCellsRGB and Divisions tiff stacks
- Making sure to correct the colors by using the ‘pick color’ and ‘fill’ function in ImageJ

###### Make Tracker output compatible with Tissue Analyser and Tissue Miner

**Input**: Projected image stack and TrackedCellsRGB and Divisions image stacks

Open ‘SaveAsTissueMiner.m’ and make sure filenames are correct

Run ‘SaveAsTissueMiner’

**Output**: individual folders containing an individual projection, TrackedCellsRGB and division image for each time point

###### Correct Tracks using Tissue Analyser

**Input**: individual folders containing projection, TrackedCellsRGB and division image for each time point, as well a list of tracking errors

Fix broken and swapped tracks using the tools in Tissue Analyser

**Output**: corrected TrackedCellsRGB files

###### Tissue Miner analysis and further Quality Control

**Input**: corrected Tracked Cells and Divisions

Use Tissue Miner to highlight cell I.D.s that disappear in red in final frame visible, these are either apoptoses or tracking and skeleton errors

In ImageJ, separate red channel

Binarize red cells and use ‘Analyze Particles’ in ImageJ to generate list of coordinates and time points of red cells

Save list of red cells as an Excel file

Use ‘SpecifyArea’ macro to find errors and annotate list of particles

Correct errors as described in Steps 11 and 13

**Output**: individual folders containing an individual projection, fully corrected TrackedCellsRGB and division images for each time point

RETURN TO **MANUAL CORRECTION**if there are significant remaining skeleton and tracking errors

*Otherwise, continue Tissue Miner data analysis*

##### Projection

The cell projection tool was developed to provide a semi-automated means of creating cell surface projections of histoblast and LEC image stacks. This was necessary as strong cell signal could often be found above and below the main cell surface which sometimes required manual corrections. The vast data volumes to be processed required an efficient approach with as little manual intervention as possible and a convenient approach to correcting the surface projection where necessary. We carried out the corrections on a sub-sampled image sequence with subsequent spatial and temporal interpolation over the whole sequence. A tool featuring a graphical user interface was developed for this purpose in Matlab.

After sub-sampling in the time domain (typically 1/10 frames) to generate key frames, the maximum intensity projection over the z-depth provided a first estimate where surface markers would be placed at depth levels expressing intensity values in the top. Only a subset of markers was used to keep the number of markers low. These markers could be deleted and added interactively by the user (Tetley et al., 2019). Markers could also be copied to the next frame and translated to correct for a z-shift. Markers would then be used to calculate a depthmap by iteratively averaging marker depth values greater than zero for all image values resulting in a smooth, interpolated depthmap.

The surface for the whole sequence was generated from the depthmaps of key frames by means of linear interpolation in the time and spatial domain. Cell surface intensities were obtained by the max intensity in the vicinity of the interpolated surface from the original 3D image volume for each time frame. The cell surface was stored as a 2D image sequence which was then imported into the image segmentation and skeletonization step.

##### Skeletonization using Skeletor

For the initial segmentation and ground-truthing for the machine-learning segmentation algorithm, we developed a filter-based watershed algorithm in Mathematica which we called Skeletor. Initially, projections were filtered using seven filter kernels, which are as follows: to highlight edges we developed a new convolution matrix which we termed the davisfilter, where the median intensity of a 3×3 pixel kernel was compared to the median intensity of an 11×11 pixel kernel, if the median for the smaller kernel was greater than for the larger kernel then the origin pixel intensity was kept, if not then the median intensity of the 11×11 pixel kernel was subtracted from the origin pixel value. The next filter was a modified salt and pepper filter called saltpepper which aimed to de-noise the images, here the three largest and smallest pixel values of an 11×11 kernel were removed, and the nearest pixel value to the mean of remaining pixels was used as the origin pixel intensity value. The third filter was a contrast enhancing median filter which we called medianfilter, where if the origin pixel intensity was greater than the median for an 11×11 pixel kernel then the origin value was kept, if not then it was replaced with 0. The fourth and fifth filters both calculated the median and the median deviation for an 11×11 pixel kernel, and kept the origin pixel value if it was greater than the median plus the median deviation, if not then filter four, which we termed MADfilter, would subtract the median for the 11×11 kernel to improve contrast; filter five which we termed MADsmoother, would replace the origin pixel value with the median of the 11×11 kernel, smoothing the background. Finally, the sixth and seventh filters were both in-built Mathematica functions which smooth edges, the sixth being an image convolution with a Shen-Castan Matrix function with an exponential radius of 5 pixels and the seventh the CurvatureFlowFilter function with curvature time of 1. The average intensity for each pixel was then calculated from all seven filtered images, and then convolved with a Shen-Castan Matrix with a 2 pixel radius.

Once the image had been filtered, it was segmented using Mathematica’s gradient descent watershed algorithm, where junctions were merged if the minimum boundary height was below a user-defined value, normally in the range of 0.2 – 0.3. The perimeter of each component was then obtained and all the perimeters combined to produce the initial skeleton. To remove erroneous segmentation two tests were performed on each junction. The first test compared intensity between of junctions to the interior of the cell in order to remove ‘phantom’ junctions; specifically, the median plus quartile deviation intensity for the cell interior was measured and set as a threshold, if the median intensity for each junction was below this value it was removed. The second test removed junctions that had a meandering topology; specifically, the length of a straight line between each junction vertexes was divided by the length of each junction and if the value was below 0.75 (i.e. junctions were 25% longer than a straight line) then they were removed. Both of these tests mainly removed erroneous junctions in the LECs with few junctions in the histoblasts being removed.

##### Skeletonization using machine learning

For the movies WT1-3, skeletonization was carried out using Skeletor. For WT4, we used a UNet Neural Network [arXiv:1606.06650v1] trained using WT1-3 data segmented with Skeletor and manually corrected. We used a 3D version of the network treating the time as a z component of input data. Different training conditions, varying the loss functions, optimizer and learning rate, were tested. After training, predictions were obtained from new images, then used in Tissue Analyzer to segment the epithelia.

##### Manual Correction of skeletons – Skeleton Correction Tool

A second interactive tool was developed to verify and correct the result of the skeletonization in an efficient manner. The tool was written in Matlab and featured a graphical user interface and the ability to use a Wacom Tablet to correct missing or extra cell junctions. The tool allows the overlay of original cell surface and skeletonized images, also blending these image layers together. To aid this process, cells could be identified from the skeletonized image and their centroid position was marked on the surface image. Potential segmentation errors could be easily spotted as the markers were off the cell center. In addition, a previous developed automated cell centroid seed tracker (Heller et al., 2016) was incorporated into the tool as a means of quick, preliminary tracking to highlight potential issues caused by segmentation errors. This would support the user to identify and correct such errors.

The tracking results from the offline tracker could also be shown as an additional image layer. Drawing and erasing of cell junctions on the skeletonized image could be performed using a pen on the Wacom Drawing Tablet. Hotkeys allowed the fast switching between different editing modes which contributes to an efficient workflow. The corrected skeleton image sequence was exported for the automated offline tracking step.

In order to quantify the changes at each stage of corrections in Figure S1B, a thickened version of the original Skeletor output skeleton was created was subtracted from the wild type movie post ‘Manual Correction’ and post ‘Manual Tracking Correction’ in ImageJ. The ‘Analyze Particles’ function was used in ImageJ to count the remaining pixels.

##### Automated Tracking procedure – Tracking Tool

The tracking algorithm was designed to fully automatically track cells in the projected skeletonized images. This was achieved in a three-step process and was performed frame by frame for all cells in the sequence. The first iteration tracked cells by identifying suitable matching cell candidates by means of the best-fit of the cell area. Only cells with a good-fit were included in this sequence with the aim is to get a first estimate of the cell surface motion. In a second step, the motion (flow) field of the whole cell surface was calculated through the interpolation of individual movement vectors from the initial tracking results. In the third step the tracking is guided by the flow estimates. Finally, the tracked cell sequence was further processed to determine the cell lineages and divisions and exported as a number of different image sequences. The following paragraphs describe these steps in more detail:

###### 1. Find initial tracks

The initial tracking is performed by identifying potential matching candidates in the next image frame. The area of each cell in a frame is obtained from the labelled binary skeleton image. Any cell in the next frame that slightly overlaps with the source cell area was a potential candidate. For each target candidate, the cell areas were aligned by centroid position. Similarities in terms of the overlapping cell area were used to determine a good fit. The confidence *c* (a metric for a good-fit) was calculated for each candidate as the ratio between the overlapping aligned cell area and the total combined area of source and target cells.

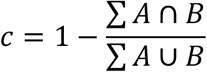

This would exclude fast moving smaller cells that were not overlapping which made the flow guided tracking step necessary.

Once a list of candidates and their confidences was established, the best candidate for each source cell was identified amongst the several potential target candidates. Each source cell was assigned the target cell with the highest confidence first. This is performed iteratively for all source cells for each iteration starting with the highest confidences first and iterating down as long as the confidences are above the minimum threshold of 0.7. The source cells were thus competing for target cell candidates as neighboring cells could have the same potential target candidates which ensured that the best candidate was assigned for each source cell. It is important to note that in the first iteration of the tracking we only used cells with high confidences ≧0.7, i.e. cells which we could be confident about to be correct. In a final step new track IDs were assigned to cells which were not paired due to having a low confidence value.

This first tracking step returned an image matrix consisting of 16-bit grayscale images. The color zero (black) denoted the cell boundaries. Each cell area had a uniform numerical value which denoted the track ID. The track ID remained the same for matching cells in subsequent frames. New cells or broken tracks received with a new track ID.

###### 2. Calculate the flow field estimate

In this step an interpolated flow field from the cell movements of the previous tracking step was calculated. The displacement vector for each cell was calculated from the centroid positions of the tracked cells. The flow field was calculated for each position in the image matrix by a weighted average of all the displacement vectors with a weighting factor which was the inverse of the distance to all the other cell positions in the frame. The influence of nearby cells is thus much greater than cells at greater distances. The interpolation was completed when all the values in the image matrix had been calculated. This step returned the interpolated flow field for each frame in the sequence.

###### 3. Re-track all cells by using the flow estimate

The final cell tracking was performed by taking into account the flow estimates of cells. As in the first tracking step, any cell in the next frame that slightly overlapped with the source cell area was a potential candidate. However, in this step, the source cell area was translated by the average flow field displacement of that area obtained from the initial tracking. The overlap between the translated area of source and target cells was then calculated as in step 1 above.

The assignment process was more complex as in the first tracking iteration. The aim was to find the best candidate for each source cell amongst the several potential target candidates. However, this was performed over several steps as all the source cells needed to be paired, not just the most obvious fits. During the first step, each source cell was assigned the candidate cell with the highest confidence first. This was run competitively for all source cells for each iteration starting with the highest confidences first and iterating down as long as the confidences were above the minimum threshold of 0.2. The source cells were thus competing for target cell candidates as neighboring cells could have the same potential target candidates which ensures that the best candidate was assigned for each source cell.

In a second step, the above process was repeated for not yet assigned source cells by going through the remaining candidates. In a third iteration, the not yet assigned source cells were assigned by distance up to a max distance to target cells.

Finally, any unassigned cells were treated as new cells which were assigned a new cellID and denoting the start of a new track sequence.

This second tracking step returned an image matrix consisting of 16-bit grayscale images. The color zero (black) denoted the cell boundaries. Each cell area had a uniform numerical value denoting the track ID. The track ID remained the same for a matching cell in subsequent frames. New cells or broken tracks received with a new track ID.

###### 4. Exporting tracking result

The track sequence was exported as an RGB image sequence whereby each cell track was given a unique RGB color. Divisions were identified by a sudden increase in cell size in a tracked sequence which also coincided with a new cell track emerging in a subsequent frame in its locality. The ratio of the cell size change between frames was used a measure to identify divisions. A cell which underwent such transformation was identified as the mother cell while a new cell it is vicinity was labelled as the daughter cell. From these divisions, a lineage sequence could be created, highlighting cell divisions in different shades of blue. Potential errors in the lineage emerged when new cells did not originate from a division event or when the lineage was not clear. Cells with such issues were labelled in red in an additional ‘Error’ sequence and exported for further manual inspection and correction if deemed necessary.

###### 5. Manual Correction of Tracking

The output from the offline tracker, including the lineage and error sequences were imported into ImageJ for further inspection and correction. Cells highlighted in red ‘Error’, i.e., cells with a new cell ID that do not originate from a division were labelled manually using the ‘Analyze Particles’ function in ImageJ. This creates a list of red cells that are manually checked and annotated according to the type of error, either a ‘Tracking’ error due to the cell migrating quickly, or a ‘Division’ error where Tracker has failed to pick up a division, or a ‘Skeleton’ error that remains. Using this information, each type of error was corrected in a different way. If the number of remaining skeleton errors was significant (above 50), then only the input skeleton would be corrected, and the automatic tracking process would start again. If the number of remaining skeleton errors was below 50, then all the errors would be corrected manually using the coordinates lifted from the error layer. First the ‘Division’ errors were corrected in the division layer using the ‘fill’ function in ImageJ to fill daughter cells of missed divisions in the correct shade of blue. Then the ‘Skeleton’ errors were corrected in both the division layer and the unique cell ID. layer using ImageJ. White lines were added or removed manually in the division channel in ImageJ, taking care to ensure all lines were maintained at a 1-pixel thickness. Then the affected area was copied and pasted into the unique cell ID. channel to ensure the white lines were the same. Using the fill function ensured the colors in the unique cell ID. channel were unaffected. Finally, the ‘Tracking’ errors were corrected in Tissue Analyser (Aigouy et al., 2010; Etournay et al., 2016). Once the affected coordinates were found, it was possible to swap tracks around, join truncated tracks or create new cell IDs wherever necessary. The corrected lineage sequence was finally converted and exported into a format that was readable by Tissue Miner.

##### Tracking validation and quality control

All sequences went through a thorough process of quality control. The tracker supported this approach by highlighting potential issues in an ‘error’ sequence. Once the skeleton errors were corrected, any subsequent errors were due to tracking errors and cell division errors. A typical sequence featured between five and six thousand cell tracks. The tracker would typically highlight a total of around 300 (5-6%) of these tracks as having potential issues which required inspection: Approx. 1-2% of the tracks had tracking errors whereby the tracker did not correctly identify a matching cell. These could be amended by linking up broken tracks or swapping tracks. However, most issues, approx. 3-4% of all the tracks arose at or in the vicinity of dividing cells when mother or daughter cells were not identified, swapped or not classified as being part of the lineage. Not all these issues were genuine errors that required correction, but all these issues required inspection by an operator to ensure a high level of quality control.

##### Tissue Miner Analysis

Once these correction steps have been completed using ImageJ and Tissue Analyser, Tissue Miner was then used to extract data from the tracked cells (Etournay et al., 2016; Etournay et al., 2015); see Supplementary Theory for details. Tissue Miner was also used to identify any remaining errors by detecting loss of cell I.D., as well as detecting with an anomalously short ‘cycle time’ in between divisions. Any errors that were detected using these criteria were corrected in the manner described under ‘Manual Correction of Tracking’, and then the final data is re-entered into Tissue Miner.

##### Temporal alignment of WT movies

A salient feature of abdomen development is the formation of the sensory organs of the adult abdomen, which are mechanosensory bristles that arise through multiple stereotypical rounds of asymmetric cell divisions from a single progenitor, the Sensory Organ Precursor (SOP) (Fabre et al., 2008). We tracked the SOP lineages and excluded them from all subsequent analysis on histoblast growth and proliferation (Figure S1F). However, we noticed that the emergence of SOPs over time, identified by the initial asymmetric division, followed a sigmoidal temporal distribution that could readily be fitted by a Hill function (Figure S1G-K). To allow comparison of the growth parameters, these curves were used to temporally align our wild type movies (WT1-4) (Figure S1K).

#### QUANTIFICATION AND STATISTICAL ANALYSIS

All statistical tests were performed using R or Graphpad Prism. Statistical tests used and number of repeats are indicated in the figure legends or in the text.

#### KEY RESOURCES TABLE

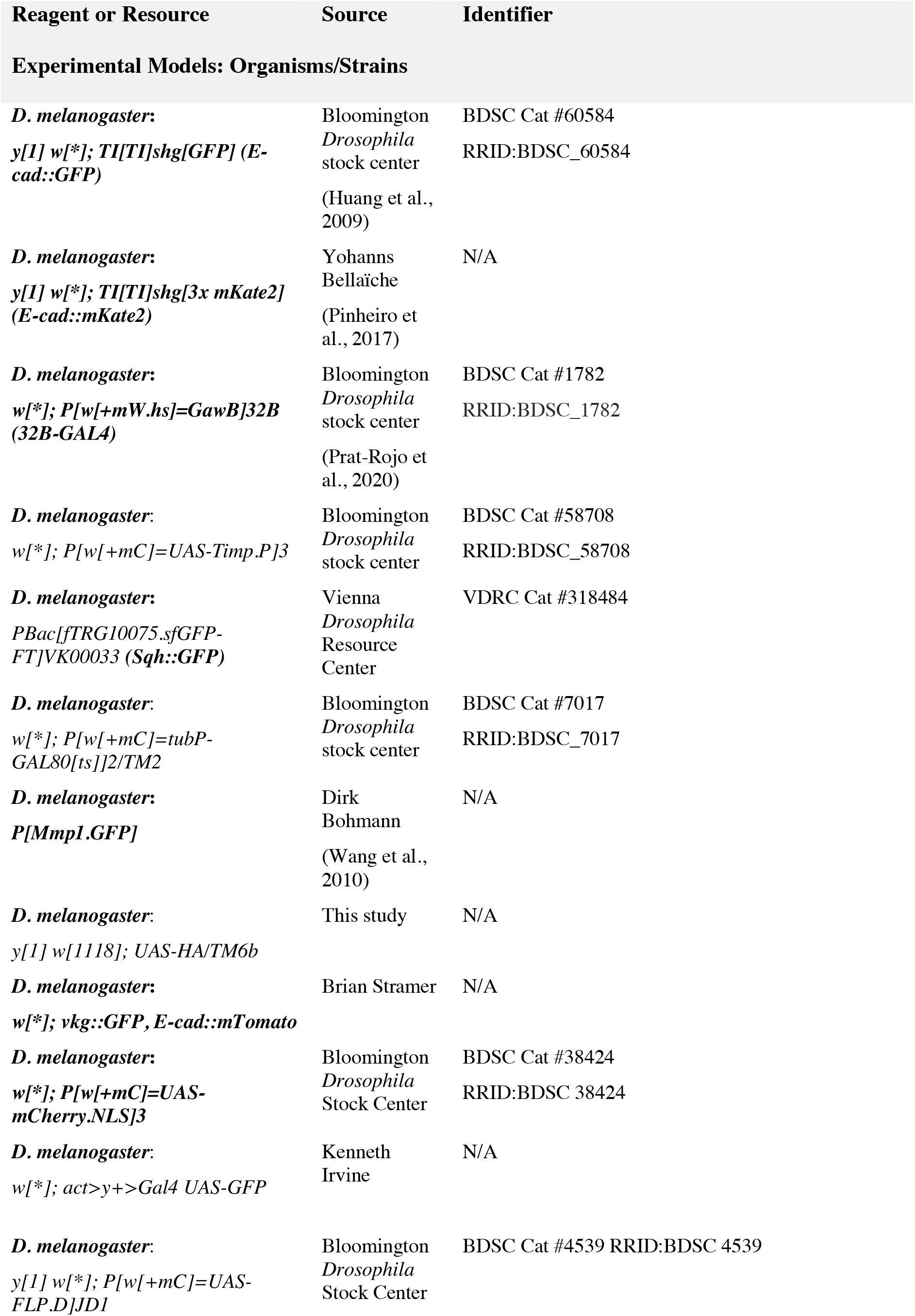
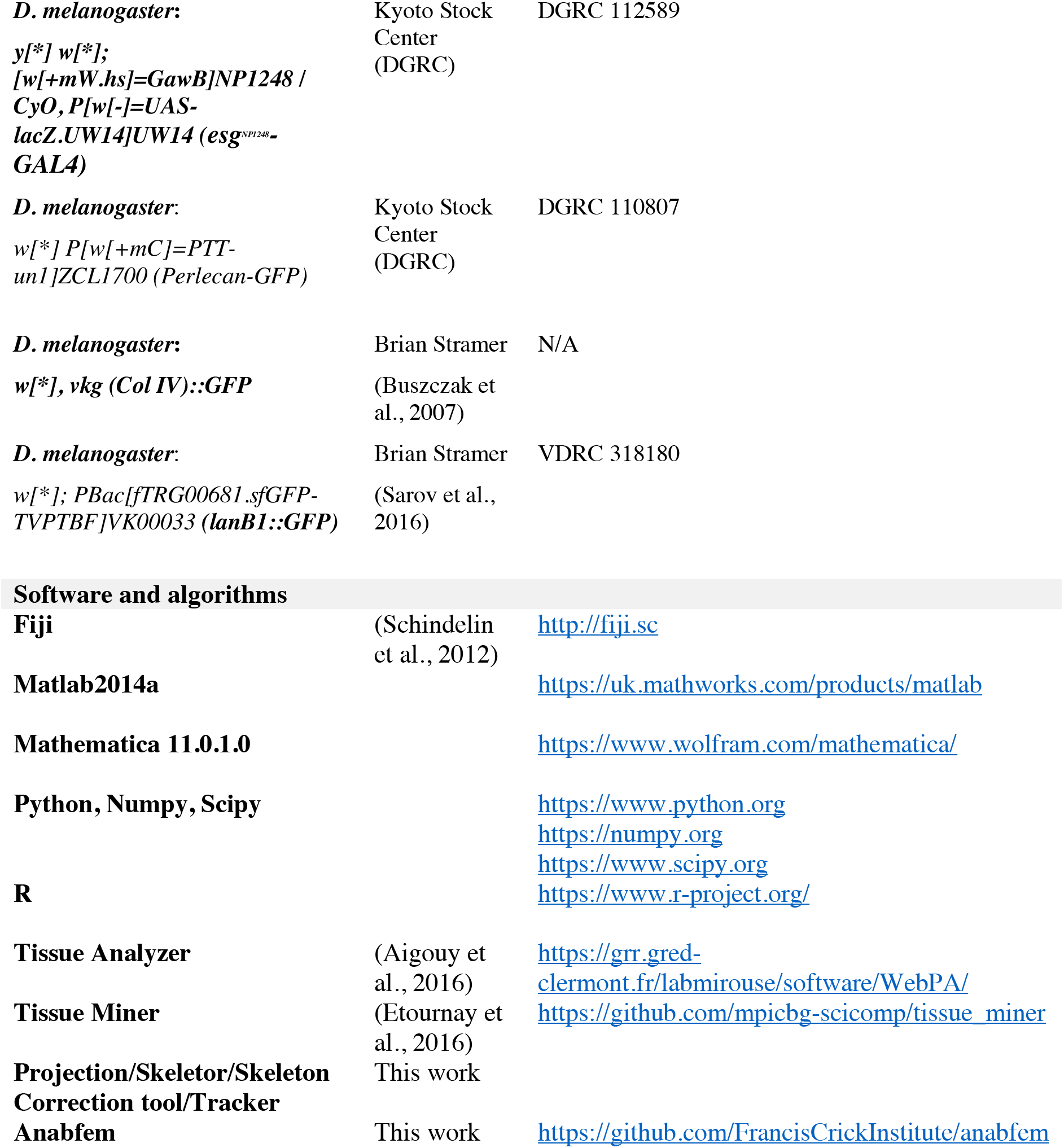

