## Supplemental Information for "ECM remodeling and spatial cell cycle coordination determine tissue growth kinetics"

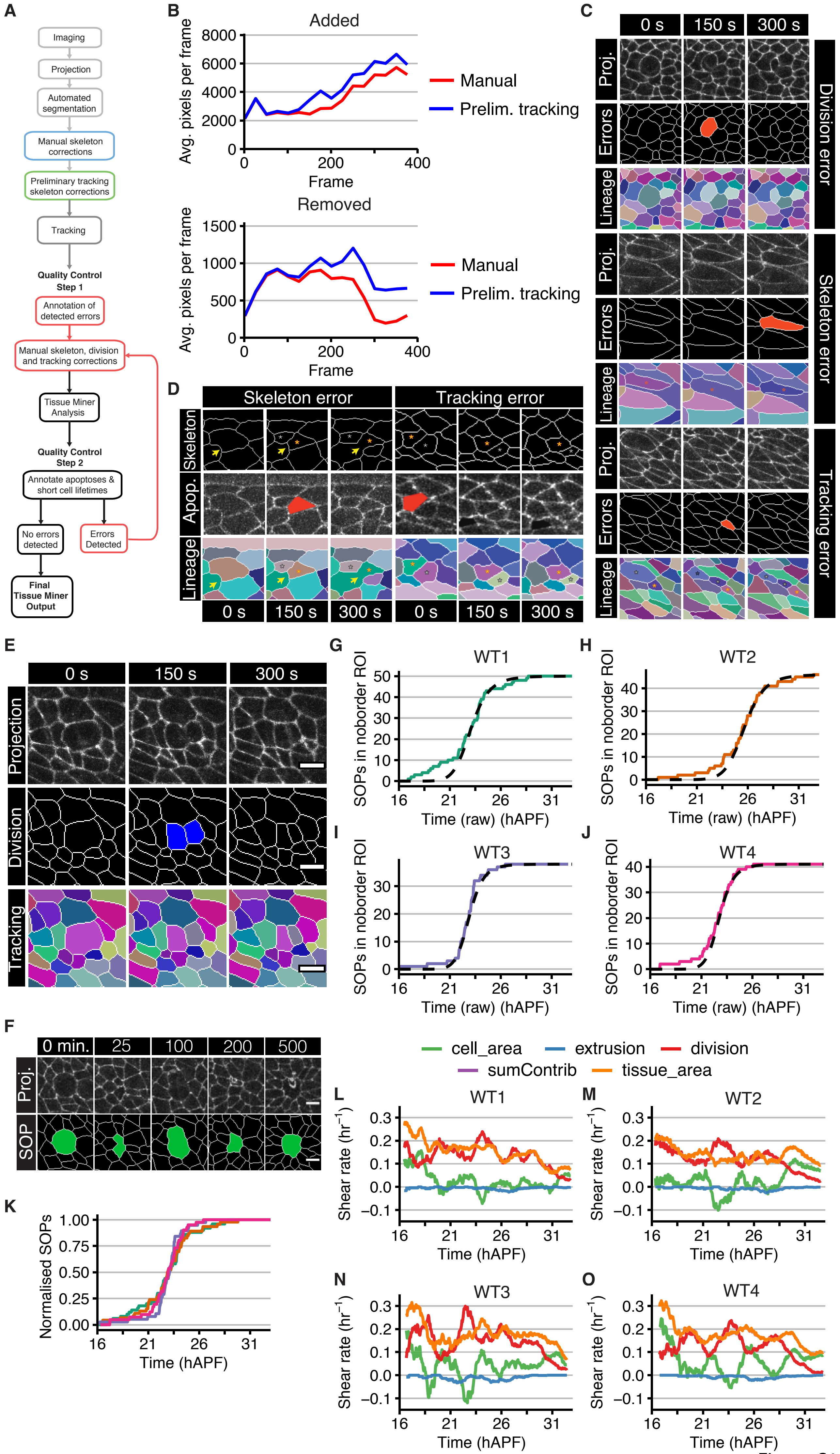
A
Imaging
Projection
Automated segmentation
Manual skeleton corrections
Preliminary tracking skeleton corrections
Tracking
Quality Control Step 1
Annotation of detected errors
Manual skeleton, division and tracking corrections
Tissue Miner Analysis
Quality Control Step 2
Annotate apoptoses & short cell lifetimes
No errors detected
Errors Detected
Final Tissue Miner Output
B
Added
Avg. pixels per frame
Frame
Manual
Prelim. tracking
Removed
C
0 s
150 s
300 s
Proj.
Errors
Lineage
Division error
Skeleton error
Tracking error
D
Skeleton error
Tracking error
Skeleton
Apop.
Lineage
E
Projection
Division
Tracking
F
0 min.
25
100
200
500
Proj.
SOP
G
WT1
SOPs in noborder ROI
Time (raw) (hAPF)
H
WT2
I
WT3
J
WT4
K
Normalised SOPs
Time (hAPF)
cell_area
extrusion
division
sumContrib
tissue_area
L
M
N
O
Shear rate (hr⁻¹)

Figure S1

**Figure S1. Segmentation, tracking and temporal alignment of histoblast movies**

(A) Overview of the pipeline including imaging, skeletonization, correction steps, tracking, quality control steps, and final output for analysis (see STAR Methods for details).

(B) Number of pixels added or removed during the process of manual corrections and tracking corrections in WT3. Values generated by subtracting the original skeleton from the corrected skeletons. Numbers are calculated over 25 frames.

(C) Examples of error types detected by Tracker. Top, stills from a movie of a live pupal abdomen expressing E-cad::GFP after undergoing contour projection. Middle, error output generated by Tracker, with errors labelled in red. Bottom, lineage output from Tracker with each cell labelled with a unique color. Red and black stars indicate the location of the same cell and daughter cells of the same lineage between frames. Division error: red cell occurs because division has not been accurately detected by Tracker, thus the new cell is detected as an error. Skeleton Error: red cell occurs because of missing junction. When junction returns at 300 s, the cell has a new I.D., which is labelled as an error. Tracking error: red cell occurs due to rapid cell migration and cell division. New daughter cell from division is mistakenly labelled as a pre-existing cell at 150 s (black star, dark purple cell), and pre-existing cell is mistakenly labelled with a new cell I.D. at 150 s (red star, pink cell), which is labelled as an error.

(D) Examples of supposed cell losses (loss of cell I.D) detected by Tissue Miner which are actually skeleton and tracking errors. Top, skeleton analyzed by Tissue Miner. Middle, stills from a movie of a pupa expressing E-cad::GFP after undergoing contour projection, with the lost cell labelled in red by Tissue Miner. Bottom, lineage output from Tracker with each cell labelled with a unique color. Yellow arrows point towards the missing junction. Red and black stars indicate the same cell between frames. Skeleton Error: Missing junction labelled with yellow arrow at 300 s means that the final frame for the brown cell is at 150 s, thus it is labelled as a cell loss event at this time. Tracking Error: Cells are migrating rapidly and Tracker mislabels the green cell as the purple cell at 150 s. The green cell is labelled as an imminent apoptosis at the final frame where it is visible at 0 s.

(E) Tracking and cell-division masks for input into the Tissue Miner software. Top row: example movie frames showing a cell division. Middle row: cell division mask of the same frames. In the first frame after the division, the newly-produced daughter cells are marked blue. Bottom row: tracking mask, in which each cell is marked with a unique color that persists over time. To match the convention Tissue Miner expects, one daughter cell retains the color of the mother, but will be parsed as a new cell by Tissue Miner. Scale bar=5 $\mu$m.

(F) Top: Stills from a movie of a pupa expressing E-cad::GFP, following a sensory organ precursor cell (SOP) undergoing differentiation. Bottom: Skeletonization of SOP as a single cell (green-filled) from the moment of first asymmetric division to simplify the segmentation and tracking process. Scale bar = 5 $\mu$m.

(G-K) Time-alignment method for multiple WT movies. (K) normalized count of the appearance of sensory organ precursors (SOPs) in the noborder ROI over time. Each movie is time-aligned relative to WT1 (whose first frame is set to be at 16 hAPF), by an offset determined by fitting Hill functions to the individual movie data shown in (G-J).

(L-O) Contributions to area expansion rate of the noborder ROI from cell area relative rate of change, cell division rate, cell extrusion rate. The sum of these 3 contributions matches the directly measured area expansion rate. Integration of the shear rate leads to the cumulative contributions, e.g. as for WT1 in Figure 1C.

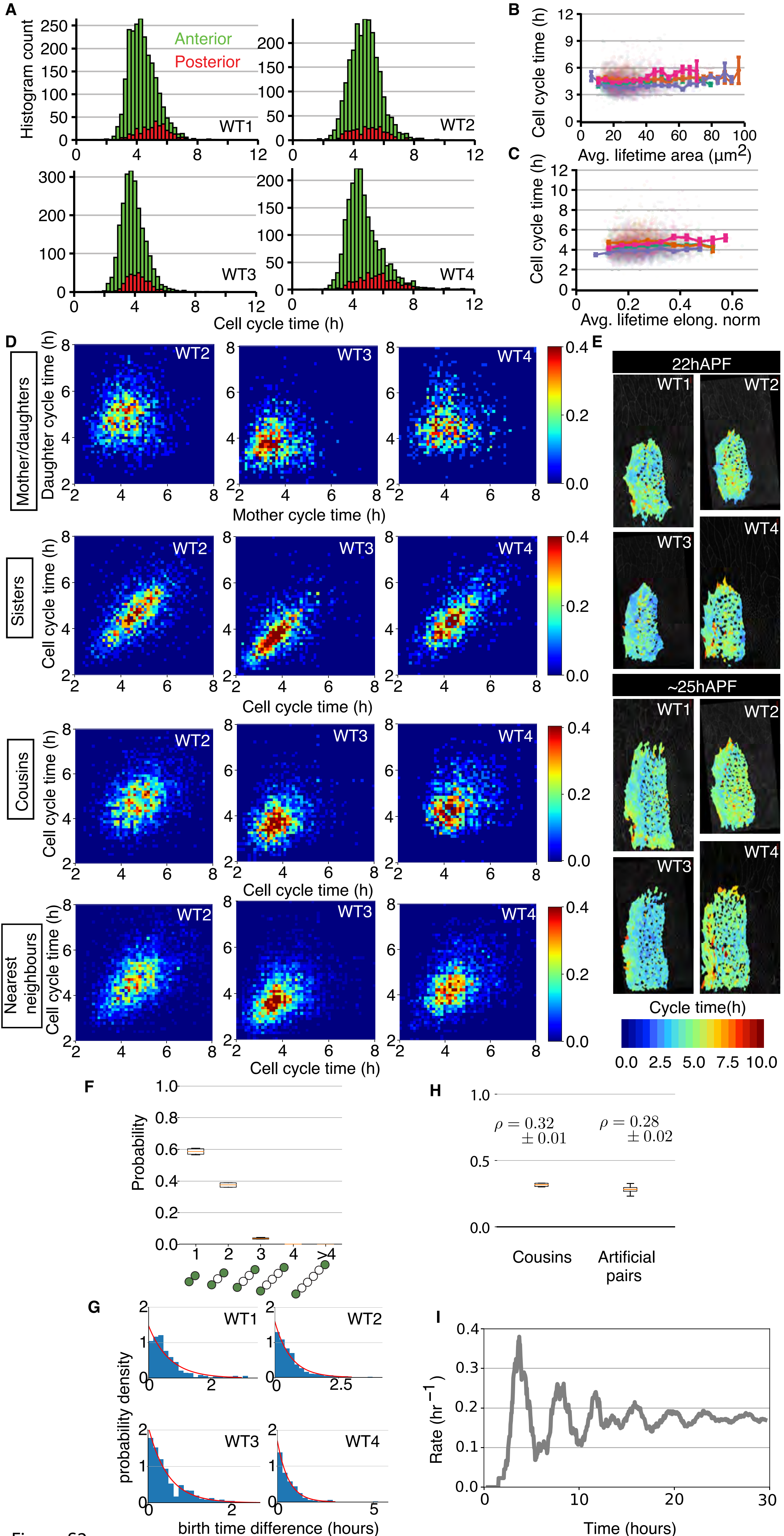
A
Histogram count
Anterior
Posterior
WT1
WT2
WT3
WT4
Cell cycle time (h)
B
Cell cycle time (h)
Avg. lifetime area (μm2)
C
Cell cycle time (h)
Avg. lifetime elong. norm
D
Mother/daughters
Daughter cycle time (h)
Mother cycle time (h)
Sisters
Cell cycle time (h)
Cousins
Nearest neighbours
E
22hAPF
~25hAPF
Cycle time(h)
0.0 2.5 5.0 7.5 10.0
F
Probability
H
ρ = 0.32 ± 0.01
ρ = 0.28 ± 0.02
Cousins
Artificial pairs
G
probability density
birth time difference (hours)
I
Rate (hr−1)
Time (hours)

Figure S2

**Figure S2. Cell cycle time analyses during abdominal development**

(A) Histograms of cell cycle times (defined only for cells that are observed appearing and disappearing via division events) in the anterior and posterior nests in WT1—WT4.

(B) Cell cycle time as a function of the cell apical area, averaged over the lifetime of the cell. Solid lines: binned data. Error bars: SEM for the bin. Transparent dots: individual data points. Color code for different WTs is as in Figure 2.

(C) Cell cycle time as a function of the cell elongation norm, averaged over the lifetime of the cell. Data presentation is as in panel B.

(D) Probability density of pairs of cell cycle times, where pairs are taken between, from top to bottom, mother-daughters, sisters, cousins, and nearest-neighbors whose birth times differ by less than 0.5 hours, excluding sisters. Experimental data from movies WT2-4 (see Figure 2D for WT1). Cells are taken from the visible anterior nest. High densities of pair along the diagonal indicate high levels of correlation in the pair of cell cycle times.

(E) Snapshots from each WT movie with histoblast cells colored according to their cycle time at two intermediate times 22 hAPF and ~25 hAPF. Uncolored (black) cells are SOPs, arrested cells, cells which cannot be tracked from birth to mitosis, or cells with a cycle time larger than 10 hours.

(F) Probability of distance relation between cousin cells (see Supplementary theory for a definition of distance relation). Most cousin cells are nearest or second-nearest neighbors.

(G) Probability distribution of birth time difference of cousin cells (blue bars). Red line, exponential fit to the probability distribution.

(H) Comparison between correlation coefficient of cousin cells, and artificially constructed cell pairs with the same neighbor relations (as in F) and distribution of birth time difference (as in G), excluding sisters and cousins. The correlation coefficient of artificially constructed pairs is close to the correlation coefficient of cousin cells, indicating that cousin correlations are largely explained by their spatial relations. Right, distribution obtained for 10 random realizations of artificial pairs for the 4 WTs. Spearman correlation coefficients: $\rho_S$=0.33±0.03 (cousins), $\rho_S$=0.31±0.02 (artificial pairs).

(I) Cell division rate for one realization of a simulation of a cell population growing during 30 hours. Cell cycle times are taken from a normal probability distribution with mean 4 hours and coefficient of variation 0.2, roughly similar to the experimental distribution in A (see Supplementary Theory for details). At t=0 h the simulation starts with 100 cells, at the beginning of their cell cycle. The initial cell synchrony results in damped oscillations in the cell division rate. A moving average is applied as in Figure 2A.

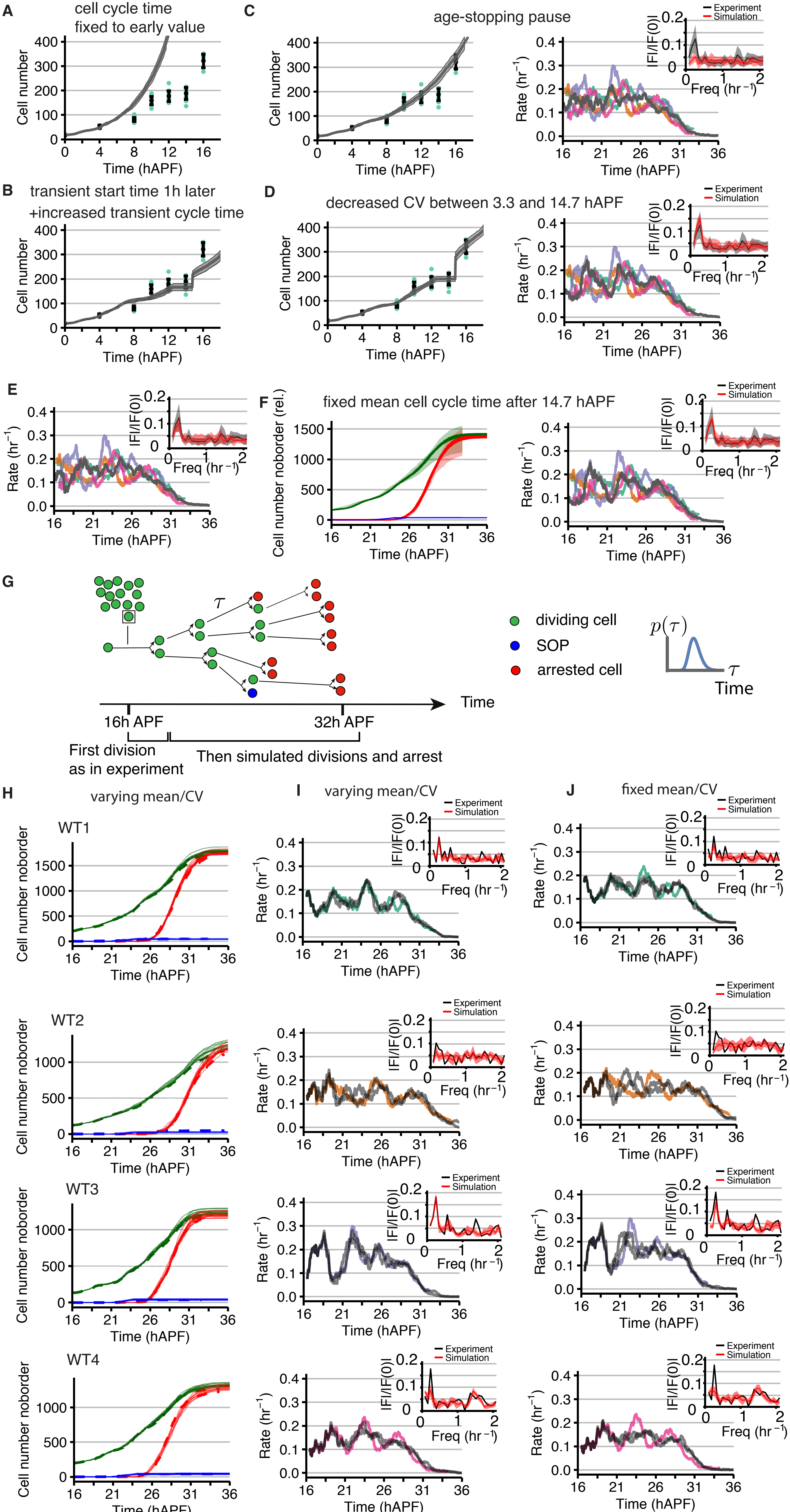
A
cell cycle time
fixed to early value
Cell number
Time (hAPF)
B
transient start time 1h later
+increased transient cycle time
C
age-stopping pause
D
decreased CV between 3.3 and 14.7 hAPF
E
F
fixed mean cell cycle time after 14.7 hAPF
Cell number noborder (rel.)
Rate (hr$^{-1}$)
|FI/|F(0)|
Freq (hr$^{-1}$)
Experiment
Simulation
G
τ
dividing cell
SOP
arrested cell
p(τ)
Time
16h APF
32h APF
First division
as in experiment
Then simulated divisions and arrest
H
varying mean/CV
WT1
WT2
WT3
WT4
Cell number noborder
I
varying mean/CV
J
fixed mean/CV

Figure S3

**Figure S3. Simulations of histoblast proliferation kinetics**

(A) Number of cells before 16 hAPF, experiment and simulations. Simulations are as in Figure 3E-H but the mean cycle time is set to be constant and equal to the value measured experimentally before 3 hAPF. The cell number increases much too quickly.

(B) Number of cells before 16 hAPF, experiment and simulations. Simulation results are as in Figure 3E-H, but the mean cell cycle time increases to 6.6 h at 4.3 hAPF, instead of increasing to 4.6 h at 3.3 hAPF. This choice leads to a roughly correct number of cells produced around 14hAPF, but agreement with the measured number of cells over time appears overall less good than in Figure 3E.

(C) Simulation result with an alternative implementation of the pause period around 12 hAPF. (Left) Number of cells in the histoblast before 16 hAPF, as in Figure 3E; (Right) Division rate as a function of time after 16 hAPF, color scheme as in Figure 3H and moving average is applied as in Figure 2A; inset, normalized absolute value of the Fourier transform of the division rate. Whereas in the base case divisions were prevented but cell aging continued, here cell aging is paused from 12.2 hAPF and is restarted at 12.7 hours. In this implementation divisions plateau for a short period around 12 hAPF, but the cell number after the pause increases more gradually. (Right). The age-stopping pause does not lead to the sudden burst of simultaneous divisions shown in Figure 3E and therefore the "re-synchronization", due to setting a large number of cell ages to zero simultaneously, does not occur.

(D) (Left) Number of cells in the histoblast before 16 hAPF, as in Figure 3E; (Right) Division rate as a function of time after 16 hAPF, color scheme as in Figure 3H and moving average is applied as in Figure 2A; inset, normalized absolute value of the Fourier transform of the division rate. Simulation results are as in Figure 3E-H but the coefficient of variation of cell cycle time between 3.3 hAPF and 14.7 hAPF has been set to 0.2 instead of 0.32. This results in oscillations in cell division rate after 16 hAPF which are slightly too large.

(E) Division rate as a function of time after 16 hAPF, color scheme as in Figure 3H and moving average is applied as in Figure 2A; inset, normalized absolute value of the Fourier transform of the division rate. Simulations as in Figure 3H, but results are obtained with the pause time delayed from 12.5 hAPF to 13.5 hAPF; resulting in less pronounced oscillations.

(F) (Left) Number of cells after 16 hAPF, as in Figure 3G; (Right) Division rate as a function of time after 16 hAPF, color scheme as in Figure 3H and moving average is applied as in Figure 2A; inset, normalized absolute value of the Fourier transform of the division rate. Simulation results are as in Figure 3G-H but the mean cycle time in the main phase is constant at 4.42 h, a value chosen by calculating the average (weighted by number of cells) of the binned cycle time data from Figure 3D. There are only subtle differences from the base case Figure 3H, for example that the second division rate peak is slightly lower.

(G) Schematic for the "short" simulations after 16 hAPF. Each cell in the noborder region at 16 hAPF undergoes its first cell division as in experiments. Subsequent cell divisions are performed according to cell cycle times taken from a statistical distribution. Newly born cells become arrested or SOPs according to a time-evolving probability distribution, as described in Supplementary Theory.

(H) Comparison of experimental (dashed) cell number with ten statistical runs (thin lines) of the short simulation, which is initialized at 16 hAPF for each individual WT movie. Color scheme as in Figure 3G.

(I) Experimental (color) and simulation (grey) division rate series for the short simulations from H. Two statistical runs of each simulation are shown, to retain clarity. Inset: Normalized absolute value of the Fourier transform of the division rate for experiment (black) and simulation (red, ribbon showing SD), allowing the magnitude of division rate peaks to be compared.

(J) As H but with an alternative version of the short simulation in which the mean cycle time does not track the experimental data of Figure 3D, but instead is fixed at a value chosen as the weighted time-average of the corresponding experiment's mean cycle time in 25-frame bins. The CV is also set at a fixed value in the same way. The magnitude of the simulation's division rate peaks is typically noticeably less than in panels I.

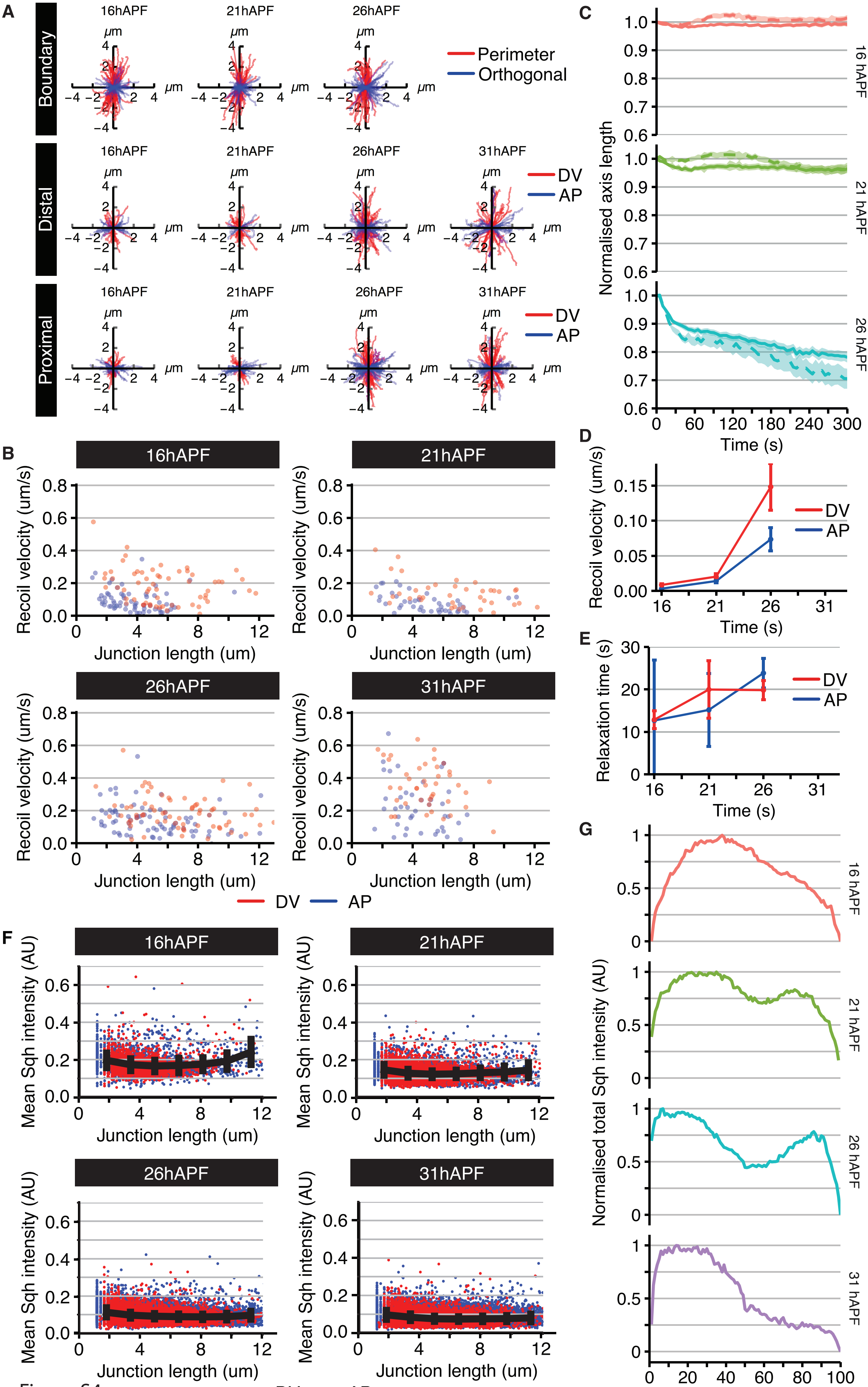

A
Boundary
Distal
Proximal
16hAPF
21hAPF
26hAPF
31hAPF
μm
Perimeter
Orthogonal
DV
AP
B
Recoil velocity (um/s)
Junction length (um)
C
Normalised axis length
Time (s)
16 hAPF
21 hAPF
26 hAPF
D
E
Relaxation time (s)
F
Mean Sqh intensity (AU)
G
Normalised total Sqh intensity (AU)
Ventral to Dorsal (%)
31 hAPF


Figure S4

**Figure S4. Histoblast junction strain increases through development and is independent of junction length or Myo II intensity**

(A) Trajectories of vertices after single junction ablations, relative to their initial positions, in the different populations of histoblasts (see Figure 4B) throughout development. Note that for boundary cells, the trajectories of vertices shared with LECs are on the left and experience reduced displacement compared to the vertex within the nest.

(B) Quantification of junction recoil velocity as a function of junction length throughout development reveals no clear correlation.

(C) Normalized length of the short (oriented along the AP axis, dashed) and long (oriented along the DV axis, solid) axes in LECs after annular ablation.

(D, E) Quantification of recoil velocity (D) and relaxation time (E) along the short (oriented along the AP axis) and long (oriented along the DV axis) axes in LECs after annular ablations.

(F) Sqh::GFP (Myo II) intensity as a function of junction length. Dots are individual data points (red: DV oriented junctions, blue: AP oriented junctions), black line and error bars: binned mean and SD.

(G) Normalized Myo II intensity along the DV axis, across the anterior histoblast nest. Note the dip in intensity at the dorsal side.

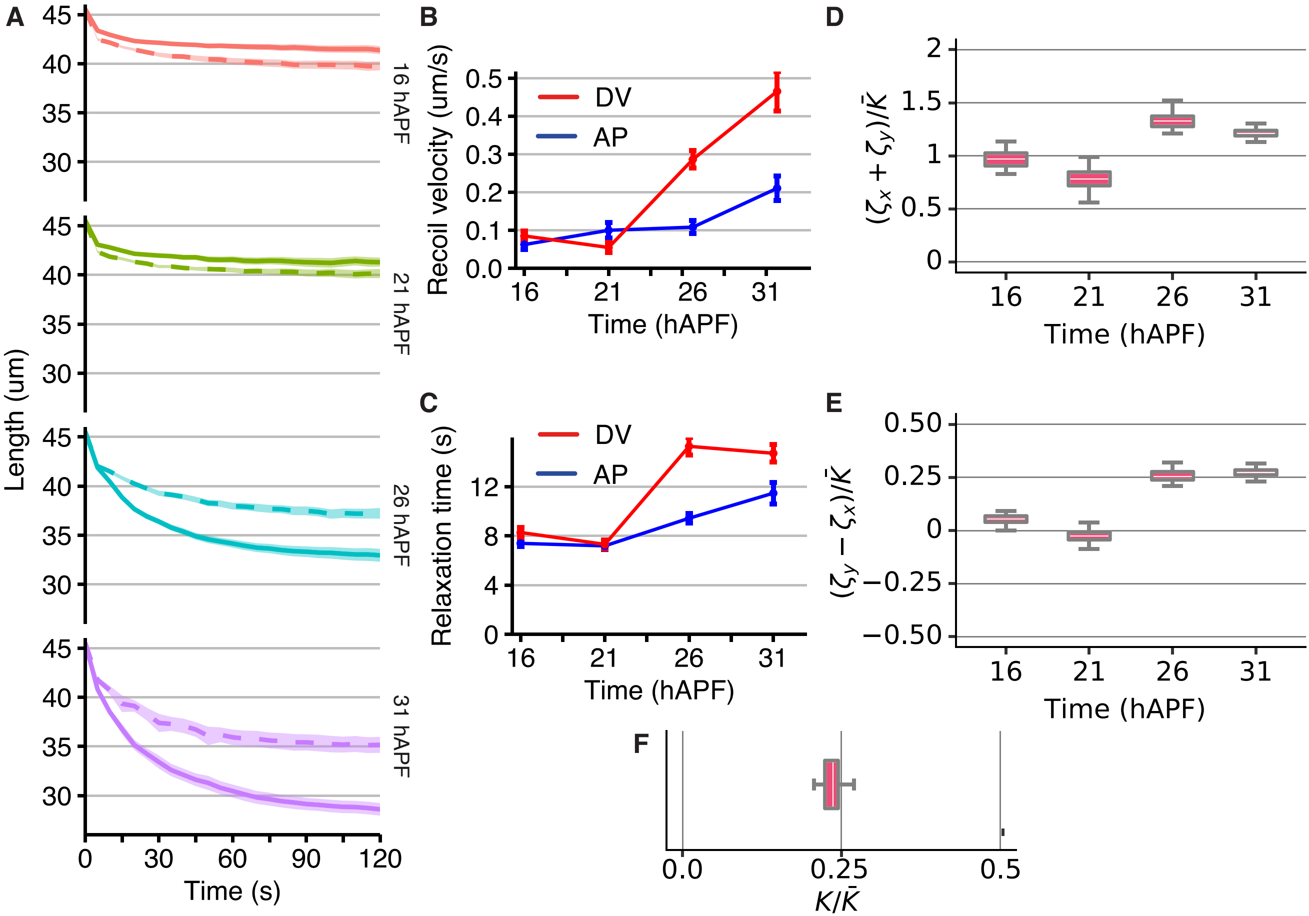
A
B
C
D
E
F
Length (um)
Time (s)
16 hAPF
21 hAPF
26 hAPF
31 hAPF
Recoil velocity (um/s)
Relaxation time (s)
DV
AP
Time (hAPF)
$(\zeta_x + \zeta_y)/\bar{K}$
$(\zeta_y - \zeta_x)/\bar{K}$
$K/\bar{K}$

Figure S5

**Figure S5. Analysis of annular ablation experiments through histoblast development**

(A) Length of AP (dashed) and DV (solid) axes in excised discs after annular ablation.

(B, C) Quantification of recoil velocity (B) and relaxation time (C) along the DV and AP axes of excised histoblasts after annular ablation, throughout development.

(D, E) Normalized isotropic (sum of AP and DV tensions, $\zeta_x + \zeta_y$, D) and anisotropic (difference of DV and AP tensions, $\zeta_y$-$\zeta_x$, E) tissue tensions.

(F) Ratio of tissue shear to elastic modulus, obtained from a fit describing the tissue as an elastic material under tension, to measured excised discs deformation following laser ablation (see Supplementary Theory).

Figure S6

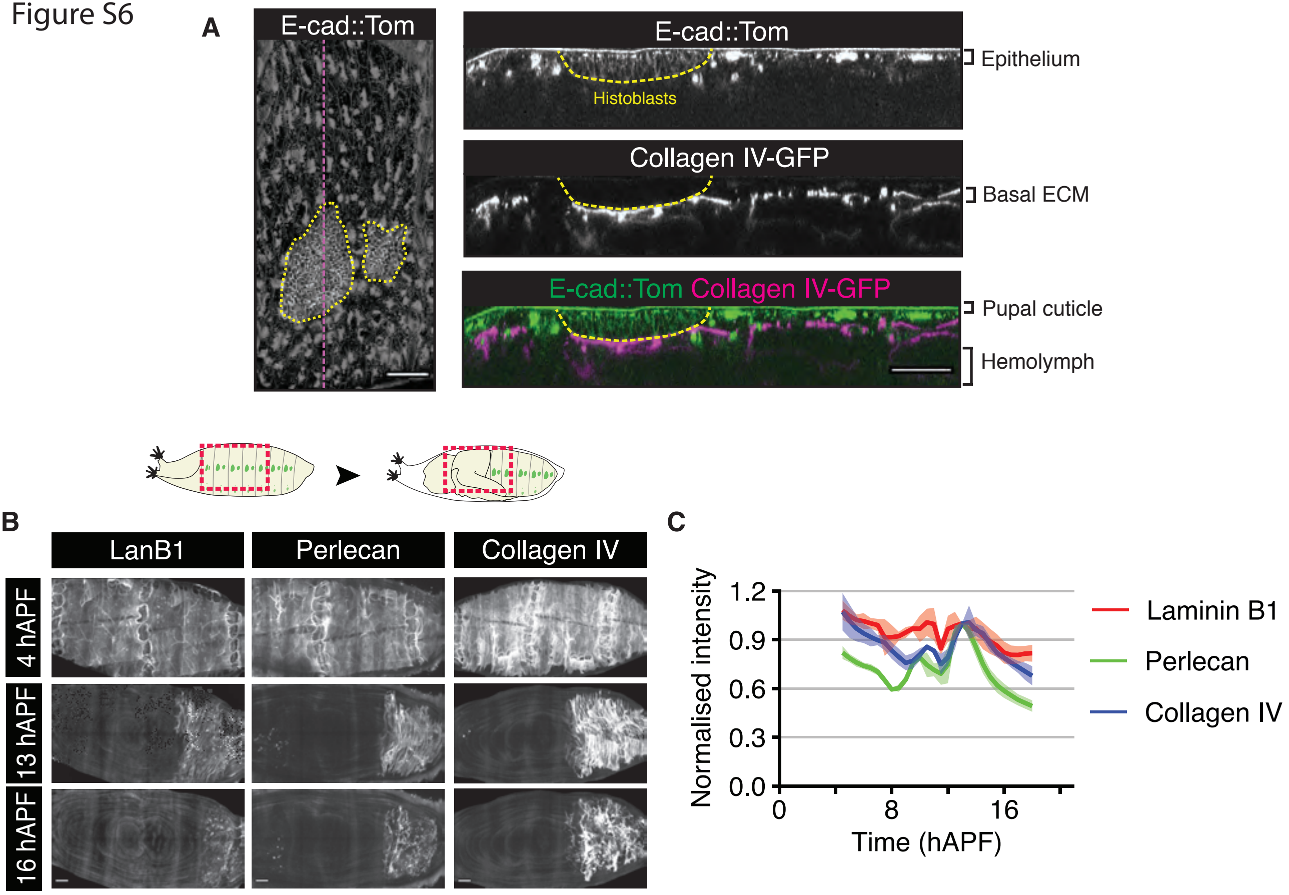
A
E-cad::Tom
E-cad::Tom
Histoblasts
Epithelium
Collagen IV-GFP
Basal ECM
E-cad::Tom Collagen IV-GFP
Pupal cuticle
Hemolymph
B
LanB1
Perlecan
Collagen IV
4 hAPF
13 hAPF
16 hAPF
C
Normalised intensity
1.2
0.9
0.6
0.3
0.0
0
8
16
Time (hAPF)
Laminin B1
Perlecan
Collagen IV

**Figure S6. Dynamics of ECM remodeling.**

(A) Snapshot of a pupa expressing E-cad::Tom and Vkg::GFP (Collagen IV) at 16 hAPF. Yellow dotted lines outline histoblasts. Magenta dotted line indicates position of orthogonal view. Scale bar = 50 $\mu$m. Left panel: Maximum projection of E-cad::Tom. Right panels: orthogonal projections.

(B) Snapshots for the major basal ECM components during the pre-pupal to early pupal stages (schematic above highlights imaging region in red). The head everts at 12 hAPF, pushing the abdomen posteriorly and is complete by 13hAPF.

(C) Quantification of ECM components intensity during early pupal development in the region where the abdomen will reside following head eversion. Measurements were normalized to intensity at 13 hAPF.

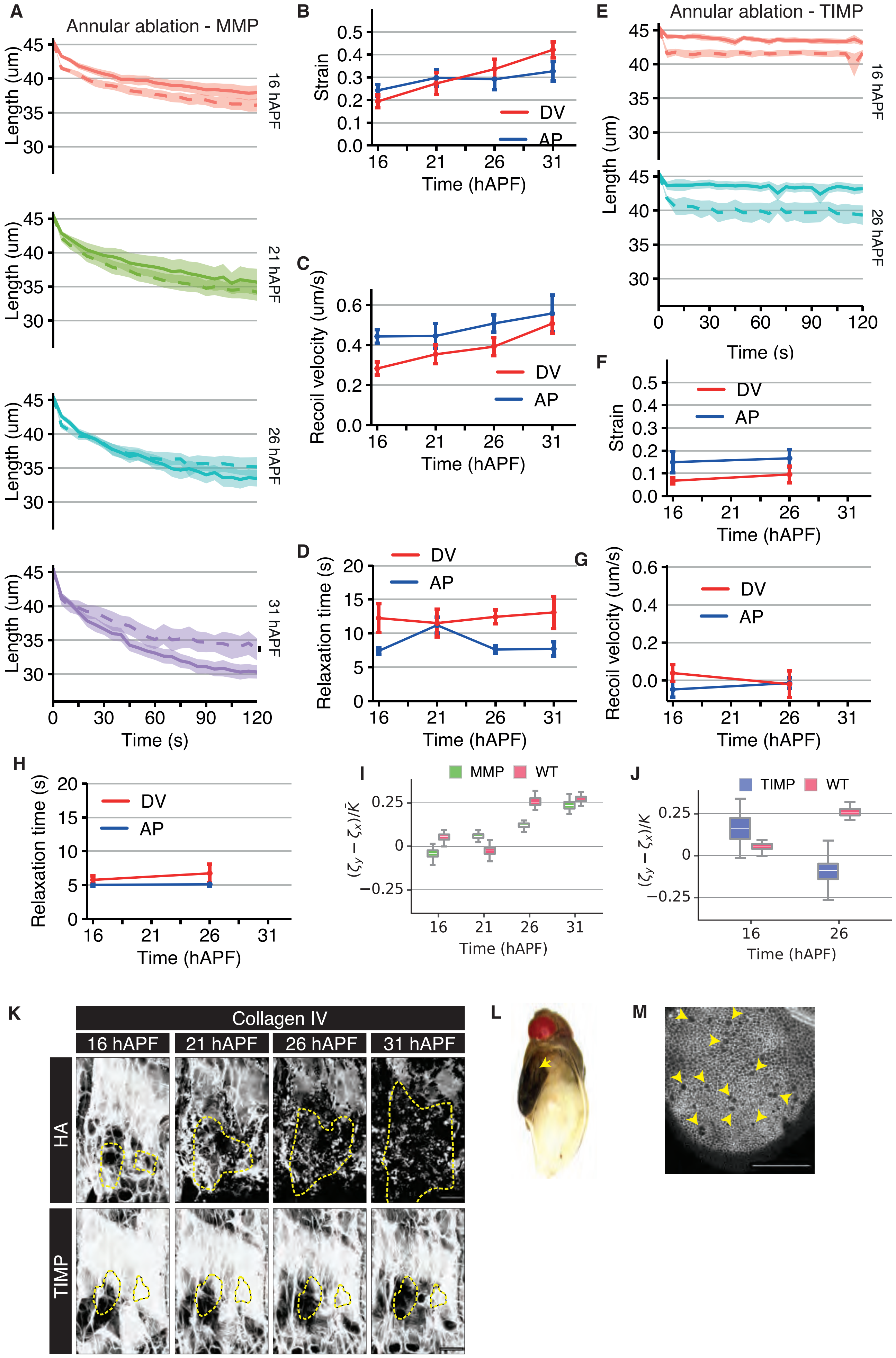
A
Annular ablation - MMP
Length (um)
16 hAPF
21 hAPF
26 hAPF
31 hAPF
Time (s)
B
Strain
DV
AP
Time (hAPF)
C
Recoil velocity (um/s)
D
Relaxation time (s)
E
Annular ablation - TIMP
F
G
H
I
MMP
WT
$(\zeta_y - \zeta_x)/\bar{K}$
J
TIMP
K
Collagen IV
16 hAPF
21 hAPF
26 hAPF
31 hAPF
HA
TIMP
L
M

Figure S7

**Figure S7. Tissue mechanics and histoblast growth are affected in pupae expressing MMP1 or TIMP.**

(A) Length of AP (dashed) and DV (solid) axes in excised cells from MMP1 expressing pupae under the control of *32B-GAL4* after annular ablation.

(B-D) Quantification of Hencky's true strain (B), recoil velocity (C) and relaxation time (D) of excised histoblasts after laser ablation, along the DV and AP axes throughout development, in MMP1 expressing pupae.

(E) Length of AP (dashed) and DV (solid) axes in excised cells from TIMP expressing pupae under the control *32B-GAL4* after annular ablation.

(F-H) Quantification of Hencky's true strain (F), recoil velocity (G) and relaxation time (H) of excised histoblasts after laser ablation, along the DV and AP axes throughout development, in TIMP expressing pupae.

(I) Normalized, anisotropic tension (difference of DV and AP tensions, $\zeta_y$-$\zeta_x$), as a function of time, obtained from fitting model parameters to excised disc deformations in pupae expressing MMP1.

(J) Normalized, anisotropic tension (difference of DV and AP tensions, $\zeta_y$-$\zeta_x$), as a function of time, obtained from fitting model parameters to excised discs deformation in pupae expressing TIMP.

(K) Snapshots of pupae expressing Vkg::GFP (Collagen IV), and overexpressing HA (control) or TIMP under the control of *32B-GAL4* throughout development. Yellow dotted line outlines the histoblasts as seen with E-cad::Tom (not shown). Scale bar = 50 $\mu$m.

(L) Pharate pupa overexpressing TIMP under the control of *32B-GAL4*. Yellow arrows indicate the normal development of the wing.

(M) Still of wing disc from a movie of a pupa expressing E-cad::GFP, and overexpressing TIMP under the control of *32B-GAL4* at the same developmental time as histoblast expansion is inhibited. Note that cell divisions are still occurring (yellow arrowheads). Scale bar = 50 $\mu$m.

| Quantity | Value |
|---|---|
| (a) Starting cell number at 0 hAPF | $18 \pm 2$ |
| (b) Cell number after cleavage divisions at 12 hAPF | $180 \pm 20$ |
| (c) Estimated cell number end of development 40 hAPF | $2900 \pm 200$ |
| (d) Avg. cleavage divisions per cell (up to 12 hAPF) | $3.3 \pm 0.2$ |
| (e) Avg. expansion divisions per cell (14 hAPF to end) | $4.0 \pm 0.2$ |
| (f) Avg. cycle time cleavage (early direct measurements) | $2.7 \pm 0.3$ hr |
| (g) Avg. cycle time cleavage (estimate over 0-12 hAPF) | $3.6 \pm 0.2$ hr |
| (h) Avg. cycle time expansion | $4.5 \pm 0.6$ hr |

**Table S1. Anterior nest growth trajectory key quantities.** Reported uncertainty is standard deviation. (**a**) Cell counts from Figure 3E ($n = 15$). (**b**) Cell counts from Figure 3E ($n = 13$). (**c**) Estimate from simulation. The base case simulation reported in Figure 3 is parameterised to the noborder ROI. Assuming this is a representative subset of the entire anterior nest, the simulation yields an extrapolation for the final cell number in the anterior nest. The reported standard deviation is from $n = 10$ simulation runs, whose variability is significantly smaller than the experimental variability (see Figure 3G). (**d**) Comparing measured cell number at 12 and 0 hAPF and using the formula $N(t)/N_0 = 2^{n_{\text{div.}}}$ where $N$ is cell number and $n_{\text{div.}}$ the average number of divisions per cell yields this estimate. Compare Figure 3F. (**e**) Using the estimated final cell number from part (c) and the measured cell number at 14 hAPF (prior to the rapid onset of expansion divisions) yields this estimate. (**f**) Direct measurements available in the first few hAPF (see Figure 3D) yield this estimate. Three movies were used, each yielding 20-40 cycle time measurements to generate an average for that movie. The value and uncertainty reported here are the average and standard deviation of those per-movie averages. (**g**) Using the formula $N(t)/N_0 = 2^{n_{\text{div.}}}$ as in part (d), the mean divisions per cell $n_{\text{div.}} = t/\mu_{\text{cleavage}}$ where $t$ is the time and $\mu_{\text{cleavage}}$ the mean cycle time of cleavage divisions. Evaluation at $t = 12$ hAPF yields the reported estimate. It is larger than for the direct measurement of part (f). This may indicate that the average cycle time of cleavage divisions really becomes longer, or that some cells do not complete the mean number of cleavage divisions. Either is compatible with the apparent slowing down of cell number increase from 0-12 hAPF which can be seen in Figure 3E. (**h**) The coloured data points in Figure 3D represent measurements of cycle time during the expansion phase from the four WT movies. A simple average of these data points yields the reported value.

| Parameter | Value | Comment |
|---|---|---|
| $N_0$ | 18 | Anterior nest cells at 0 hAPF |
| $[\mathrm{ttd}]_0$ | 1.58 h | Mean time to first division at 0 hAPF |
| $\mu_0$ | 2.67 h | Mean cell cycle time up to 3.3 hAPF |
| $\mathrm{CV}_0$ | 0.24 | Cell cycle time CV up to 3.3 hAPF |
| $\mu_1$ | 4.6 h | Mean cell cycle time between 3.3 and 12.5 hAPF |
| $\mathrm{CV}_1$ | 0.32 | Cell cycle time CV between 3.3 and 12.5 hAPF |
| $t_{\text{pause-on}}$ | 12.5 hAPF | Pause in cell division starts |
| $t_{\text{main}}$ | 14.7 hAPF | Pause in cell division ends |
| $n_{\text{sub.}}$ | 95 | Subset at $t_{\text{main}}$ to be noborder ROI |
| $\mu_2(t)$ | Tracks expt. data | Mean cell cycle time after 14.7h APF |
| $\mathrm{CV}_2$ | 0.2 | Cell cycle time CV after 14.7h APF |
| $\rho_{\text{s}}$ | 0.55 | Correlation coefficient of sister cycle times |
| $s_p$ | 25.7 hAPF | Hill function switch times for arrested cell creation |
| $s_\alpha$ | 24.7 hAPF | Hill function switch times for arrested cell creation |
| $h_p = h_\alpha$ | 34 | Hill function coefficients for arrested cell creation |
| $p_{\text{SOP}}$ | 0.09 | Probability that a new non-arrested cell is an SOP during SOP period |
| $t_{\text{SOP-on}}$ | 20.6 hAPF | SOP period start time |
| $t_{\text{SOP-off}}$ | 24 hAPF | SOP period stop time |

**Table S2.** Table of parameters for the base case of the full simulation (Figures 3E-H). CV: coefficient of variation.

| Parameter | Note | Value/comment |
|---|---|---|
| $P(\tau_{\text{TTD}})$ | From expt. | Time-to-division distribution (frame 0) |
| $\mu(t)$ | Tracks expt. | Mean cycle time |
| $\sigma(t)$ | Tracks expt. | SD cycle time |
| $\mu_{\text{fix}}$ | Fixed mean/CV version | $4.43, 4.74, 4, 4.92$ h (WT1–4) |
| $\text{CV}_{\text{fix}}$ | Fixed mean/CV version | $0.19, 0.2, 0.2, 0.23$ (WT1–4) |
| $s_p$ | Measured in expt. | $26.4, 27.8, 26, 25.4$ hAPF (WT1–4) |
| $s_\alpha$ | Measured in expt. | $25, 27, 25.2, 24.6$ hAPF (WT1–4) |
| $h_p$ | Measured in expt. | $42, 32, 40, 46$ (WT1–4) |
| $h_\alpha$ | Measured in expt. | $42, 38, 40, 36$ (WT1–4) |
| $p_{\text{SOP}}$ | As full simulation, chosen | 0.09 |
| $t_{\text{SOP-on}}, t_{\text{SOP-off}}$ | Chosen | $20.6, 24$ hAPF (all WTs) |

**Table S3.** Parameter information for the short (starts at 16 hAPF) simulation (Figure S3G-J). In contrast to the full simulation, nearly all parameters are directly determined from experiment. An individual simulation set is performed for each WT experimental movie. Times are given for each movie before time-alignment

### ECM REMODELING AND SPATIAL CELL CYCLE COORDINATION DETERMINE TISSUE GROWTH KINETICS - SUPPLEMENTARY THEORY

ANNA P. AINSLIE, JOHN ROBERT DAVIS, JOHN J. WILLIAMSON, ANA FERREIRA, ALEJANDRO TORRES-SÁNCHEZ, ANDREAS HOPPE, FEDERICA MANGIONE, MATTHEW B. SMITH, ENRIQUE MARTIN-BLANCO, GUILLAUME SALBREUX, AND NICOLAS TAPON

#### 1. Analysis of tracked and segmented movies

Each tracked and segmented movie is stored on disk in the form of a consecutive series of folders, one for each frame of the movie, to be passed on to the Tissue Miner analysis package [2, 3]. Each frame-folder contains three images: the original confocal projection image, which is used only as a background on which to plot analysed quantities (e.g., Figure 2F); a tracking mask, comprising a skeleton in which each cell is given a unique colour which persists over time, encoding the tracking; a division mask, comprising a skeleton in which each pair of newly-divided daughter cells in a given frame is highlighted.

Tissue Miner parses these images, converting the information they encode into a series of tables describing the trajectory of the movie: vertex positions in each frame, cell centroid coordinates, cell areas, division events, cell extrusions, and so on. Exhaustive detail is given in Refs. [2, 3], which we will not recapitulate here.

For the purpose of our analyses, three aspects of Tissue Miner are particularly useful. First, the table *cellsDB*, which contains 'dynamic' information on each cell through time – the position and area of a cell in each frame, for example. Second, the table *cellinfoDB* contains meta-information on each cell. For example, whether and when a cell appears by a division event, or is present from the start of the movie, and whether a cell disappears by dividing itself, or by exiting the field-of-view. Thirdly, Tissue Miner contains a number of ready-made scripts which automate common types of analysis. For example, such

*Date*: November 5, 2020.

scripts can be used to obtain the decomposition of tissue deformation into various cellular contributions (see Figure 1).

Tissue Miner's data can be interrogated and manipulated in both the R and Python languages (we have mostly used R), which allows to create customised analyses such as our quantification of arrested cell appearance (Figure 2G).

1.1. **Definition of noborder ROI.** During development, cells both enter and leave the movie field of view. For much of our analysis, it is convenient to work with a population of histoblast cells that are not affected by this. We therefore define the 'noborder' ROI. It contains histoblast cells that are members of lineages of which no member encounters the boundary of the field of view at any point in the movie. In practice, the noborder ROI ends up being a subset of the anterior histoblast nest, since the posterior nest disappears entirely from view during the course of the movie.

1.2. **Isotropic shear decomposition.** Following Ref. [2], we perform a decomposition of the tissue area expansion rate of the noborder ROI, $v = \frac{1}{A}\frac{dA}{dt}$ with $A$ the tissue area, as follows:

$$v = \frac{1}{a}\frac{da}{dt} + k_d - k_e \ , \quad [1]$$

where $a$ is the average cell area, $k_d$ is the cell division rate and $k_e$ the cell extrusion rate. This decomposition can be understood from the simple relation:

$$A = a \times N \ , \quad [2]$$

where $N$ is the number of cells in the tissue. Differentiating this relation gives Eq. [1], following the identification $\frac{1}{N}\frac{dN}{dt} = k_d - k_e$.

Integrating Eq. 1 between $t_0$ and $t$ yields the cumulative shear relation

$$\ln \frac{A(t)}{A_0} = \ln \frac{a(t)}{a_0} + \int_{t_0}^{t} dt k_d - \int_{t_0}^{t} dt k_e \ , \quad [3]$$

with $A_0 = A(t_0)$ and $a_0 = a(t_0)$. In Figures S1L-O different terms of Eq. [1], calculated from finite differences between movie frames, are plotted as a function of time. Summing these contributions yields a cumulative shear decomposition plot (Figures 1C-D) which corresponds to Eq. [3], up to time discretization errors.

1.3. **Cell elongation calculation.** In laser ablation experiments reported in Figures 5 and 7, we obtain the change in cell elongation before and after ablation. We use cell elongation obtained from cell segmentations, as described in [3]. Briefly, the cell centroid $\mathbf{r}_c$ is defined by an integral over all points $\mathbf{r}$ within the cell, which has area $A_c$:

$$\mathbf{r}_c = \frac{1}{A_c} \int \mathbf{r} \, \mathrm{d}A \; . \qquad [4]$$

The components $Q_{xx} = -Q_{yy}$ and $Q_{xy} = Q_{xy}$ of the 2D nematic tensor describing cell elongation are calculated as follows:

$$\begin{aligned} Q_{xx} &= \frac{1}{A_c} \int \cos(2\phi) \, \mathrm{d}A \\ Q_{xy} &= \frac{1}{A_c} \int \sin(2\phi) \, \mathrm{d}A \; , \end{aligned} \qquad [5]$$

in which $\phi$ is the angle between the vector $\mathbf{r} - \mathbf{r}_c$ and the $x$ axis (approximately the dorsoventral axis of the animal in our case). The elongation norm (Figure S2C) is defined as $|Q| = \sqrt{Q_{xx}^2 + Q_{xy}^2}$.

1.4. **Cell-cell distance and neighbour relations.** To define a distance between two cells based on neighbour relations (Figures 2D, 2E, S2D, S2F) we proceed as follows. We define as nearest-neighbours cells which are, at any time point in the lifetime, sharing a common edge. We then obtain second, third,.., nth nearest neighbours iteratively: for instance second-nearest neighbours have a common nearest-neighbour but are not themselves nearest neighbours.

1.5. **Calculation of cell cycle time correlations.** To calculate correlations between cell cycle time of pairs of cells (Figure 2), we proceed as follows. We consider the set of dividing cells of the anterior nest which can be tracked from their birth to their division, $D$, and for every cell $x \in D$ we denote the cell cycle time $\tau_x$. For a given pairing condition (for instance, mother and daughters), we find all cell pairs $(x, y)$ for which the pairing condition is satisfied, and the Pearson correlation coefficient $\rho$ is then calculated as:

$$\rho = \frac{1}{N} \frac{\sum\limits_{(x,y)} (\tau_x - \tau_X)(\tau_y - \tau_Y)}{\sigma_X \sigma_Y} \quad [6]$$

$$\tau_X = \frac{1}{N} \sum_{(x,y)} \tau_x \quad [7]$$

$$\tau_Y = \frac{1}{N} \sum_{(x,y)} \tau_y \quad [8]$$

$$\sigma_X^2 = \frac{1}{N} \sum_{(x,y)} (\tau_x - \tau_X)^2 \quad [9]$$

$$\sigma_Y^2 = \frac{1}{N} \sum_{(x,y)} (\tau_x - \tau_Y)^2 \quad [10]$$

where $N$ is the number of pairs $(x, y)$. In practice we use the scipy.stats.pearsonr function. By construction, $\rho = 0$ if all possible pairs in $D$ are included, and $\rho = 1$ if only identical cells are included (i.e., considering only pairs $(x, x)$ for $x \in D$).

To obtain the Spearman correlation coefficient $\rho_S$ for a given pairing condition, we find all cell pairs $(x, y)$ for which the pairing condition is satisfied. We then directly use the scipy.stats.spearmanr function on this data.

We also calculated correlation coefficients defined such that, for every cell $x$, a cell $y$ is chosen randomly among possible pairs (for instance, when a mother has two daughters, only one daughter is associated to each mother). For different pairing conditions, we formed set of pairs defined in this way a 100 times, and calculated the average correlation coefficients. We found similar results with this method than by taking into account all possible pairs: the Spearman correlation coefficient was $\rho_S = 0.43 \pm 0.03$, $0.33 \pm 0.03$, $0.25 \pm 0.02$, $0.2 \pm 0.02$,$0.08 \pm 0.01$ for cells born within 30 minutes of each other, excluding sisters, and positioned at distance 1, 2, 3, 4, and $> 4$ from each other; $\rho_S = 0.06 \pm 0.03$ for mother-daughters, and $\rho_S = 0.33 \pm 0.04$ for cousins (mean$\pm$standard deviations are obtained from 4 WTs).

To test to what extent correlations in cell cycle time between cousins can be explained by spatial relationships, we generated artificial cell pairs where, for each cell $x$ which can be tracked from birth to division, we allocate two cells $y$ which are neither sister or cousins of $x$, but are chosen according to the following rules:

- With probability $p_i$, one selects the subset of cells $Y$ which are at distance $i$ from $x$, where $p_i$ is the fraction of cousins cells which are at distance $i$ from each other (Figure S2F).
- Within $Y$ and for cells which can be tracked from birth to divisions, a cell $y$ is chosen with a probability that depends on their birth time relative to the birth time of $x$, $|\tau_y - \tau_x|$. This probability is chosen to reflect the distribution of birth time differences of cousins, which is roughly exponentially distributed (Figure S2G).

The correlation coefficient of these artifically constructed pairs is then calculated, for 10 different random realizations (Figure S2H).

#### 2. Simulation of tissue growth

Here we describe simulations conducted to model proliferation of the anterior-nest histoblast cells. The simulations do not attempt to resolve spatial behaviour, rather they aim to model the increase of cell number over time and the timing and intensity of waves of cell division. We first describe general features of the simulations before specifying to two distinct versions we have used. Briefly, we track a time-evolving population of $N(t)$ cells, consisting of $N_d(t)$ dividing cells, $N_a(t)$ arrested cells, and $N_{\mathrm{SOP}}(t)$ non-proliferative sensory organ precursors (SOPs). Each dividing cell is assigned a cell cycle time $\tau$, stochastically selected from a specified distribution $P(\tau)$. At the end of the cell cycle time, a dividing cell divides to give rise to two daughter cells. After division, the type of daughter cells is determined stochastically according to a time-changing probability.

In practice the cell cycle time distribution $P(\tau)$, with mean $\mu$ and standard deviation $\sigma$, is chosen to be bivariate normal,

$$P_{\mathrm{bvn.}}(\tau_1, \tau_2) = \frac{1}{2\pi\sigma^2\sqrt{1-\rho_{\mathrm{s}}^2}} \exp\left(-\frac{z}{2(1-\rho_{\mathrm{s}}^2)}\right), \qquad [11]$$

where

$$z \equiv \frac{(\tau_1-\mu)^2 + (\tau_2-\mu)^2 - 2\rho_{\mathrm{s}}(\tau_1-\mu)(\tau_2-\mu)}{\sigma^2} \qquad [12]$$

and $\rho_{\mathrm{s}}$ is the correlation coefficient, such that correlated cycle times $\tau_1$ and $\tau_2$ are selected in pairs for sister cells (see Figures 2D, S2D). The mean $\mu$ and standard deviation $\sigma$ of the cycle time distribution may vary in time, as described further below. The correlation coefficients between sister cells is set to the measured experimental average between wild-types, $\rho_{\mathrm{s}} = 0.55$ (Figure 2D).

A simulation timestep consists of incrementing each cell's age $a$ and then handling any resulting divisions events. Cells divide when their age exceeds their cycle time $\tau_i$, i.e.

$$\text{cell } i \text{ divides if } a_i > \tau_i \ . \tag{13}$$

The default behaviour is to create a pair of daughter cells that will each divide in turn. In Figure S2I we show the cell division rate for a simple simulation where all cells are dividing (no SOP or arrested cells), cell cycle times of sister cells are taken from the probability distribution $P_{\mathrm{bvn.}}(\tau_1, \tau_2)$, and all cells are at age 0 at the beginning of the simulation.

In simulations shown in other panels, a division event can lead to the generation of zero, one or two arrested cells, with respective probabilities $p_0$, $p_1$, $p_2$, that change in time according to:

$$p_0(t) = 1 - p(t) \tag{14}$$

$$p_1(t) = p(t)(1 - \alpha(t)) \tag{15}$$

$$p_2(t) = p(t)\alpha(t) \tag{16}$$

The time changing variables $p$ and $\alpha$ are set to be Hill-functions:

$$p(t) = \frac{\left(\frac{t}{s_p}\right)^{h_p}}{1 + \left(\frac{t}{s_p}\right)^{h_p}}, \quad \alpha(t) = \frac{\left(\frac{t}{s_\alpha}\right)^{h_\alpha}}{1 + \left(\frac{t}{s_\alpha}\right)^{h_\alpha}} \tag{17}$$

with switch times $s_p$, $s_\alpha$ and Hill coefficients $h_p$, $h_\alpha$ fitted to experimental data as in Figures 2J, K and reported in Table S2. Before 14.7 hAPF the probabilities $p$ and $\alpha$ are set to 0.

Creation of SOP cells is handled simply by setting a window of simulation time $t$ during which there is a fixed probability that a newly-born cell which is not arrested becomes a

SOP:

$$\text{SOP probability } = \begin{cases} p_{\text{SOP}} & \text{if } t_{\text{SOP-on}} < t < t_{\text{SOP-off}} \\ 0 & \text{otherwise} \end{cases} \quad [18]$$

In the experiments, "SOP" cells, once specified, continue to proliferate but do so in a specialised manner: they undergo one in-plane division followed by one or two out-of-plane divisions.. Therefore, for the purpose of our analysis and simulation, they are treated as non-dividing cells once they have been created. The fitted simulation parameter $p_{\text{SOP}}$ encodes the probability per newly-created cell that the cell is an SOP, conditioned to the cell not being arrested.

We will next describe both the 'full' simulation which begins at 0 hAPF and a restricted, modified 'short' version which begins at 16 hAPF. Where relevant we will give parameters for the base case simulation. Parameters are either determined by experimental data, or are free parameters that are chosen to produce simulations results which match experimental data visually. Modifications from the base case described below, are described in the captions of Figures 3 and S3.

2.1. **Full simulation.** The full simulation begins at 0 hAPF and connects the available observations pre-16 hAPF to those in the main movies post-16 hAPF. Here for simplicity we neglect the possibility of cell delamination. We separate the simulation into two phases.

2.1.1. *First phase: from 0hAPF to 14.7hAPF.* The simulation begins with $N_0 = 18$ cells at 0 hAPF (average number of cells measured in the anterior nest at this time, Figure 3E). In the first phase, we can compare simulation results to measurements of anterior nest cell number, taken from fixed animals at this stage (Figure 3E).

During this phase, cell cycle times could be measured in live movies taken in the first $\sim 3$ hAPF (Figure 3D). Therefore the distribution of cycle times in the first $\sim 3$ hAPF of simulations is set to match available cell cycle measurements taken at this stage: $\mu_0 = 2.67$ h, and the coefficient of variation $\text{CV}_0 \equiv \sigma_0/\mu_0 = 0.24$, where $\sigma_0$ is the cycle time standard deviation (determined parameters from experimental data in Figure 3D). The mean time-to-division from 0 hAPF for the first set of divisions was measured to be on average $\sim 1.1$

h shorter than $\mu_0$. We mimic this in simulation by shifting the first round of divisions forward by $\sim 1.1$ h, so that the mean time-to-division from 0 hAPF is $[ttd]_0 = 1.58$ h.

For times $t_{\text{trans}} = 3.3\text{hAPF} \lesssim t \leq 14.7$ hAPF, no experimentally-measured cycle times are available. Measurements of anterior nest cell numbers (Figure 3E) suggest a pause, or at least a dramatic reduction, in cell divisions at around 12 hAPF. We therefore introduce a pause in divisions in the simulation at $t_{\text{pause-on}} = 12.5$ h (chosen parameter).

In between 3.3hAPF and 12.5hAPF, we chose for simplicity a constant mean cell cycle time. Keeping the early, $\leq$ 3.3hAPF value of mean cell cycle time $\mu_0$ does not account for the measured number of cells in the anterior nest (Figure S3A). Instead we therefore set the mean cell cycle time to $\mu_1 = 4.6$h, chosen to roughly match the measured cycle times at the start of the second phase (see Figure 3D). This number leads to predicted increase in cell numbers in agreement with experiments (see Figure 3E and compare with Figure S3A). The coefficient of variation in this stage is also a free parameter, which we chose to be $\text{CV}_1 = \sigma_1/\mu_1 = 0.32$, large enough to avoid oscillatory peaks stronger than experimentally measured after 16hAPF (chosen parameter, see Figure S3D). As illustrated in Figure S3B and the associated caption, the pair of parameters $\mu_1$ and $t_{\text{trans}}$ are reasonably well constrained considering that no cycle time measurements are available for direct comparison. Indeed, $t_{\text{trans}}$ cannot decrease by $\sim 1$ h because it would contradict the early cycle time measurements. On the other hand, increasing $t_{\text{trans}}$ by $\sim 1$ h would require to increase the cell cycle time $\mu_1$ to compensate for the excess number of cells created, and we find that the cell number measurement is then not matched as well as with our choice of parameters (Figure S3B). Of course, we cannot rule out that the cycle time during the phase $3.3\text{hAPF} \leq t \leq 14.7$ hAPF changes in some more complex manner, but the simple choices made here represent a parsimonious explanation for the measured data.

In between 12.5hAPF and 14.7hAPF, a sharp increase in the number of cells occurs (Figure 3E), which we attribute to a burst in cell division. Here we assume that the preceding pause in anterior nest cell number increase is due to inhibition of cell division, while cell ages are still increasing. Therefore in simulations when the pause is released there is a sudden increase in cell number as a sub-population of cells that have exceeded their cycle time during the pause all divide at once. The population of cells that experiences this sudden division is effectively 're-synchronised' in terms of birth times, which has a sharpening effect on the division rate peaks in the main phase (compare Figure S3C in which a different implementation of the pause, which does not result in such re-synchronisation, was used). Therefore, as well as matching the cell number measurements of Figure 3E, another consideration when choosing the pause period was to attain comparable peak sharpness as shown in Figure 3H (see Figure S3E for the outcome with a delay in the pause). The only other free parameter with an effect on this sharpness, $\mathrm{CV}_1$, has a far weaker role, because it only affects the decorrelation of division times prior to the subsequent re-synchronisation caused by the pause period.

Although we do not have live experimental data in between $\sim$ 3.3hAPF and 14.7hAPF, it seems reasonable to describe the pause in cell number increase occuring around 12hAPF (Figure 3E) as the end of the 'cleavage' divisions characteristic of the early stages, and the subsequent cell number increase around $14-16$hAPF as the beginning of the 'expansion' divisions characteristic of the main experimental window.

2.1.2. *Second phase: from 14.7hAPF to 40hAPF.* At 14.7hAPF, a subset of $n_{\text{sub.}} = 95$ cells is selected to represent the noborder ROI of the movies. The mean cycle time $\mu_2(t)$ is hereafter set to track the experimental data as shown in Figure 3D (determined parameter, a 3rd-order polynomial fit to the data up to 28 hAPF and a constant value afterwards), while the coefficient of variation is fixed at $\mathrm{CV}_2 = 0.2$ (determined parameter), a representative CV over all time and over all movies. The Hill functions for $p(t)$ and $\alpha(t)$ have switch times of $s_\alpha = 24.7$ hAPF, $s_p = 25.7$ hAPF and Hill coefficients $h_p = h_\alpha = 34$, found by manual fitting of the collected time-aligned experimental data in Figures 2J, K.

During this phase, SOPs are generated with a probability of $p_{\text{SOP}} = 0.09$ per new cell between $t_{\text{SOP-on}} = 20.6$ and $t_{\text{SOP-off}} = 24$ hAPF, conditioned on the cell not being arrested (chosen parameters).

The full simulation is not compared on a movie-by-movie basis to the post-16 hAPF experiments. Rather we create a ribbon from the mean and SD of the experimental cell number data, for the comparison in Figure 3G, and aim for a simulation which goes roughly through the middle of the ribbon. Creating the experimental ribbon is subtle, because of arbitrary differences in the size of the noborder ROI across experiments (the noborder ROI is not biologically meaningful, as it simply comprises lineages not affected by contact with the boundary of the field-of-view). We normalise each movie's noborder cell number as $N_{\mathrm{dl}}(t) = N(t)/N_f$, where $N(t)$ is the total number of cells and $N_f$ is the cell number of that movie evaluated in the 'first common frame' – the earliest frame that exists in all movies once the movies have been time-aligned. The dimensionless cell number $N_{\mathrm{dl}}(t)$ is used to calculate a time-dependent mean $\bar{N}_{\mathrm{dl}}(t)$ and associated SD across the movies. This mean and SD are finally remultiplied by $\bar{N}_f$, the average of $N_f$ across movies, to give a representative ribbon of cell number as shown in Figure 3G. The SOP cell number and arrested cell number are treated in the same way.

We perform 10 runs of the simulation with different random seeds. These lead to the simulation mean and SD shown in Figures 3E and 3G. The simulation division rate in Figure 3H is one representative run, showing that the amplitude and period of oscillations in cell division rate are comparable to experiment.

2.2. **Short simulation.** As described above, the full simulation has some freedom in parameters, principally due to the phase between $\sim$ 3.3hAPF and 14.7hAPF in which no measurements of cell cycle time, and only limited measurements of cell number, are available from experiment. We therefore performed another type of simulation focused only on the post-16 hAPF main movie window, aiming to have parameters tightly constrained by experimental data. We aim to match closely the cell number time series of each movie, and the sharpness of the peaks in the division rate. The short simulations are specialised to each WT movie. Time-alignment of movies relative to each other is irrelevant in this context so is not performed.

The starting condition for the simulation is the initial time-to-division (TTD) distribution $P(\tau_{\mathrm{TTD}})$ measured in frame 0 of the corresponding experimental movie. The TTD is defined, for each cell in the noborder ROI in the first frame, by the time until that cell divides. After the first round of cell divisions, subsequent sister cycle times $\tau$ are selected from a bivariate normal distribution $P_{\mathrm{bvn.}}(\tau_1, \tau_2)$ as in Eq. [11], in which the parameters $\mu$ and $\sigma$ track the time-dependent values from the corresponding movie (determined parameters).

The probability Hill functions $p$ and $\alpha$ for arrested cell creation are set to match those fitted individually to each movie (see Figures 2J and 2K) (determined parameters, Table S3). Creation of the small number of SOPs is handled similarly to the full simulation (chosen parameters, Table S3).

Unlike the full simulations, the short simulations also account for the small amount of histoblast delamination, since we aim to closely match the experimental cell number in each WT movie. In each simulation timestep, the number of delaminations occurring in the corresponding frame of the corresponding movie is read-in. The required number of dividing cells are selected at random and deleted from the simulation.

Like the full simulation, 10 runs of the short simulations with different random seeds are performed.

Thus, unlike the full simulation, the short simulations have essentially no free or chosen parameters (excepting the SOP creation probabilities, which bear very little on the overall behaviour because so few SOPs are created).

The close match to experimental data for cell number (Figure S3H) gives confidence that the short simulation captures the important features of histoblast population growth, with minimal free parameters. The division rate matches experiments well overall (Figure S3I), though in WT2 and WT4 the absolute value of the Fourier transform exhibits on average a peak of smaller magnitude than the experiment. We note that in simulations in which the mean and CV were fixed rather than tracking experimental data, the peak in the absolute value of the Fourier transform was less pronounced or lost (Figure S3J), indicating that the experimentally measured variation in cell cycle time (Figure 3D) tends to favour oscillations in the cell division rate.

#### 3. Active elastic deformation model for annular ablations

In this section we describe an active elastic model for the tissue deformation following laser ablation. The tissue is described as an elastic material, connected to an external material by elastic links, and subjected to active anisotropic tension. We solve the model numerically using a finite element method and compare solutions to experimentally measured deformations. This is done by segmenting cell centers and cell shapes before and after annular ablation to obtain the deformation field of the cut circular piece of tissue, and comparing it to model predictions. We also take into account the deformation of the outer boundary of the cut, measured along two orthogonal axis.

3.1. **Elastic model for the constriction of the excised disc.** We consider a 2D linear-elastic material, with shear and bulk moduli $K$ and $\bar{K}$ respectively, subjected to a homogeneous, possibly anisotropic, active tension and adhering to a substrate with elastic bonds of elasticity $k$. We denote $\mathbf{u}$ the displacement field, $t_{ij}$ the tension tensor with latin indices referring to $x, y$ cartesian coordinates. The force balance then reads

$$\partial_i t_{ij} - k u_j = 0 \ , \qquad [19]$$

and the constitutive equation for the total tension tensor reads

$$t_{ij} = 2K\tilde{u}_{ij} + \bar{K} u_{kk}\delta_{ij} + t_{ij}^{\mathrm{act}}, \qquad [20]$$

with $t_{ij}^{\mathrm{act}}$ the active tension tensor, $u_{ij} = \frac{1}{2}(\partial_j u_i + \partial_i u_j)$ is the strain tensor, $\tilde{u}_{ij} = \left(u_{ij} - \frac{1}{2}u_{kk}\delta_{ij}\right)$ its anisotropic shear component, and $u_{kk}$ the isotropic shear. We consider here $t_{ij}^{\mathrm{act}} = \zeta_x \delta_{ix}\delta_{jx} + \zeta_y \delta_{iy}\delta_{jy}$ with $\zeta_x$ and $\zeta_y$ the principal components of the active tension. We assume that the tissue tension after laser ablation is the sum of the pre-existing uniform tension tensor $t_{ij}^{\mathrm{act}}$ (identified here with the active tension, in an active elastic description of the material) and additional elastic stresses arising from deformation following laser ablation.

Combining the force balance equation [19] and the constitutive equation [20] results in the following PDE in terms of the components of the displacement $u_i(x, y)$:

$$\partial_i \left[2K\tilde{u}_{ij} + \bar{K} u_{kk}\delta_{ij} + t_{ij}^{\mathrm{act}}\right] - k u_j = 0, \qquad [21]$$

For a patch of material, Eq. [21] is supplemented with the traction-free boundary condition

$$t_{ij} n_i = \left[2K\tilde{u}_{ij} + \bar{K} u_{kk}\delta_{ij} + t_{ij}^{\mathrm{act}}\right] n_i = 0, \quad \text{at } \partial\Omega, \qquad [22]$$

where $n_i$ is the outer normal to the domain $\Omega$ at the boundary $\partial\Omega$.

Before proceeding further we note that if the active tension is uniform ($t_{ij} = \zeta\delta_{ij}$) and the elastic external resistance vanishes ($k = 0$), the force balance equation [21] together with the boundary condition [22] is solved in polar coordinates $(r, \theta)$ by the radial displacement field:

$$u_r = -\frac{\zeta}{2\bar{K}}r \qquad [23]$$

corresponding to a uniform isotropic shear, $u_{kk} = \partial_r u_r + \frac{u_r}{r} = -\frac{\zeta}{\bar{K}}$. As excised discs area contraction occurs non-uniformly at early developmental times (see deformation field at 16hAPF in Figure 5C), we conclude that these simplifying hypothesis do not apply in practice.

3.2. **Numerical resolution of the equations using a finite element method.** To solve Eq. [21]-[22] numerically we employ a finite element method (FEM). In the finite element method, the weak form of Eq. [21] is obtained by multiplying Eq. [21] by an arbitrary test function $w(x, y)$ and integrating over $\Omega$

$$\begin{aligned} 0 &= \int_\Omega \left\{\partial_i \left[2K\left(u_{ij} - \frac{1}{2}u_{kk}\delta_{ij}\right) + \bar{K}u_{kk}\delta_{ij} + t_{ij}^{\text{act}}\right] - ku_i\right\} w dxdy \\ &= \int_\Omega \left\{-(\partial_i w)\left[2K\left(u_{ij} - \frac{1}{2}u_{kk}\delta_{ij}\right) + \bar{K}u_{kk}\delta_{ij} + t_{ij}^{\text{act}}\right] - ku_i w\right\} dxdy, \end{aligned} \qquad [24]$$

where to get to the last line we have integrated by parts the term in the square brackets, and used the divergence theorem and Eq. [22] to eliminate the term in the boundary. This is the weak form of Eqs. [21] and [22]. We now consider a discretisation of our domain $\Omega$ into $n_t$ triangles $\Omega_t$ with $n$ vertices. We discretise the displacement as

$$u_i^h(x, y) = \sum_{a=1}^{n_v} u_i^a N^a(x, y), \qquad [25]$$

where the superscript $^h$ indicates that the solution is approximate, $a$ is an index denoting the label of a vertex in the discretisation and $N^a(x, y)$ is its basis function; here we employ linear interpolants, also known as tent functions. In a Galerkin FEM, the set of basis functions $N^a(x, y)$ is also employed as a set of weight functions. The discrete version of Eq. [24] becomes

$$\sum_{b=1}^{n}\int_{\Omega}\left[-\partial_i N^a\left(K\left(u_j^b\partial_i N^b+u_i^b\partial_j N^b-u_k^b\delta_{ij}\partial_k N^b\right)\right.\right.$$
$$\left.\left.+\bar{K}u_k^b\delta_{ij}\partial_k N^b\right)-kN^aN^bu_i^b\right]dxdy=\int_{\Omega}t_{ij}^{\mathrm{act}}\partial_i N^a dxdy$$
$$\sum_{b=1}^{n}\left[\int_{\Omega}\left[\partial_i N^a\left(K\left(\delta_{jk}\partial_i N^b+\delta_{ik}\partial_j N^b-\delta_{ij}\partial_k N^b\right)+\bar{K}\delta_{ij}\partial_k N^b\right)\right.\right.$$
$$\left.\left.+kN^aN^b\delta_{ik}\right]dxdy\right]u_k^b=-\int_{\Omega}t_{ij}^{\mathrm{act}}\partial_i N^a dxdy, \quad [26]$$

which we can write as a linear system of equations

$$\left(K\mathsf{A}^K+\bar{K}\mathsf{A}^{\bar{K}}+k\mathsf{A}^k\right)\mathsf{u}=\zeta_x\mathsf{B}^x+\zeta_y\mathsf{B}^y. \quad [27]$$

Here

$$\mathsf{u}=\begin{pmatrix}u_x^1\\u_y^1\\\vdots\\u_x^n\\u_y^n\end{pmatrix}, \quad [28]$$

is the vector of unknowns,

$$\mathsf{B}^x=\begin{pmatrix}-\int_{\Omega}\partial_x N^1 dxdy\\0\\\vdots\\-\int_{\Omega}\partial_x N^n dxdy\\0\end{pmatrix},\qquad \mathsf{B}^y=\begin{pmatrix}0\\-\int_{\Omega}\partial_y N^1 dxdy\\\\\vdots\\0\\-\int_{\Omega}\partial_y N^n dxdy\end{pmatrix}, \quad [29]$$

are the vectors in the right-hand side, and the matrices have the form

$$\mathsf{A}^X = \begin{pmatrix} A^X_{1,x,1,x} & A^X_{1,x,1,y} & A^X_{1,x,2,x} & A^X_{1,x,2,y} & \cdots & A^X_{1,x,n,x} & A^X_{1,x,n,y} \\ A^X_{1,y,1,x} & A^X_{1,y,1,y} & A^X_{1,y,2,x} & A^X_{1,y,2,y} & \cdots & A^X_{1,y,n,x} & A^X_{1,y,n,y} \\ A^X_{2,x,1,x} & A^X_{2,x,1,y} & A^X_{2,x,2,x} & A^X_{2,x,2,y} & \cdots & A^X_{2,x,n,x} & A^X_{2,x,n,y} \\ A^X_{2,y,1,x} & A^X_{2,y,1,y} & A^X_{2,y,2,x} & A^X_{2,y,2,y} & \cdots & A^X_{2,y,n,x} & A^X_{2,y,n,y} \\ \vdots & \vdots & \vdots & \vdots & \ddots & \vdots & \vdots \\ A^X_{n,x,1,x} & A^X_{n,x,1,y} & A^X_{n,x,2,x} & A^X_{n,x,2,y} & \cdots & A^X_{n,x,n,x} & A^X_{n,x,n,y} \\ A^X_{n,y,1,x} & A^X_{n,y,1,y} & A^X_{n,y,2,x} & A^X_{n,y,2,y} & \cdots & A^X_{n,y,n,x} & A^X_{n,y,n,y} \end{pmatrix}, \quad [30]$$

for $X = K, \bar{K}, k$, with components

$$\begin{aligned} \mathsf{A}^K_{a,j,b,k} &= \int_\Omega \left[ (\partial_i N^a) \left(\partial_i N^b\right) \delta_{jk} + (\partial_k N^a) \left(\partial_j N^b\right) - (\partial_j N^a) \left(\partial_k N^b\right) \right] dxdy, \\ \mathsf{A}^{\bar{K}}_{a,j,b,k} &= \int_\Omega (\partial_j N^a) \left(\partial_k N^b\right) dxdy, \\ \mathsf{A}^k_{a,j,b,k} &= \int_\Omega N^a N^b dxdy. \end{aligned} \quad [31]$$

To compute these integrals, we partition the domain $\Omega$ into the triangles $\Omega_t$ and use Gaussian integration to approximate the integral in each $\Omega_t$,

$$\int_\Omega f(x,y)dxdy = \sum_{t=1}^{n_t} \int_{\Omega_t} f(x,y)dxdy = \sum_{t=1}^{n_t} \sum_{g=1}^{n_g} w_g f(x_g^t, y_g^t) J^t \quad [32]$$

where $w_g$ are the Gaussian integration weights, $(x_g^t, y_g^t)$ the coordinates of the Gauss points in triangle $t$, and $J^t$ the triangle area.

3.3. **Least-squares procedure for parameter fitting from annular ablation experiments.** From annular ablation experiments, we can extract (1) cell-centre displacements $u_i(x_c, y_c)$, (2) relative cell area change $\lambda(x_c, y_c)$ and (3) cell elongation change in the form of a nematic tensor $Q_{ij}(x_c, y_c)$, where $(x_c, y_c)$ represent the cell centre right before ablation. For the cell centre displacements, we remove the center-of-mass motion of the inner disc. We would then like to use the FEM solution to fit the parameters $\boldsymbol{p} = (K/\bar{K}, \zeta_x/\bar{K}, \zeta_y/\bar{K}, k/\bar{K})$; note that these parameters are normalised by $\bar{K}$ and form a set of independent parameters in Eq. [21]. We first define the function

$$\begin{aligned}\mathcal{L}(\boldsymbol{p}; u, \lambda, Q_{ij}) = \frac{1}{2}\sum_c \Big[ & k_u \left|u_i^{cc}(x_c, y_c) - u_i^h(x_c, y_c; \boldsymbol{p})\right|^2 \\ & + k_\lambda \left(\lambda(x_c, y_c) - \partial_i u_i^h((x_c, y_c; \boldsymbol{p})\right)^2 \\ & + k_Q \left|Q_{ij}(x_c, y_c) - \tilde{u}_{ij}^h(x_c, y_c; \boldsymbol{p})\right|^2 \Big]\end{aligned} \qquad [33]$$

where $k_u$, $k_\lambda$ and $k_Q$ are penalty parameters (for the fits in this article we have used $k_u = 1/R^2$, $k_\lambda = k_Q = 1$ although we checked that the fits are relatively insensitive to variations of the relative weight of the displacement and shear penalty parameters within a range $k_u/k_\lambda = 0 \leftrightarrow 20/R^2$). This function penalises deviations of the finite element solution $u^h(x, y)$ obtained for the set of parameters $\boldsymbol{p}$ from experimental data. To fit experimental data and obtain the model parameters $\boldsymbol{p}$, we then minimise

$$\boldsymbol{p} = \underset{\text{arg min}}{\boldsymbol{p}'} \; \mathcal{L}(\boldsymbol{p}'; u, \lambda, Q_{ij}), \qquad [34]$$

for which we use a L-BFGS algorithm as implemented in the function `minimize` of the python `Scipy` package. We compute gradients of $\mathcal{L}$ as

$$\begin{aligned}\frac{\partial \mathcal{L}}{\partial \boldsymbol{p}} = \sum_c \Big[ & k_u \left(u_i^{cc}(x_c, y_c) - u_i^h(x_c, y_c; \boldsymbol{p})\right) \frac{\partial u_i^h}{\partial \boldsymbol{p}} \\ & + k_\lambda \left(\lambda(x_c, y_c) - \partial_i u_i^h((x_c, y_c; \boldsymbol{p})\right) \frac{\partial(\partial_i u_i^h)}{\partial \boldsymbol{p}} \\ & + k_Q \left(Q_{ij}(x_c, y_c) - \tilde{u}_{ij}^h(x_c, y_c; \boldsymbol{p})\right) \frac{\partial \tilde{u}_{ij}^h}{\partial \boldsymbol{p}} \Big].\end{aligned} \qquad [35]$$

To compute $\frac{\partial u_i^h}{\partial \boldsymbol{p}}, \frac{\partial(\partial_i u_i^h)}{\partial \boldsymbol{p}}, \frac{\partial \tilde{u}_{ij}^h}{\partial \boldsymbol{p}}$, we note that

$$\begin{aligned}&\frac{\partial u_i^h}{\partial \boldsymbol{p}} = \sum_{a=1}^{n_v} N^a \frac{\partial u_i^a}{\partial \boldsymbol{p}}, \qquad \frac{\partial\left(\partial_i u_i^h\right)}{\partial \boldsymbol{p}} = \sum_{a=1}^{n_v} (\partial_i N^a) \frac{\partial u_i^a}{\partial \boldsymbol{p}}, \\ &\frac{\partial \tilde{u}_{ij}^h}{\partial \boldsymbol{p}} = \frac{1}{2}\sum_{a=1}^{n_v} \left[\partial_j N^a \delta_{ik} + \partial_i N^a \delta_{jk} - \partial_k N^a \delta_{ij}\right] \frac{\partial u_k^a}{\partial \boldsymbol{p}},\end{aligned} \qquad [36]$$

and $\frac{\partial u_i^a}{\partial \boldsymbol{p}}$ are the components of $\frac{\partial \mathsf{u}}{\partial \boldsymbol{p}}$. To calculate these variations, we note that for instance varying $K$ by $\delta K$, the solution $\mathsf{u} + \delta \mathsf{u}$ satisfies

$$\left((K + \delta K)\mathsf{A}^K + \bar{K}\mathsf{A}^{\bar{K}} + k\mathsf{A}^k\right)(\mathsf{u} + \delta \mathsf{u}) = \zeta_x \mathsf{B}^x + \zeta_y \mathsf{B}^y, \qquad [37]$$

and since $\mathsf{u}$ is a solution of the original problem, we have up to first order terms

$$\left(K\mathsf{A}^K + \bar{K}\mathsf{A}^{\bar{K}} + k\mathsf{A}^k\right)\delta\mathsf{u} = -\delta K\mathsf{A}^K\mathsf{u}, \tag{38}$$

so

$$\left(K\mathsf{A}^K + \bar{K}\mathsf{A}^{\bar{K}} + k\mathsf{A}^k\right)\frac{\partial\mathsf{u}}{\partial K} = -\mathsf{A}^K\mathsf{u}. \tag{39}$$

Equivalently, one finds that

$$\begin{aligned}\left(K\mathsf{A}^K + \bar{K}\mathsf{A}^{\bar{K}} + k\mathsf{A}^k\right)\frac{\partial\mathsf{u}}{\partial k} &= -\mathsf{A}^k\mathsf{u},\\ \left(K\mathsf{A}^K + \bar{K}\mathsf{A}^{\bar{K}} + k\mathsf{A}^k\right)\frac{\partial\mathsf{u}}{\partial\zeta_x} &= \mathsf{B}^x,\\ \left(K\mathsf{A}^K + \bar{K}\mathsf{A}^{\bar{K}} + k\mathsf{A}^k\right)\frac{\partial\mathsf{u}}{\partial\zeta_y} &= \mathsf{B}^y.\end{aligned} \tag{40}$$

Note that variations with respect to the normalised parameters $(K/\bar{K}, \zeta_x/\bar{K}, \zeta_y/\bar{K}, k/\bar{K})$ can be directly obtained from the variations with respect to the non-normalised parameters $(K, \zeta_x, \zeta_y)$, e.g. $\frac{\partial\mathsf{u}}{\partial(K/\bar{K})} = \bar{K}\frac{\partial\mathsf{u}}{\partial K}$. In practice, we perform a single fit including all times and mutants and minimise the function $\mathcal{L} = \sum_t\sum_M \mathcal{L}_t^{\mathrm{M}}$ where $t$ =16 hAPF, 21 hAPF, 26 hAPF, 31 hAPF, and M = WT, MMP and TIMP, with the restriction that $K/\bar{K}$ is constant over time and across mutants, and data for TIMP are only for $t$=16 and 26 hAPF.

Together with cell centre displacements, relative cell area change and cell elongation change in the inner patch obtained from our segmented movies, we also included the displacement of the X and Y axes of the outer ablation contour, which are treated as a cell centre displacement in the data (but without an associated changes in cell area and cell elongation). To get a measure of variability of the fit, we performed a bootstrapping analysis: once we got the best fit for the data, we used the list of residuals (the errors in displacement, relative cell area changes and cell elongation change for each cell) to produce new, synthetic samples by assigning to each cell the displacement, relative cell area change and cell elongation change predicted by the fit plus one of the residuals chosen from the list of residuals with replacement. The fit was then repeated for each of these synthetic samples to obtain a distribution for the fitting parameters, which we report in box plots for the different mutants in Figures 5F-H, S5D-E, 7F-I, S7I-J. We produced 100 of these synthetic samples.

We show simulations results using the best fit parameters for the different mutants in Figures 5E and 7D-E. Results from experiments (top) and simulations (bottom) are shown on the computational mesh for comparison. We use a colormap to represent the isotropic shear $u_{ii}$, with linear interpolation on each triangle of the mesh. We use lines to represent the anisotropic shear $\tilde{u}_{ij}$, plotted at each node of the mesh. To plot these lines, we note that, since it is a nematic tensor, the anisotropic shear $\tilde{u}_{ij}$ can be written as $S(n_i n_j - \delta_{ij}/2)$ for some $S$ and $\boldsymbol{n}$, which represent the magnitude ($S$) and orientation ($\boldsymbol{n}$) of the shear. We then find $S$ and $\boldsymbol{n}$ and plot the vector $S\boldsymbol{n}$ as a line. To obtain the experimental values for isotropic and anisotropic shear in the computational mesh, we interpolate the results in the nodes of the mesh using Gaussian kernels centred at the nodes with a width of $R = 3\mu m$, which is also used as a cutoff; each experimental point then contributes with a Gaussian weight to mesh nodes located within a radius $R$ and these weights are normalised to add up to 1 for each node.
